# Morphomechanically-Informed Spatial Curvature Sequencing in Prostate Cancer

**DOI:** 10.64898/2026.07.22.740088

**Authors:** Ivan Kordic, Felix G. Rivera Moctezuma, Hoseyn A. Amiri, Alan Liu, Shuangyi Cai, Mehdia Nadeem Rajab Ali, Lourdes Brea, Abhijeet Venkataraman, Hongshun Shi, Lara Harik, Todd A Sulchek, Jindan Yu, Ahmet F. Coskun

## Abstract

Morphological changes in prostate glands, assessed by Gleason grading, remain the gold standard for diagnosing prostate cancer, yet molecular biomarkers associated with gland shape are not well understood. Here, we introduce CurvSeq, a mechanomorphology-informed framework for spatial sequencing data, and CurvSee, its complementary version for proteomic and imaging datasets. These methods integrate gland boundary curvature, pocket architecture, microenvironmental composition, and molecular profiles to study morphomechanical relationships in prostate adenocarcinoma. Using five independent spatial transcriptomic and multiplexed imaging datasets, we segmented individual prostate glands, extracted gland contours, quantified local curvature and pocket-like concavities, and projected these features onto spatially resolved gene and protein measurements. In Xenium data, CurvSeq distinguished benign and GG1 glands, identifying cancer-associated genes such as PCA3 and AMACR in GG1 glands and basal, basement membrane, and mechanotransduction-associated programs in benign glands. In Visium data, a diffusion-based morphomechanical score ordered benign glands by area, circularity, pocket number, smooth muscle abundance, immune-cell proximity, and remodeling-associated genes including MMP7. In GG4 glands, CurvSeq identified neuroendocrine-like boundary regions associated with MMP7 expression, COL1A1-rich adjacent stroma, and immune-cell accumulation. Finally, CurvSee extended this framework to multiplexed protein imaging, where combined morphology and protein-expression features distinguished Gleason-associated gland states. Together, CurvSeq and CurvSee provide a quantitative framework for linking gland architecture, local microenvironment, and molecular state, showing that prostate gland morphology can be integrated with spatial omics to identify morphomechanical niches associated with cancer progression.

## Introduction

According to the American Cancer Society, nearly 300,000 new cases of Prostate Cancer (PC) were estimated in the United States in 2024. While the number of deaths has been declining over the years, Prostate Cancer (PC) continues to be the second most common cause of cancer-related mortality among men in the US, as an estimated 1 in 44 men is expected to die from the disease ^1^. The most common form of prostate cancer, making up over 90% of cases, is adenocarcinoma, which originates from the epithelial cells that line the prostate gland.

Prostate cancer develops through the uncontrolled proliferation of secretory luminal cells, which produce seminal fluid□^2^. To assess how aggressive and advanced the disease is, pathologists use the Gleason scoring system, which is the current gold standard for prostate cancer grading. This system evaluates the architecture of prostate glands and assigns a score based on the two most common glandular patterns, with values ranging from 2 to 10□^3^. Benign prostate glands typically retain an organized luminal architecture surrounded by a continuous basal-cell layer, whereas invasive adenocarcinoma is characterized by loss of basal cells and disruption of normal glandular organization. Precursor lesions, including prostatic intraepithelial neoplasia and atypical intraductal proliferation, may preserve a partial basal-cell layer despite showing epithelial atypia and increased architectural complexity. These histological distinctions are reflected in Gleason grading, which considers features such as gland fusion, cribriform growth, and loss of luminal organization. In most patients, the disease remains localized, and treatment typically involves prostatectomy or radiation therapy. In cases where the cancer has spread, androgen deprivation therapy (ADT) is the standard approach□^4^. Symptoms and health-related quality-of-life effects vary with disease stage and treatment and can include urinary obstruction or incontinence, pain, fatigue, sexual dysfunction, and hormonal symptoms □^5^.

Prostate cancer is characterized by a high degree of heterogeneity, presenting significant challenges for accurate diagnosis, prognosis, and research ^6^. Genomic methodologies have facilitated the identification of consistent gene expression patterns that correlate with specific histological or clinical features ^7^. Furthermore, prostate morphology, as indicated by Gleason grading, has been demonstrated to align with underlying biological characteristics ^8^. Recent advancements in spatial transcriptomics (ST) now permit the simultaneous analysis of tissue architecture and gene expression, thereby providing new opportunities to investigate the relationship between morphology and molecular profiles. Previous spatial transcriptomic studies have demonstrated that molecular heterogeneity in prostate cancer is spatially organized across multifocal tumors and histologically distinct Gleason regions. Berglund et al. mapped transcriptomic variation across intact multifocal prostate cancer tissue, while Quan et al. showed that spatial transcriptional states align with histological patterns and change across Gleason score progression ^9,10^. More recent studies integrating single-cell and spatial transcriptomics have further resolved interactions between epithelial populations and the surrounding immune and stromal microenvironment, and these developments have been summarized in a recent review by Xu et al ^11^. Our study complements this literature by using gland architecture itself as a spatial reference system. Specifically, CurvSeq integrates gland boundaries, local curvature, pocket-like invaginations, and peri-glandular cellular composition with spatial molecular measurements, enabling gene and protein expression to be interpreted relative to local gland structure and its surrounding microenvironment. Nevertheless, the high dimensionality of ST data and the visual diversity of histological patterns necessitate computational approaches capable of identifying meaningful trends. Artificial intelligence (AI), particularly deep convolutional neural networks, has proven to be effective in extracting morphological features from tissue images and in distinguishing between normal and cancerous regions ^12^. This study integrates spatial transcriptomics with quantitative analyses of prostate gland morphology to investigate how the local tissue structure affects gene expression in prostate adenocarcinoma. By correlating spatially defined gene expression with morphological characteristics such as gland curvature and structural irregularities, we aim to identify transcriptional programs, particularly those associated with mechano-transduction, epithelial plasticity, and immune interactions, which are influenced by the physical organization of the tumor microenvironment. Our objective is to demonstrate how variations in gland structure, as indicated by Gleason patterns, correspond to underlying molecular states, ultimately revealing spatially localized biomarkers that could improve prognostication or guide targeted therapies in prostate cancer.

## Results

### CurvSeq/CurvSee Enables Morphomechanical Analysis of Gland Architecture

We developed a pipeline that quantifies prostate gland morphology and the surrounding cellular microenvironment and integrates these features with spatially resolved transcriptomic and proteomic data **(Fig. 1**). Using custom multiplex imaging and publicly available spatial transcriptomics such as Xenium, we segmented individual prostate glands and extracted their contours **(Fig. 1, Step 1)**.

**Figure 1.**
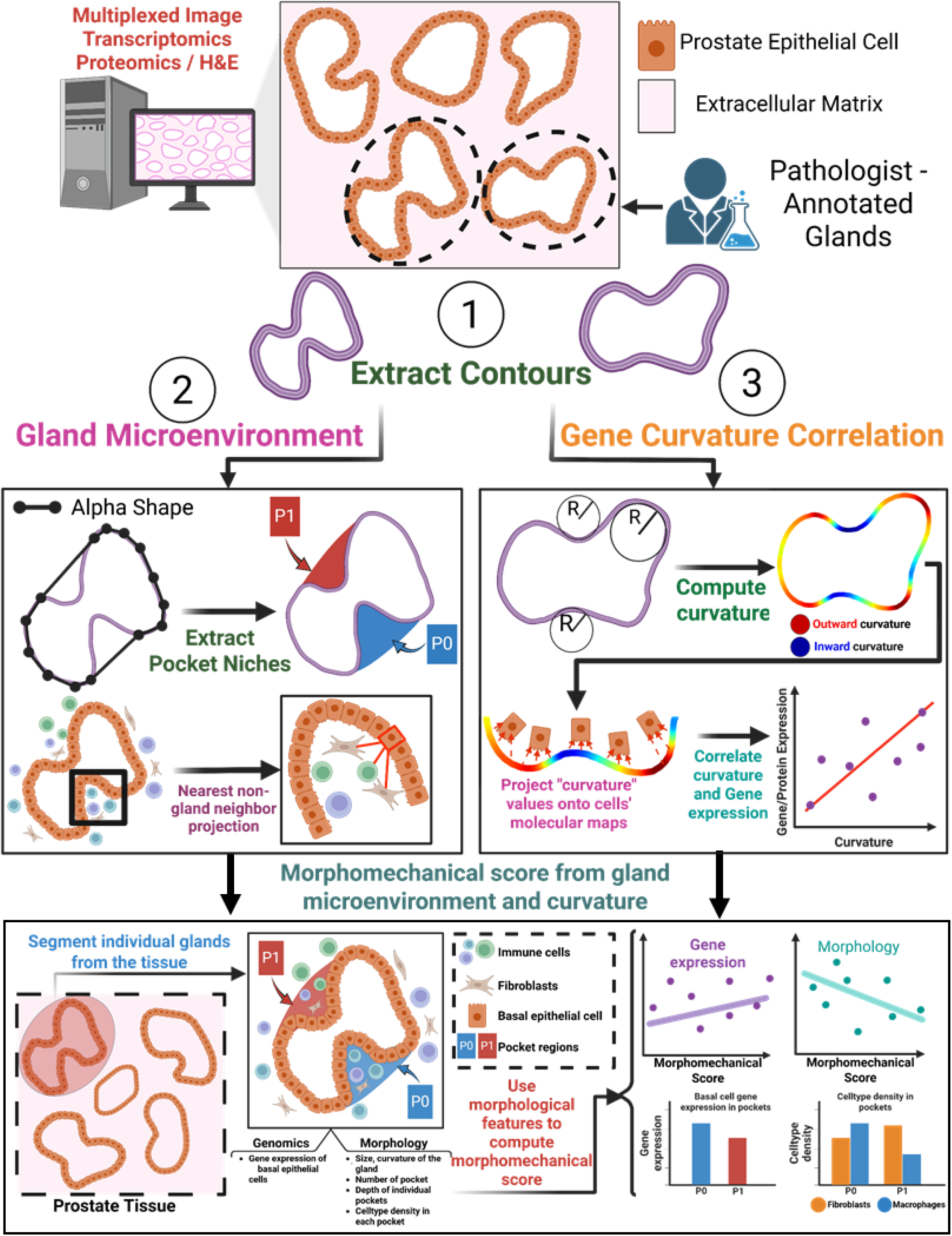
CurvSeq/CurvSee workflow for integrating gland morphology, curvature, molecular expression, and the gland microenvironment. Multiplexed image transcriptomics, spatial proteomics, and H&E-stained prostate tissue sections are used to identify and segment individual glandular structures. Gland contours are extracted from annotated epithelial boundaries and used as the geometric basis for downstream morphomechanical analysis. First, gland microenvironment features are quantified by detecting pocket-like invaginations using alpha-shape-based contour simplification and by projecting neighboring non-gland cells onto the closest gland boundary regions. This enables pocket-specific quantification of stromal, immune, fibroblast, and epithelial cell-type composition. Second, local boundary curvature is computed along each gland contour using radius-based curvature estimation, distinguishing outward and inward curvature regions. Curvature values are then projected onto spatial molecular maps, allowing correlation between local gland curvature and gene or protein expression in epithelial cells. Finally, gland-level morphology, pocket architecture, curvature features, local molecular expression, and surrounding cellular microenvironment measurements are integrated to compute a morphomechanical score. This score provides a quantitative axis for comparing gland states and identifying associations between gland architecture, epithelial gene/protein expression, and microenvironmental organization.

We computed the alpha shape of each extracted gland contour to define an outer geometric envelope of the gland. Pocket regions were then identified by calculating the geometric difference between the alpha-shape envelope and the original segmented contour, capturing inward concave regions along the gland boundary. Within these pocket regions, we quantified cell-type composition and density. We also analyzed the neighboring cellular microenvironment surrounding glandular epithelial cells to determine which external cell types were spatially associated with local gland regions **(Fig. 1, Step 2)**.

We next computed local curvature along the glandular boundary by fitting osculating circles to points along each extracted gland contour. The local curvature was defined as the inverse radius of the fitted osculating circle, with larger curvature values corresponding to sharper boundary bending. Curvature values were then projected onto nearby epithelial cells located inside the gland boundary. For each epithelial cell, the assigned curvature value represented the local boundary morphology closest to that cell. We then correlated projected curvature values with spatially resolved gene or protein expression to identify molecular programs associated with local gland morphology **(Fig. 1, Step 2)**. This enabled the integration of local curvature values as an additional feature assigned to cells within the gland, linking tissue shape with cellular gene expression profiles. The correlation indicates whether gene expression tends to be higher in specific gland regions based on boundary shape. A positive correlation suggests that gene expression is generally elevated in areas where the gland boundary is convex, while a negative correlation indicates higher expression in concave regions. A near-zero correlation implies no consistent spatial pattern. Importantly, a positive or negative correlation does not mean that every convex or concave region shows higher expression; rather, it represents an overall trend across the tissue.

Finally, we integrated gland curvature, pocket architecture, peri-glandular microenvironmental composition, and additional gland-level morphological features, including area, perimeter, circularity, and aspect ratio, to compute a continuous morphomechanical score. These features were assembled into a gland-level feature matrix and standardized before applying a diffusion-based ordering approach in Scanpy. The resulting score summarizes variation in gland shape, boundary organization, pocket formation, and surrounding cellular context into a single relative descriptor of gland morphomechanical state.

Additional details on image processing, curvature, and pocket analysis are provided in the **Methods** section.

### Morphomechanical profiling distinguishes benign and GG1 prostate glands

We began our analysis using a publicly available Xenium prostate adenocarcinoma spatial transcriptomics dataset. This dataset contains transcript counts for 5,000 genes together with morphological stains that enable visualization of glandular tissue architecture. Representative images highlight the heterogeneous folding patterns of individual glands and their corresponding spatial basal-cell gene-expression profiles (**Supplementary Fig. 1)**. Using pathologist-provided annotations, we stratified glands according to histological disease status. Glands were assigned to two groups: benign glands **(Fig. 2a, right)** and Gleason Grade Group 1 (GG1) glands **(Fig. 2a, left)**.

**Figure 2.**
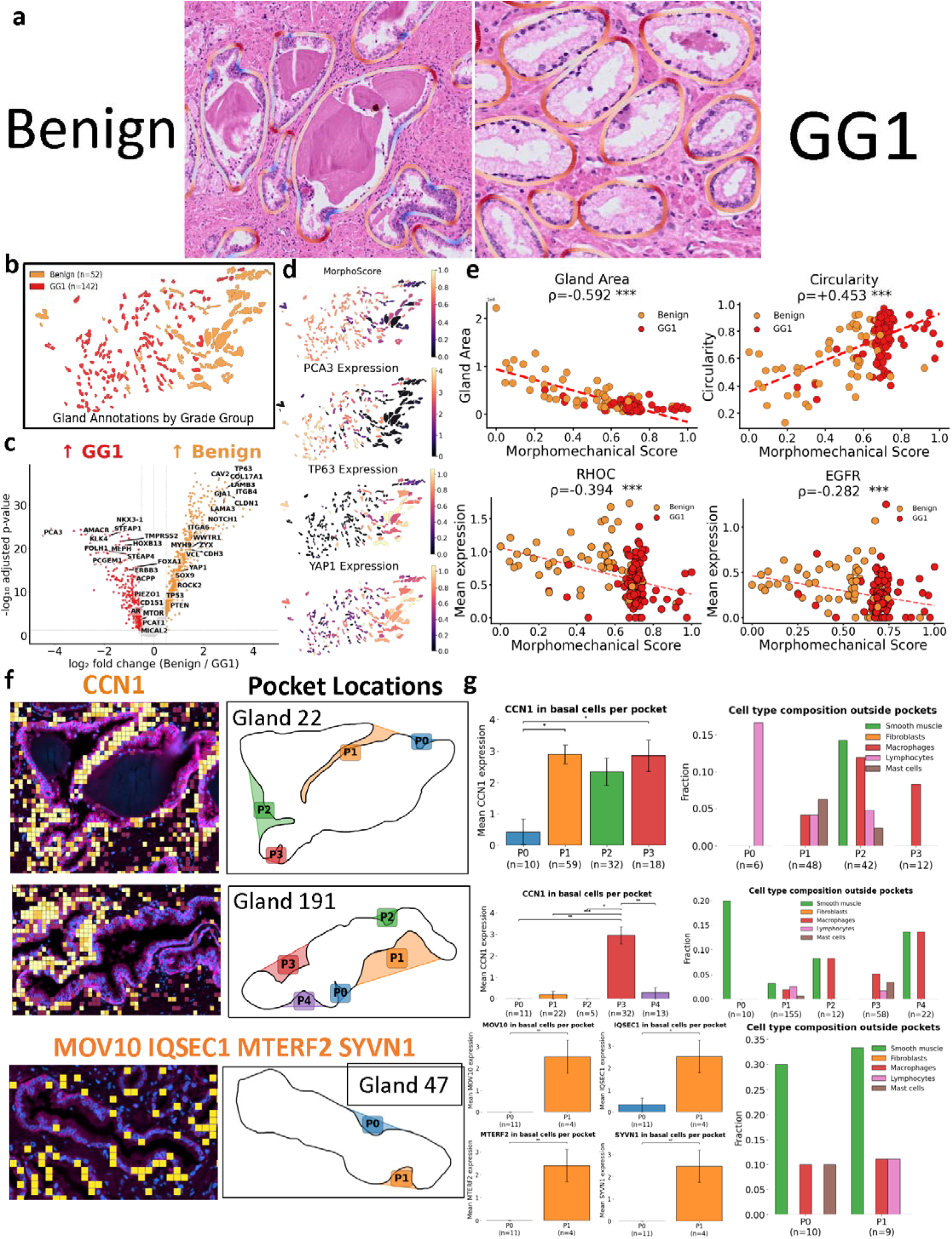
Spatial mapping of gland morphology reveals morphomechanical features associated with epithelial gene expression patterns. **(a)** Representative H&E images of benign and Gleason grade group 1 (GG1) prostate glands with gland boundaries overlaid, illustrating differences in gland architecture between non-malignant and low-grade cancer regions. **(b)** Spatial map of segmented gland annotations colored by grade group, including benign glands (n = 52) and GG1 glands (n = 142). **(c)** Differential gene expression analysis of basal epithelial cells comparing benign and GG1 glands. Genes enriched in GG1 are shown on the left, whereas genes enriched in benign glands are shown on the right. Selected prostate epithelial, tumor-associated, and basal/stromal-associated genes are labeled. **(d)** Spatial visualization of morphomechanical score and representative gene expression patterns across segmented glands, including PCA3, TP63, and YAP1. **(e)** Association between morphomechanical score and gland-level features or gene expression. Spearman correlation coefficients are shown; *** indicates statistical significance. **(f)** Representative single-gland examples showing heterogeneous pocket-specific expression. CCN1 expression is enriched in selected pocket regions in benign glands 22 and 191, while MOV10, IQSEC1, MTERF2, and SYVN1 show localized enrichment in a pocket region of GG1 gland 47. Pocket locations are shown as contour maps with individually labeled pocket regions. **(g)** Quantification of pocket-level basal epithelial gene expression and corresponding cellular composition outside each pocket.

We extracted contours from a total of 194 glands, including 52 benign glands and 142 GG1 glands. Visual inspection revealed that benign glands often displayed larger and more complex folded morphologies compared with GG1 glands **(Fig. 2b)**. In addition, benign glands generally exhibited more clearly defined luminal structures, whereas GG1 glands showed reduced or disrupted lumen organization, consistent with architectural changes associated with early prostate cancer progression ^13^. We then averaged basal-cell gene expression within each gland to obtain a single expression value per gland for each gene. Using these gland-level expression profiles, we performed differential expression analysis between benign and GG1 glands **(Fig. 2c)**. Gland-level comparison of basal epithelial expression profiles identified expected prostate cancer-associated genes, including PCA3 and AMACR, among genes enriched in GG1 glands ^14^. GG1-enriched genes also included prostate lineage and androgen-regulated epithelial markers such as FOLH1, KLK4, NKX3-1, STEAP1, TMPRSS2, HOXB13, and FOXA1 ^15–20^. In contrast, benign glands showed higher expression of basal epithelial and basement membrane-associated genes, including TP63, COL17A1, LAMB3, LAMA3, ITGB4, and ITGA6. Benign-enriched genes also included adhesion, junctional, and mechanotransduction-associated factors such as CLDN1, CAV2, GJA1, VCL, YAP1, WWTR1, ROCK2, and PIEZO1, consistent with a more organized basal-adhesive epithelial architecture and enrichment of mechanotransduction-associated genes.

We next computed the morphomechanical score for pathologist-annotated glands to evaluate whether this continuous morphological axis captured differences between benign and GG1 gland states. The largest benign gland by area was selected as the root for diffusion-based ordering. When the morphomechanical score was overlaid onto the tissue, lower-score glands were predominantly associated with benign annotations and higher TP63 expression, whereas higher-score glands were more frequently associated with GG1 annotations and increased PCA3 expression. In addition, glands with lower morphomechanical scores showed higher YAP1 expression, further supporting the enrichment of mechanotransduction-associated programs in benign glands **(Fig. 2d)**. Higher morphomechanical scores were enriched among GG1 glands and were associated with smaller gland area and increased circularity. In contrast, lower-score glands were more frequently benign and showed higher RHOC and EGFR expression. These results suggest that the morphomechanical score summarizes a coordinated architectural and epithelial-state axis, distinguishing larger, folded benign glands from smaller, more compact GG1 glands (**Fig. 2e**).

To evaluate how much local gland morphology and microenvironmental context were associated with gene-expression variation, we fitted linear models using morphological and cellular-neighborhood features as predictors of gene expression. These features included pocket membership, local concavity, pocket depth, cell depth within pockets, distance to the gland boundary, local epithelial density, and the composition of neighboring immune, stromal, and epithelial cells outside the pocket region. When all glands were pooled into a single model, the explanatory power was limited, and no genes showed high variance explained by these features. This suggested that morphology–expression relationships were not uniform across glands. We therefore repeated the regression analysis at the individual-gland level, allowing each gland to be modeled independently.

We fitted the regression on three benign glands that showed complex morphological folding **(Supplementary Figs. 2 and 3)**. In Gland 22, the strongest positive correlations were observed between pocket-associated genes and specific microenvironmental features. EMP1 showed the highest correlation with mast cells outside the pocket (ρ = 0.55), followed by CCN1 with outside mast cells (ρ = 0.53) and EMP1 with total cells in the pocket (ρ = 0.50). ERRFI1 also showed strong positive correlations with mast cells outside the pocket (ρ = 0.48), local epithelial density (ρ = 0.44), and total cells in the pocket (ρ = 0.40). In contrast, genes associated with differentiated prostate epithelial identity showed negative pocket-associated patterns, with NKX3-1 negatively correlated with pocket membership (ρ = −0.36), total cells in pocket (ρ = −0.32), and cell depth within pockets (ρ = −0.30). These results suggest that, in this gland, pocket-associated regions are associated with higher expression of CCN1, EMP1, EDN1, and ERRFI1, while showing reduced expression of NKX3-1 **(Supplementary Fig. 2a, b)**.

Finally, we analyzed cell-type composition and gene enrichment within individual pocket regions. Pocket-level analysis revealed substantial heterogeneity in CCN1 expression and adjacent microenvironmental composition across individual glands. CCN1 has been reported to be upregulated in human benign prostatic hyperplasia and to localize to hyperplastic epithelial and stromal compartments ^21^. In the first representative gland, most pocket regions showed elevated CCN1 expression, whereas the pocket with lower CCN1 expression was associated with a higher fraction of nearby lymphocytes (**Fig. 2f, 2g Gland 22**). In the second gland, CCN1 enrichment was restricted to a single pocket. This CCN1-high pocket lacked adjacent smooth muscle cells and was located near a neighboring gland that also showed elevated CCN1 expression, suggesting that pocket-associated expression may reflect both local pocket context and broader gland-neighborhood effects (**Fig. 2f, 2g, Gland 191**). We also examined cancer glands containing pocket-like regions. In one representative gland, MOV10, IQSEC1, MTERF2, and SYVN1 were elevated in one pocket region but not in another, showing localized transcriptional heterogeneity within the same gland (**Fig. 2f, g, Gland 47)**. Pocket-level microenvironmental analysis showed increased lymphocyte abundance in the gene-enriched pocket. These results suggest that pockets represent local morphologic niches whose molecular state is associated with gland-specific epithelial organization and microenvironmental context rather than a universal pocket-associated gene program

### Morphomechanical profiling identifies remodeling-associated pockets in benign prostate glands

We extended our analysis to a second publicly available Visium spatial transcriptomics dataset, which contains transcript counts for approximately 18,000 genes. This dataset also includes matched H&E-stained tissue images, which were used to manually annotate glandular regions. Based on pathologist-provided annotations, glands were stratified into two histological categories: benign glands **(Fig. 3a)** and Gleason Grade Group 4 (GG4) glands **(Fig. 4a)**. Because benign and GG4 glands exhibit markedly different morphologies, we analyzed them separately in the Visium dataset. Benign glands generally retained open luminal structures, whereas GG4 glands were more densely filled with tumor cells and lacked well-defined glandular lumens. To account for these architectural differences, downstream morphology, curvature, and gene-expression analyses were performed independently for benign and GG4 glands **(Supplementary Fig. 4)**.

**Figure 3.**
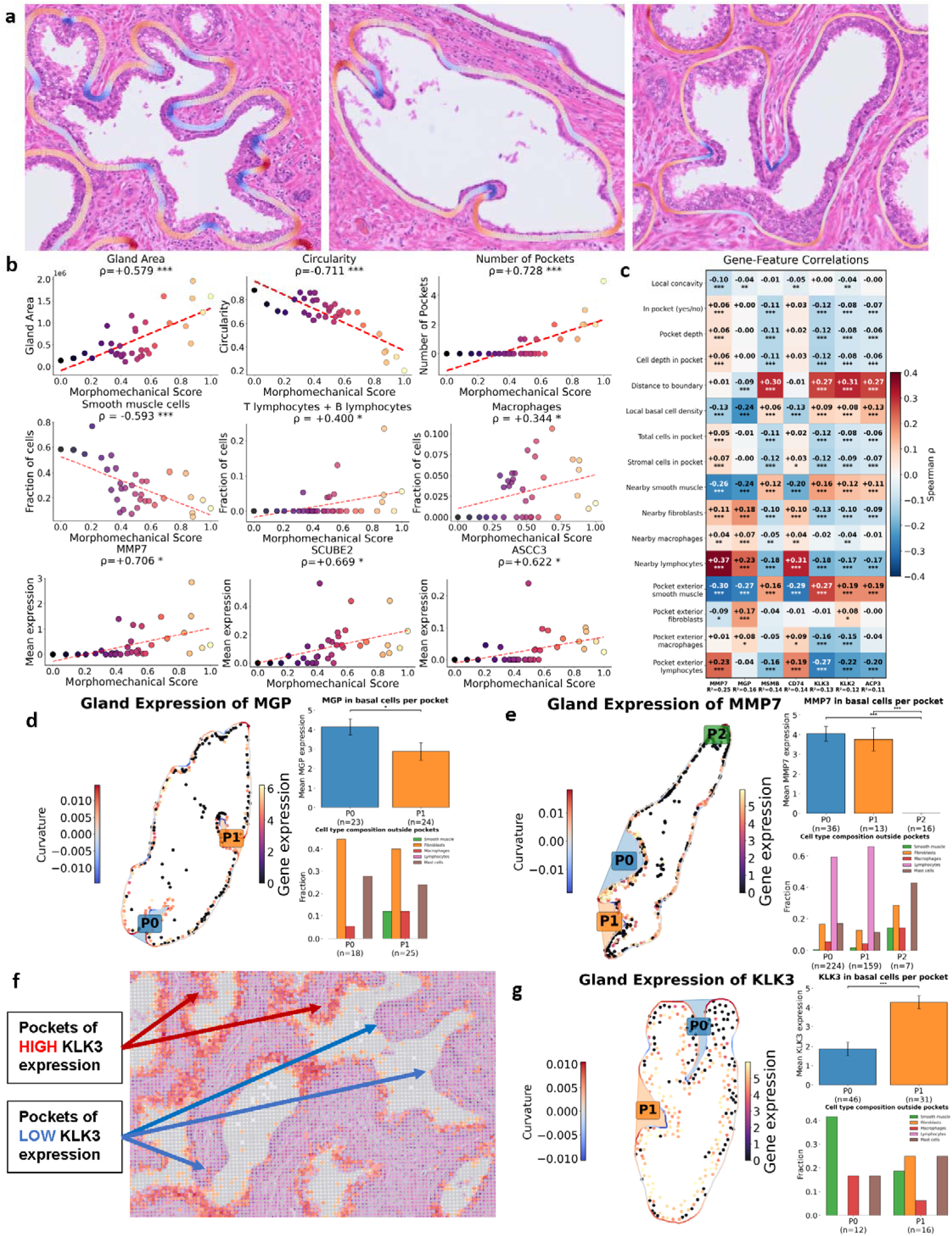
Morphomechanical variation in benign glands is associated with stromal remodeling, immune infiltration, and heterogeneous pocket-level epithelial gene expression. (a) Representative H&E images of benign prostate glands with segmented gland boundaries and pocket regions overlaid, illustrating variation in gland size, circularity, and epithelial invagination patterns. (b) Correlation between the morphomechanical score and gland-level morphology, cellular microenvironment composition, and epithelial gene expression. Higher morphomechanical score is associated with larger gland area, reduced circularity, increased number of pockets, reduced smooth muscle fraction, increased T and B lymphocyte and macrophage fractions, and increased expression of MMP7, SCUBE2, and ASCC3. Spearman correlation coefficients are shown for each comparison; * and *** indicate statistical significance. (c) Heatmap showing Spearman correlations between epithelial gene expression and gland morphology, pocket architecture, local epithelial organization, and peri-glandular microenvironmental features. MMP7, MGP, MSMB, CD74, KLK3, KLK2, and ACP3 show distinct associations with local concavity, pocket depth, distance to boundary, basal cell density, smooth muscle, fibroblast, macrophage, and lymphocyte features. (d) Representative gland showing MGP expression projected onto the gland contour and pocket regions. Quantification of MGP expression in basal cells across pockets and cell-type composition outside each pocket shows pocket-associated variation in expression and microenvironmental context. (e) Representative gland showing MMP7 expression and curvature projected along the gland boundary. Pocket-level quantification shows enrichment of MMP7 in selected pockets and corresponding differences in local cell-type composition outside pocket regions. (f) Spatial map highlighting pockets with high and low KLK3 expression, demonstrating that expression of luminal/prostate epithelial genes can vary across morphologically similar pocket regions within the same tissue area. (g) Representative gland showing KLK3 expression projected across pocket regions, with pocket-level quantification of KLK3 expression and adjacent cell-type composition.

**Figure 4.**
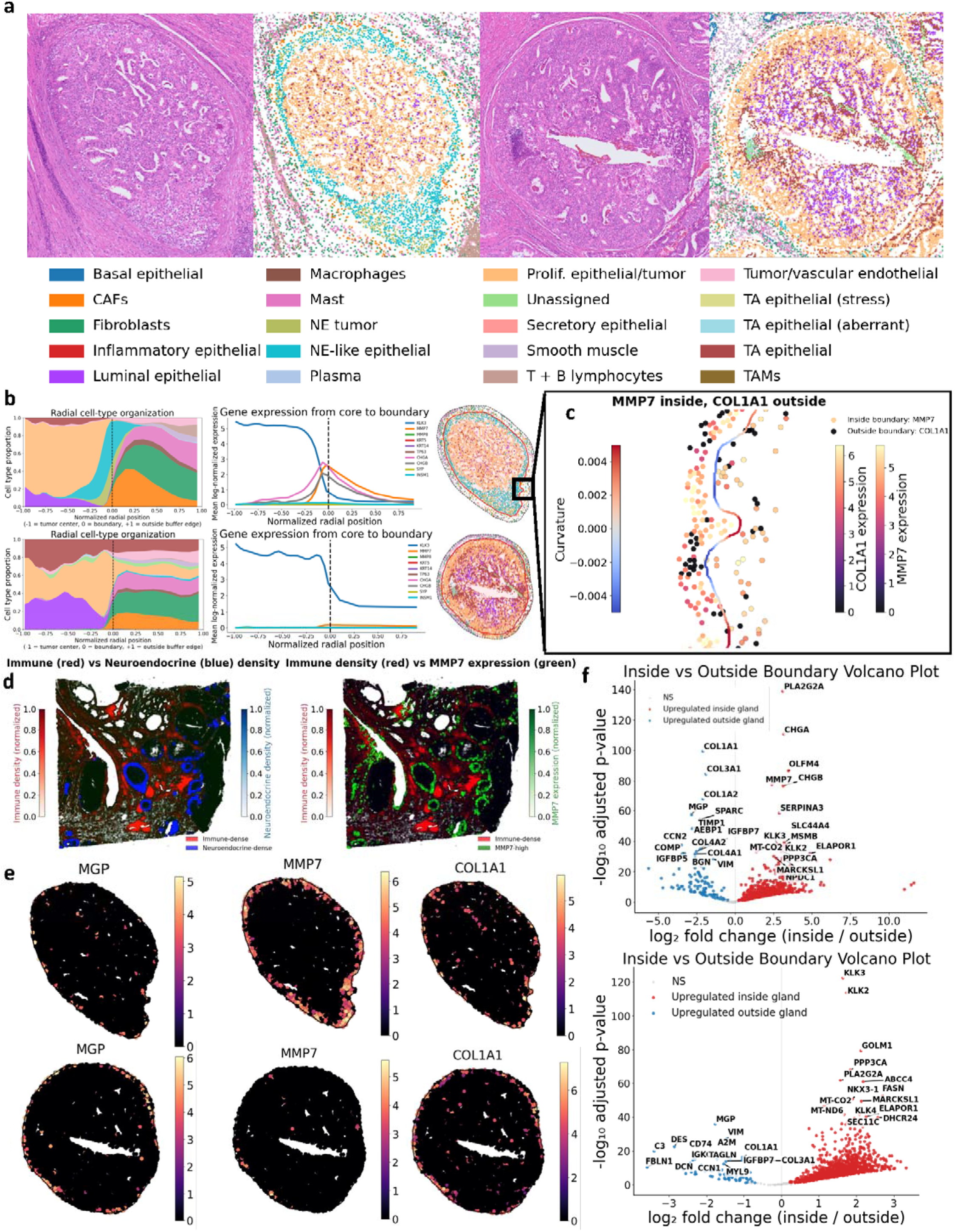
Spatial analysis of high-grade prostate cancer glands reveals boundary-associated cellular organization and compartment-specific gene expression. **(a)** Representative H&E images and corresponding spatial cell-type maps of GG4 prostate cancer regions, showing heterogeneous tumor architecture and distinct cellular organization within and around glandular structures. Cell-type annotations include basal epithelial cells, luminal epithelial cells, tumor-associated epithelial populations, proliferative epithelial/tumor cells, neuroendocrine-like epithelial cells, CAFs, fibroblasts, smooth muscle, macrophages, mast cells, plasma cells, T and B lymphocytes, and vascular/endothelial populations. **(b)** Radial cell-type organization and gene expression profiles from the gland core to the gland boundary and surrounding tissue. Two representative gland states are shown: glands with enrichment of neuroendocrine-like epithelial cells near the boundary and glands with a less sharply defined boundary organization.**(c)** Representative boundary-level analysis showing curvature together with epithelial MMP7 expression inside the gland boundary and COL1A1 expression outside the gland boundary, illustrating spatial coupling between epithelial and stromal molecular programs at the tumor–microenvironment interface. **(d)** Spatial density maps comparing immune cell density, neuroendocrine-like epithelial density, and MMP7 expression across the tissue section.**(e)** Representative gland-level maps of MGP, MMP7, and COL1A1 expression, showing compartmentalized expression patterns along gland boundaries and adjacent stromal regions. **(f)** Differential expression analysis comparing cells inside versus outside gland boundaries. Volcano plots show genes enriched inside glands, including epithelial and prostate-associated genes, and genes enriched outside glands, including stromal, extracellular matrix, immune, and fibroblast-associated genes.

We first applied the CurvSeq pipeline to benign glands in the Visium dataset. The smallest and most circular benign gland was selected as the root for diffusion-based morphomechanical ordering. In benign Visium glands, the morphomechanical score ordered glands along a continuous architectural axis. The score was strongly and significantly correlated with gland area (ρ = 0.579), circularity (ρ = −0.711), and number of pockets (ρ = 0.728), indicating that higher-score glands were larger, less circular, and contained more pocket-like structures. The score was also associated with changes in the peri-glandular microenvironment: smooth muscle cell fraction decreased with increasing score (ρ = −0.593), whereas immune populations increased, including T and B lymphocytes (ρ = 0.400) and macrophages (ρ = 0.344). At the gene-expression level, higher morphomechanical scores were associated with increased expression of MMP7 (ρ = 0.706) and SCUBE2 (ρ = 0.669) **(Fig. 3b)**. Together, these results indicate that more folded and pocket-rich benign glands are accompanied by reduced smooth muscle cell abundance, increased immune-cell presence, and higher MMP7 expression. MMP7 is a matrix metalloproteinase that promotes extracellular matrix remodeling and has been associated with aggressive and invasive prostate cancer phenotypes ^22^. In contrast, SCUBE2 is a secreted glycoprotein with reported tumor-suppressive functions, including inhibition of tumor cell migration, and is often downregulated in higher-grade tumors ^23,24^. The positive association of both MMP7 and SCUBE2 with the morphomechanical score suggests that folded benign glands may exhibit features of extracellular matrix remodeling while also retaining programs reported to have tumor-suppressive or differentiation-associated functions.

We then applied the regression framework jointly across all benign glands using the same morphological and microenvironmental features. Although overall model performance remained modest, consistent with substantial inter-gland heterogeneity, selected genes showed shared associations across glands. MMP7 showed the strongest overall model fit (R^2^ = 0.25), with higher expression associated with nearby lymphocytes (ρ = 0.37) and pocket-exterior lymphocytes (ρ = 0.23), and lower expression associated with nearby smooth muscle cells (ρ = −0.26) and pocket-exterior smooth muscle cells (ρ = −0.30). MGP also showed measurable variance explained by the model (R^2^ = 0.16), with negative associations with local basal-cell density (ρ = −0.24) and nearby smooth muscle cells (ρ = −0.24), and positive associations with nearby fibroblasts (ρ = 0.18) and pocket-exterior fibroblasts (ρ = 0.27). These results suggest that, despite gland-to-gland variability, selected remodeling-associated genes show consistent associations with stromal composition, immune proximity, and reduced smooth-muscle-rich microenvironments in benign glands **(Fig. 3c)**.

Finally, we examined pocket-level expression patterns in selected folded benign glands. In one representative gland, MGP expression was slightly reduced in a pocket with increased adjacent smooth muscle cell abundance **(Fig. 3d)**, consistent with the negative association between MGP expression and smooth muscle cell presence observed in the pooled regression analysis. In another gland, MMP7 expression was elevated in pockets with increased lymphocyte presence and little to no adjacent smooth muscle cells **(Fig. 3e)**. This pattern was also consistent with the pooled regression framework, in which MMP7 was positively associated with lymphocytes and negatively associated with smooth muscle cells. Together, these examples suggest that selected pocket-level expression patterns can reflect the broader morphology–microenvironment associations identified across benign glands, while still preserving substantial gland-specific heterogeneity.

Complex folded benign glands also showed heterogeneous KLK3 expression across pocket regions, with some pockets exhibiting high KLK3 expression and others showing reduced expression within the same gland **(Fig. 3f)**. KLK3 encodes prostate-specific antigen (PSA), a clinically important prostate epithelial marker used in prostate cancer assessment. In the pooled regression framework, KLK3 expression was negatively associated with nearby lymphocytes and positively associated with nearby smooth muscle cells. However, individual glands showed variable behavior, with some glands maintaining high KLK3 expression despite lymphocyte-rich neighborhoods and others displaying inverse or pocket-specific patterns **(Fig. 3g)**. These observations further support that pooled morphology–microenvironment associations capture broad trends, but individual glands can deviate substantially depending on their local epithelial state and microenvironmental context.

### Neuroendocrine-like GG4 glands show boundary-associated epithelial and stromal organization

GG4 glands showed distinct morphologies compared with benign glands. They were generally larger, more densely cellular, and exhibited more circular or solid glandular architectures, with poorly defined or absent luminal spaces **(Supplementary Fig. 4)**. Cell phenotyping revealed heterogeneity among GG4 glands. Some glands contained a peripheral layer of neuroendocrine-like epithelial cells along the gland boundary, whereas others lacked this border-associated neuroendocrine-like population **(Fig. 4a)**. Radial cell-type organization further revealed distinct spatial architectures among GG4 glands. In glands containing a neuroendocrine-like epithelial border, these cells were enriched near the gland boundary, separating the inner tumor compartment from the surrounding stromal and immune microenvironment. This boundary-associated neuroendocrine-like region was also accompanied by an increase in cancer-associated fibroblasts, suggesting a localized stromal remodeling niche around these cells. In contrast, GG4 glands lacking this neuroendocrine-like boundary showed a different radial organization, with tumor-associated and luminal epithelial populations distributed more broadly across the gland interior and a less defined boundary-associated epithelial layer **(Fig. 4b)**.

Gene-expression profiles along the normalized radial axis further supported the presence of a distinct boundary-associated epithelial state in GG4 glands. KLK3 expression was highest in the gland interior and decreased sharply near the gland boundary, whereas several boundary-associated markers increased near the gland edge. Neuroendocrine-associated genes, including CHGA, CHGB, SYP, and INSM1, showed localized enrichment around the gland boundary, consistent with the spatial accumulation of neuroendocrine-like epithelial cells observed in the cell-type analysis (**Fig. 4b**). At the gland boundary, neuroendocrine-like regions displayed thin protrusive extensions into the surrounding stromal compartment, with elevated MMP7 expression on the gland-internal side and increased COL1A1 signal in the adjacent external region (**Fig. 4c**). Neuroendocrine-like epithelial cells expressed higher levels of MMP7 than non-neuroendocrine epithelial populations, suggesting increased extracellular matrix-remodeling potential at these boundary-associated regions (**Fig. 4d**).Differential expression analysis between cells inside and outside the gland boundary further distinguished GG4 glands with and without neuroendocrine-like border organization.

In glands with a neuroendocrine-like boundary, inside-gland enriched genes included neuroendocrine-associated markers such as CHGA and CHGB, together with MMP7, supporting the presence of a boundary-associated neuroendocrine-like epithelial state with increased remodeling potential. In contrast, outside-boundary enriched genes included extracellular matrix and stromal-associated genes such as COL1A1, COL3A1, COL1A2, SPARC, TIMP1, and MGP, consistent with a collagen-rich stromal compartment surrounding these glands. In GG4 glands lacking a neuroendocrine-like boundary, inside-gland enriched genes were dominated by prostate epithelial markers such as KLK3, KLK2, NKX3-1, and FASN, whereas outside-boundary enriched genes included stromal and extracellular matrix-associated genes such as MGP, VIM, COL1A1, COL3A1, IGFBP7, and DCN (**Fig. 4e, f**).

### Spatial curvature proteomics distinguishes tumor grades

We next extended CurvSeq to multiplexed protein imaging across tissue sections representing different Gleason groups. Glands were segmented using pathologist-provided annotations, and gland-level morphological features, including curvature, area, and circularity, were computed for each annotated gland. These morphological features were integrated with protein-expression measurements to evaluate whether combined morphologic and molecular profiles could distinguish glandular disease states (**Fig. 5a**). We first analyzed a multiplexed proteomic dataset stained for P63, PSA, CK HMW, H3K9Ac, H3K4me2, H3K27me3, H4K12Ac, and AMACR/P504S. For each gland, protein expression was averaged across cells within the gland to generate a gland-level protein-expression profile. Heatmap analysis of these profiles showed separation between benign and higher-grade gland groups, with expected marker patterns, including higher P63 and CK HMW expression in benign glands and higher AMACR/P504S expression in malignant glands **(Fig. 5b**). We then applied a random forest classifier to predict the Gleason group of each gland using the combined gland-level morphology and protein-expression features. The classifier showed partial discrimination among several disease-associated gland groups but showed difficulty separating benign glands from non-cancerous tissue and benign glands located within cancer-containing tissue sections (**Fig. 5c**). Feature-importance analysis identified gland area as one of the most informative predictors, indicating that gland size contributes to distinguishing Gleason-associated gland states (**Fig. 5d**).

**Figure 5.**
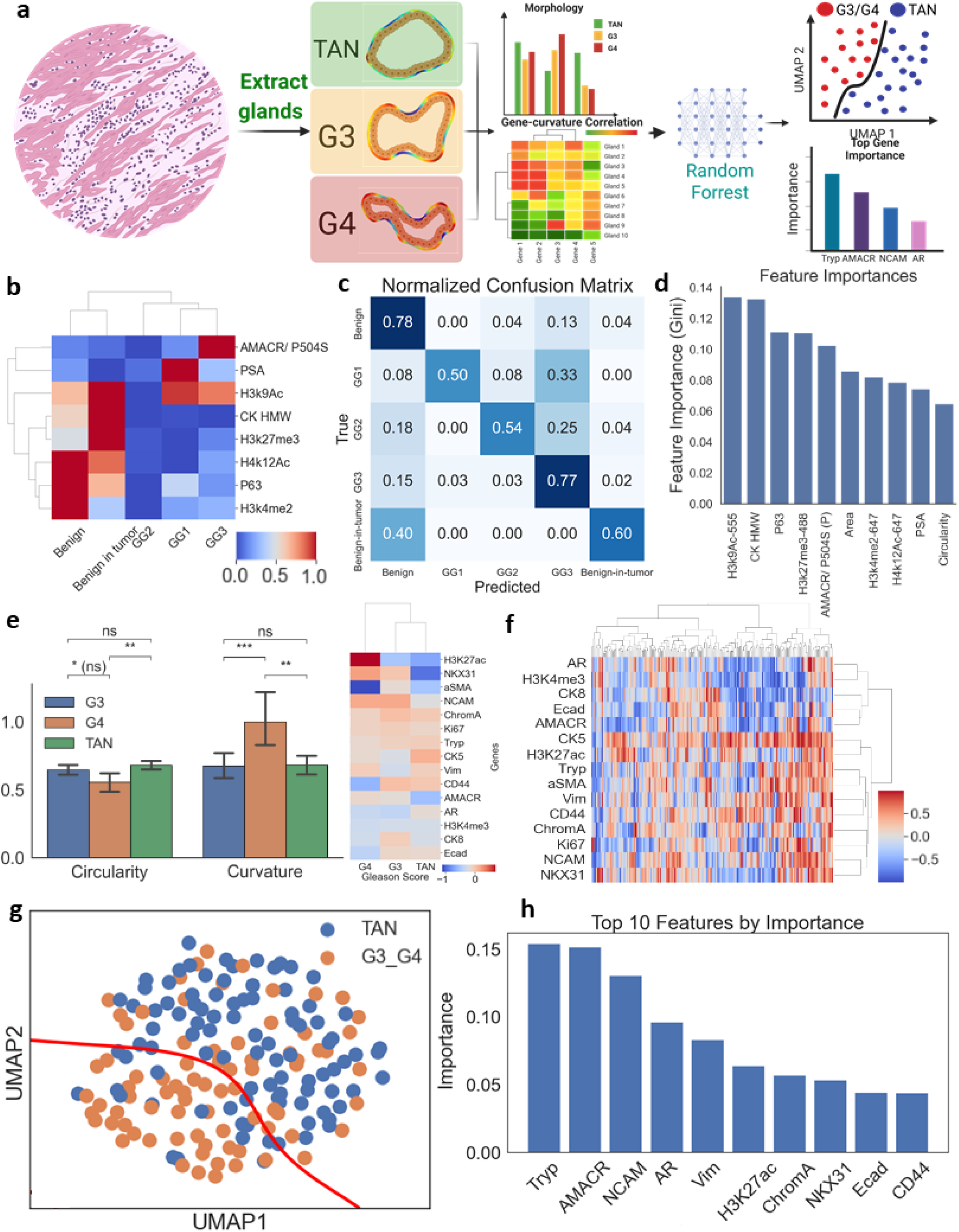
CurvSee integrates gland morphology and protein expression to classify prostate gland states across TAN and Gleason grade groups. **(a)** Schematic overview of the analysis workflow. Individual glands are extracted from multiplexed imaging or H&E-stained tissue sections and assigned different Gleason grades. Gland-level morphology, curvature-derived features, and protein expression profiles are integrated to assess gene/protein–curvature relationships and to classify gland states using random forest models. **(b)** Heatmap of mean protein marker expression across gland categories, showing distinct epithelial, basal, tumor-associated, and chromatin-associated protein profiles across benign, benign-in-tumor, GG1, GG2, and GG3 glands. **(c)** Normalized confusion matrix from a random forest classifier trained using combined expression and morphology features, showing classification performance across benign, GG1, GG2, GG3, and benign-in-tumor glands. **(d)** Feature importance analysis from the random forest model, identifying protein and morphology features that contribute to gland-state classification, including H3K9Ac, CK HMW, P63, H3K27me3, AMACR/P504S, PSA, and circularity. **(e)** Quantification of curvature variation and circularity across TAN, G3, and G4 glands, showing grade-associated differences in gland morphology. The accompanying heatmap summarizes protein expression differences across Gleason groups. **(f)** Hierarchical clustering of gland-level protein-curvature correlations across markers, including AR, H3K4me3, CK8, E-cadherin, AMACR, CK5, H3K27ac, Tryptase (Tryp), αSMA, vimentin, CD44, chromogranin A, Ki67, NCAM, and NKX3.1. **(g)** UMAP representation of gland-level features separating TAN from combined G3/G4 glands, indicating partial stratification of tumor-associated gland states based on integrated morphology and protein measurements. **(h)** Top-ranked features contributing to TAN versus G3/G4 classification, including Tryp, AMACR, NCAM, AR, vimentin, H3K27ac, chromogranin A, NKX3.1, E-cadherin, and CD44.

We applied our pipeline to an independent dataset comprising tissues from patients with Gleason grade 3 (G3) and grade 4 (G4) tumors, as well as Tumor Adjacent Normal (TAN) tissue. As a first step, we manually segmented the glands and quantified two key morphological features: curvature variation and circularity (see Methods) to assess structural differences among prostate glands classified as G3, G4, or TAN. While TAN and G3 glands exhibited no significant differences in either morphological metric, G4 glands differed significantly from both TAN and G3. Specifically, G4 glands were, on average, less circular and showed greater curvature variations along their boundaries (**Fig. 5e**). Gland area also varied across benign, GG1-GG3, and benign-in-tumor gland categories (**Supplementary Fig. 5a**).

We aggregated the protein expression within each gland as in the previous section. The clustermap (**Fig. 5e**) effectively grouped the G3 and TAN glands together, distinguishing them from the G4 glands. This separation highlights distinct expression patterns in several genes, including *H3k27ac, CD44, NKX3-1*, and NCAM. Specifically, G4 glands showed elevated *H3K27ac* and *NCAM* expression, along with reduced *NKX3-1* levels compared to G3 and TAN. Following the analysis of glandular protein expression, we generated a heatmap by performing linear regression between the expression level of each protein and the corresponding projected gland curvature (**Fig. 5f**). We then projected the regression-derived values into a latent space using UMAP and labeled glands according to their classification (G3 or G4 and TAN). Visually, the projection revealed partial separation between TAN glands and the combined G3/G4 group (**Fig. 5g**). Although some overlap existed between G3/G4 and TAN glands, G3/G4 glands occupied a distinct region of the embedding space. To formalize this separation, we applied Support Vector Classification (SVC) and plotted a decision boundary distinguishing the two groups.

To assess whether curvature–protein expression relationships contained information that could distinguish tumor from non-tumor glands, we trained a Random Forest classifier to separate TAN from Gleason-labeled glands using the heatmap from **Fig. 5f**. The classifier achieved an accuracy of 74% on test data, with an area under the ROC curve (AUC) of 0.74, suggesting moderate predictive capacity (**Supplementary Fig. 5b)**. Interestingly, the confusion matrix shows that the random forest classifier identified TAN glands more accurately than glands belonging to Gleason-labeled tissues (**Supplementary Fig. 5c)**. Feature importance analysis identified *Tryp, AMACR, NCAM, H3K27ac*, and *NKX3-1* as top contributors to classification **(Fig. 5h)**. Tryptase (Tryp) is a mast-cell marker, and mast-cell infiltration has been investigated as a prognostic feature in prostate cancer ^25^. NCAM (Neural Cell Adhesion Molecule) plays a crucial role in cell–cell adhesion and is involved in various developmental processes; however, its re-expression has been observed in certain malignancies. In prostate cancer, increased NCAM expression has been linked to neuroendocrine differentiation and more aggressive tumor behavior ^26,27^.

### Tissue stiffness and curvature exhibit spatial associations in *Foxa1* knockout and control tissues

In this study, we analyzed a dataset comprising mouse prostate tissues in which the *Foxa1* gene has been knocked out in combination with tumor suppressor gene *Pten*, alongside control mouse prostate tumor tissues in which *Pten* alone is knocked out and *Foxa1* remains intact. FOXA1 is a pioneer transcription factor involved in organ development, cell differentiation, and tumor progression. Previous studies have reported that *Foxa1* loss induces *TGFB3* expression and activates TGF-β signaling ^28^. We analyzed a cohort of prostate tissues from mice stratified by Foxa1 status, with “P” (Pten Knockout) indicating tissues retaining Foxa1 expression and “PF” (Pten/Foxa1 double Knockout) indicating tissues lacking Foxa1. P mice were used as the control group for comparison with PF mice. The dataset also included labels indicating the week of the mouse’s lifespan, measured in weeks at the time of sampling. The dataset included two P mice and three PF mice, sampled at weeks 12, 13, and 19 (**Supplementary Fig. 6**). **Fig. 6a** displays the manually segmented glands generated using QuPath. Boundary curvature was quantified and mapped onto the neighboring segmented cells. The left panel depicts P12 sample, whereas the right panel illustrates the PF12 sample. The glands were segmented from the lateral-ventral regions (LVP**) (Supplementary Fig. 7)**.

**Figure 6.**
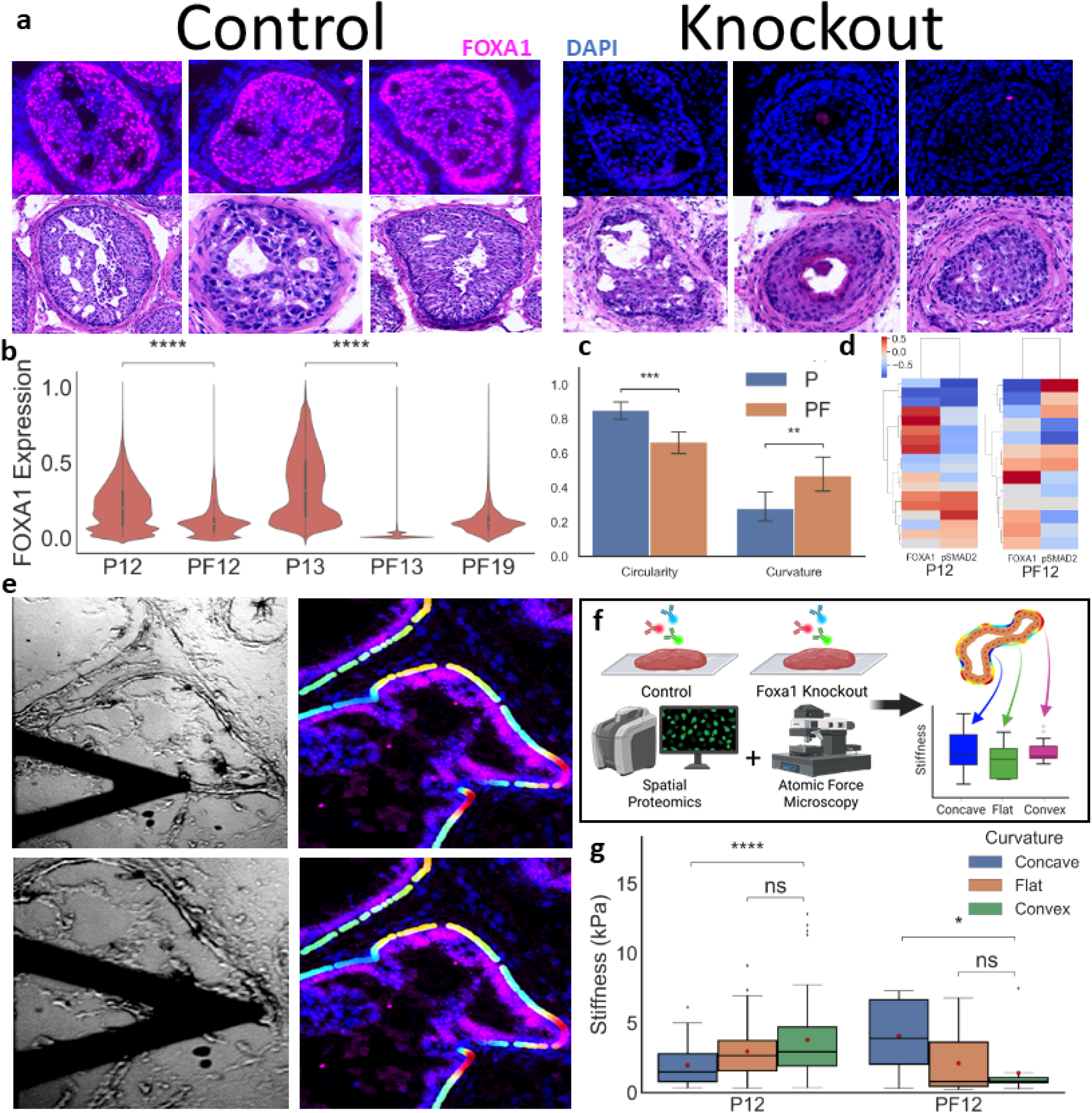
FOXA1 loss is associated with altered gland morphology, protein organization, and curvature-dependent tissue stiffness. **(a)** Representative FOXA1 immunofluorescence and matched H&E images of control and FOXA1-knockout prostate glands. FOXA1 is shown in magenta and nuclei are labeled with DAPI in blue. **(b)** Violin plots showing FOXA1 expression across control and FOXA1-knockout samples at the indicated time points. FOXA1 expression is reduced in PF12 and PF13 tissues relative to their corresponding controls, while PF19 displays residual FOXA1 expression. **(c)** Quantification of gland circularity and curvature in control P and FOXA1-knockout PF tissues. Control glands exhibit higher circularity, whereas knockout glands exhibit greater curvature. **(d)** Heatmaps showing gland-level FOXA1 and pSMAD2 expression in P12 and PF12 tissues, illustrating altered spatial protein organization between control and knockout glands. **(e)** Representative atomic force microscopy images and corresponding DAPI/FOXA1 fluorescence images used to align mechanical measurements with gland architecture and protein expression. **(f)** Schematic of the integrated spatial proteomics and atomic force microscopy workflow. Control and FOXA1-knockout tissues were analyzed to compare stiffness across concave, flat, and convex gland boundary regions. **(g)** Quantification of tissue stiffness across concave, flat, and convex regions in P12 and PF12 samples. In P12 tissues, convex regions were stiffer than concave regions, whereas PF12 tissues showed higher stiffness in concave regions and reduced stiffness in convex regions. Statistical significance is indicated by asterisks; ns, not significant. Together, these results show that FOXA1 loss is accompanied by altered gland morphology, spatial protein organization, and mechanical patterning along gland boundaries.

The samples were stained for FOXA1 and pSMAD2. As expected, we observe a reduction in cellular expression of the FOXA1 protein in the PF samples (**Fig. 6b; Supplementary Fig. 8b**). In the P samples, we observe that the expression of FOXA1 is more localized in and around the nuclei, while PF mice exhibited an overall loss of FOXA1 nuclear staining. We observed a reduction in circularity and increase in curvature in week 12 PF mice (**Fig. 6c**).

We applied the CurvSee pipeline to the P12 and PF12 samples to assess differences in protein–curvature correlations between them (**Fig. 6d**). The P12 samples contained a total of 18 segmented prostate glands, while the PF12 samples included 13 glands in total. The pSMAD2–curvature relationship varied with genotype and age: P12 showed a strongly negative correlation, PF12 showed a near-zero correlation, and PF13 showed a lower correlation than P13 (**Supplementary Fig. 8a**). Although the PF samples appear to show a high correlation for FOXA1, this value is not biologically meaningful since *Foxa1* was knocked out in these tissues, resulting in minimal variability in its expression. Interestingly, while the control samples show little variability across age groups in the correlation with *pSMAD2*, the PF samples exhibit markedly higher variability.

We used Atomic Force Microscopy (AFM) to measure tissue stiffness along the borders of prostate glands in both *Foxa1* control and *Foxa1*-knockout samples, focusing on regions of negative and positive curvature **(Fig. 6e, f)**. Negative curvature corresponded to inward-bending, concave regions of the gland boundary, whereas positive curvature corresponded to outward-bending, convex regions. Stiffness was calculated by fitting a Hertzian model to the cantilever deflection curves. Only curves with an overall goodness of fit of R^2^ > 0.95 were retained for analysis. More details on the procedures and parameters are given in the **Methods** section. In fresh tissues, the *Foxa1* control sample (P12) displayed a positive trend, with higher stiffness values observed in regions of higher curvature. In contrast, the Foxa1-knockout sample (PF12) exhibited the opposite pattern, showing greater stiffness in regions of lower curvature (**Fig. 6g**). To further characterize the mechanical properties of the tissues, we performed AFM-based measurements of height and adhesion on the same regions analyzed for stiffness (**Supplementary Figs. 9 and 10**). Sample P12 showed more uniform adhesion, whereas PF12 exhibited greater variability, with adhesion forces changing across regions of different curvature (**Supplementary Fig. 11**).

### Stiffness mapping reveals altered gland-boundary mechanics following FOXA1 loss

We extended our analysis using the PAVONE nanoindenter to directly measure tissue stiffness in control P12 and Foxa1-knockout PF12 prostate tissues. Rather than restricting measurements to predefined high- or low-curvature regions, we performed whole-region scans to generate spatial maps of tissue indentation and stiffness. The indentation maps were used to identify gland boundaries and distinguish tissue regions from the underlying glass substrate. This step was necessary because local tissue thickness influences the measured indentation response, and inclusion of substrate-associated measurements could artificially alter the estimated stiffness values (**Fig. 7a**).

**Figure 7.**
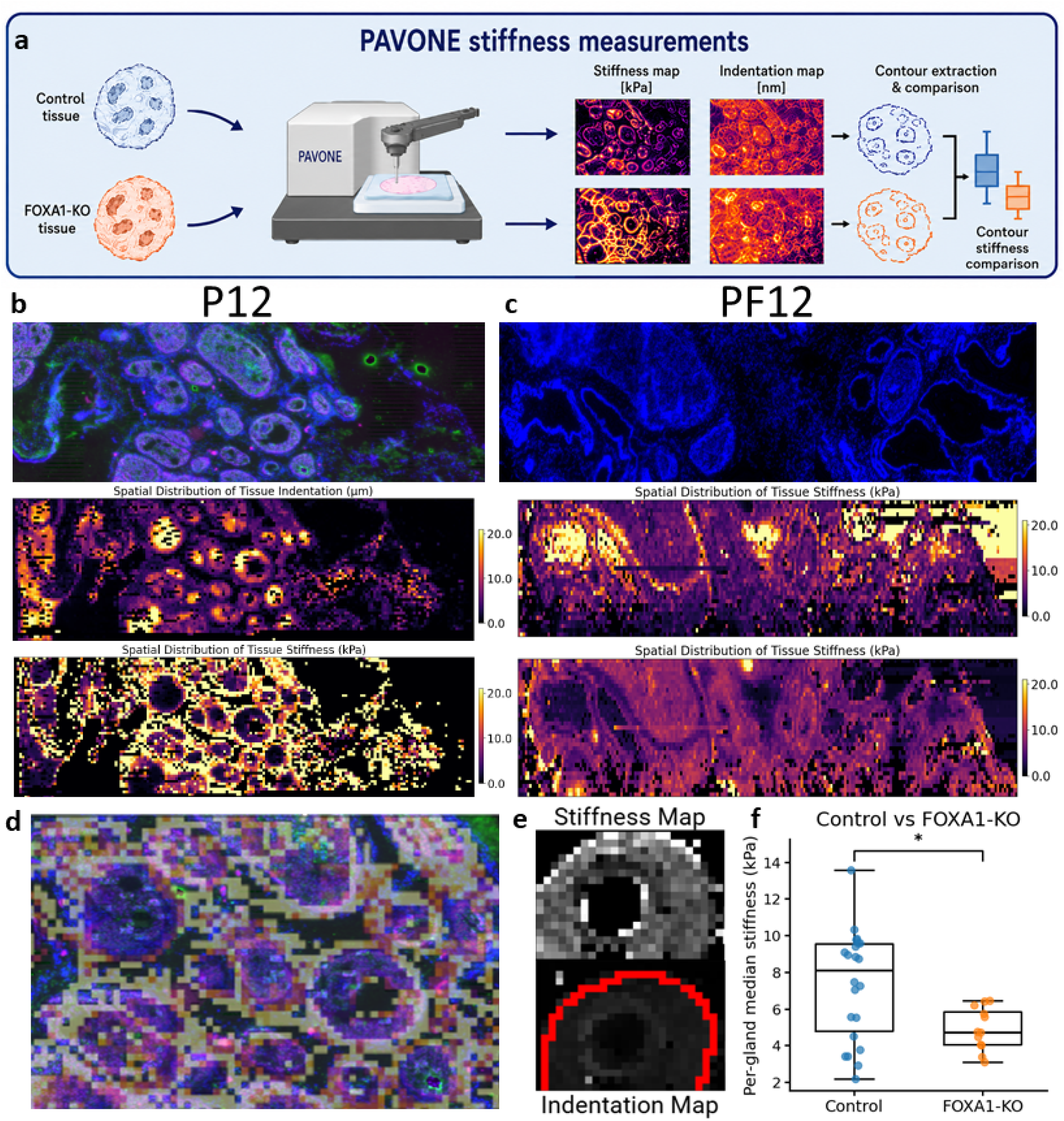
Whole-region PAVONE mapping reveals altered gland-associated stiffness in FOXA1-knockout prostate tissue. **(a)** Schematic of the PAVONE nanoindenter workflow used to compare control and FOXA1-knockout tissues. Whole tissue regions were scanned to generate spatial stiffness and indentation maps, followed by gland contour extraction and comparison of gland-associated mechanical properties between conditions. **(b)** Representative control tissue region showing the corresponding fluorescence image, indentation map, and stiffness map. The indentation map preserves the spatial organization of glandular structures and was used to distinguish tissue regions from the underlying substrate. **(c)** Representative Foxa1-knockout tissue region showing the corresponding DAPI image and indentation and stiffness maps. Spatially heterogeneous mechanical properties are observed across the scanned tissue region. **(d)** Overlay of the fluorescence image with the stiffness map, illustrating spatial alignment between gland architecture and local mechanical measurements. **(e)** Representative gland identified in the stiffness and indentation maps. The gland contour, shown in red, was extracted from the indentation map and transferred to the stiffness map to isolate gland-associated measurements while excluding surrounding or substrate-dominated regions. **(f)** Quantification of median gland stiffness in control and FOXA1-knockout tissues. FOXA1-knockout glands display lower median stiffness than control glands. Individual points represent glands, and the asterisk indicates statistical significance.

Representative control and Foxa1-knockout tissue regions showed that PAVONE-derived indentation and stiffness maps preserved the spatial organization of glandular structures (**Fig. 7b,c,d**). Areas corresponding to gland contours could be identified in the indentation maps and subsequently matched to the stiffness maps, enabling extraction of gland-associated mechanical measurements while excluding substrate-dominated regions. The stiffness maps also revealed substantial spatial heterogeneity within and between glands, indicating that local mechanical properties varied across the scanned tissue regions.

Comparison of gland-boundary stiffness between the two groups showed that control glands exhibited higher average stiffness and greater variability than Foxa1-knockout glands (**Fig. 7e,f**). In contrast, Foxa1-knockout glands displayed a narrower distribution with lower contour-associated stiffness. This difference was statistically significant, consistent with an association between Foxa1 status and altered mechanical properties at the gland boundary. These findings provide direct mechanical measurements that complement the morphologic and molecular analyses and suggest that architectural differences between Foxa1-intact and Foxa1-knockout tissues are accompanied by differences in local tissue stiffness.

## Discussion

In this paper, we demonstrate that our pipeline, which extracts glandular shapes from immunofluorescence (IF) images, enables the integration of morphological information across multiple datasets. This approach allows us to examine relationships between gland architecture, molecular expression, and mechanical properties in prostate tissue. Using our Curvseq pipeline, we found groups of genes that exhibit distinct spatial arrangements within prostate glands, with genes corresponding to different molecular pathways and functional behaviors. These spatially organized genes indicate coordinated variation in molecular activity relative to local gland architecture. Using our CurvSee pipeline, we integrated spatial proteomics with detailed gland morphology to investigate how tissue structure and protein expression varies across Gleason-associated prostate cancer states. Curvature-informed mapping of tumor-adjacent normal (TAN) and malignant Gleason-graded glands (G3–G4) captured distinct morphological and molecular signatures. Random forest classification identified curvature-associated proteins that distinguish normal from tumor glands. Additionally, our results suggest that altering glandular transcriptional programs can be accompanied by changes in the mechanical organization of prostate glands (**Fig. 8**). Importantly, CurvSeq is not intended to outperform existing spatial omics methods in classification accuracy or predictive performance. Its main contribution is to introduce gland architecture as a spatial reference system that integrates local curvature, pocket-like structures, gland-level morphology, and the surrounding cellular microenvironment with spatial molecular measurements. This allows gene and protein expression to be interpreted relative to the organization of the gland and its neighboring stromal and immune compartments, rather than only by cell type or broad tissue region. CurvSeq therefore provides a complementary framework for examining how gland shape and local microenvironmental context are associated with molecular heterogeneity across prostate tissues.

**Figure 8.**
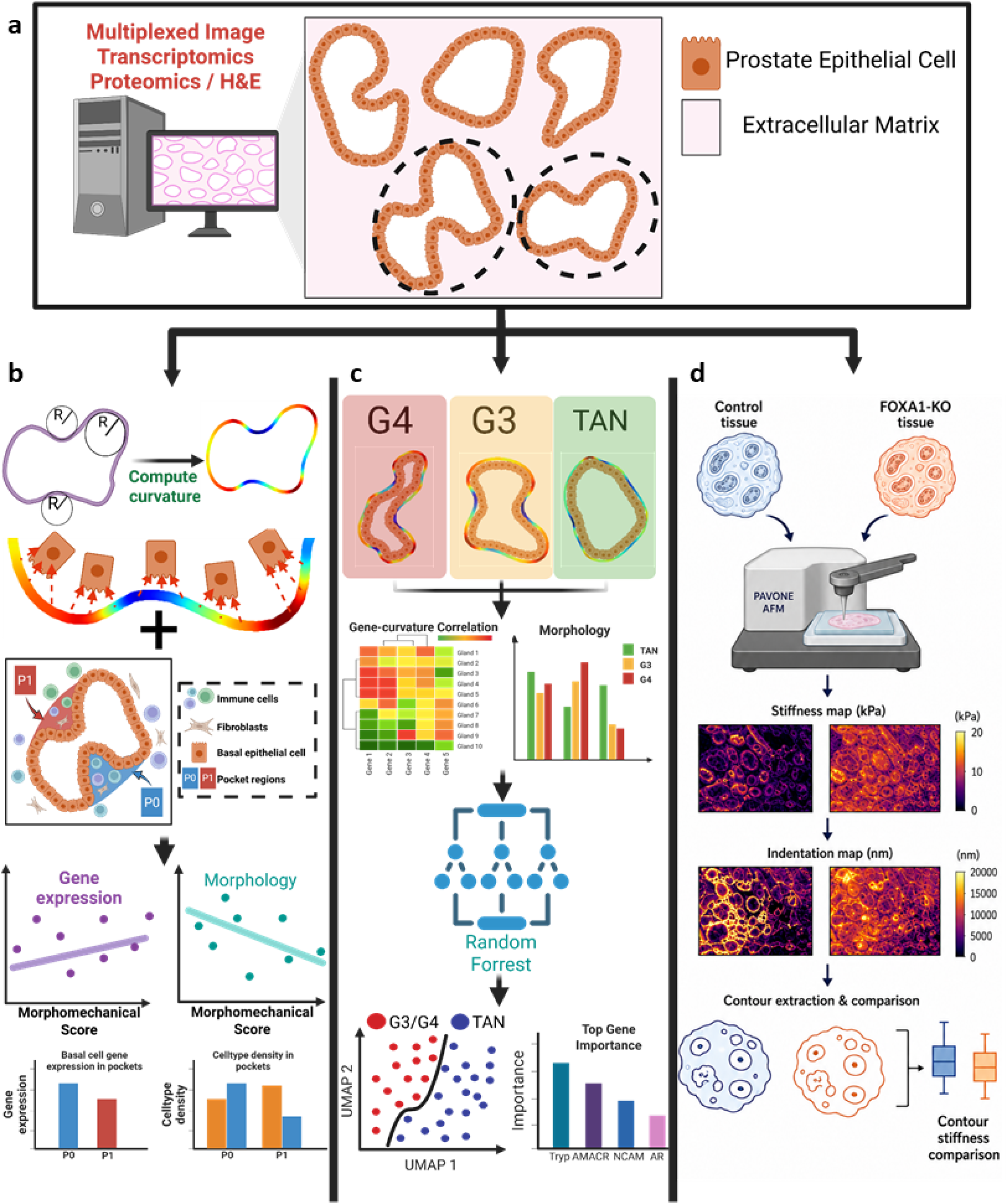
Integrated overview of curvature-based morphomechanical analysis across spatial transcriptomics, spatial proteomics, and atomic force microscopy. **(a)** Multiplexed image transcriptomics, spatial proteomics, and H&E-stained tissue sections are used to identify and segment individual prostate glands for downstream morphologic and mechanical analyses. **(b)** Curvature is computed along gland contours and integrated with epithelial molecular measurements, pocket architecture, and the surrounding cellular microenvironment. These features are combined into a morphomechanical score that captures gland-level variation in gene expression, morphology, and pocket-associated cell-type composition. **(c)** Spatial proteomics extends the framework across tumor-adjacent normal (TAN), Gleason grade 3 (G3), and Gleason grade 4 (G4) glands. Gene/protein–curvature relationships and gland morphology are integrated in random forest models to evaluate separation of TAN and tumor-associated glands and to determine the relative contribution of molecular and morphologic features. **(d)** Whole-region PAVONE atomic force microscopy is used to compare control and FOXA1-knockout prostate tissues. Spatial stiffness and indentation maps are generated, gland contours are extracted, and gland-associated stiffness measurements are compared between conditions. Together, this framework integrates gland shape, local curvature, molecular organization, cellular microenvironment, and tissue mechanics across complementary spatial profiling platforms.

For a long time, Gleason grading has relied on specific morphological features, including gland fusion, loss of lumen, and irregular architecture. However, the molecular basis of these features is not fully understood. The connection between gland curvature and transcriptional modules could provide a clearer understanding of how morphology and gene regulation are linked. The convex and concave areas of glands show different local stress patterns and cell packing, which may change the shape of nuclei, how chromatin is organized, and the availability of transcription factors.

Several computational approaches have been developed to study spatial correlations in tissue imaging and spatial omics. For example, neighborhood analysis and spatial autocorrelation metrics such as Moran’s I or Ripley’s K have been widely applied to identify patterns of cell–cell proximity, clustering, or co-localization of molecular features ^29,30^. These methods offer valuable insight into microenvironmental structure and identify transcriptional (protein and RNA) spatial patterns. However, they primarily focus on proximity to neighboring cells. These algorithms fail to capture the physical architecture of tissues due to the limited structural information available. This motivated us to incorporate curvature as an additional spatial descriptor. By projecting curvature values onto neighboring cells and correlating them with gene or protein expression, our framework links potential morphological stress states of the tissue to the molecular activities of the neighboring cells.

Recent advances in physics-informed artificial intelligence have emphasized the integration of geometric priors and curvature-based constraints into learning frameworks to better capture the underlying physical laws of complex systems. Geometry-aware neural solvers now incorporate manifold embeddings and curvature-adaptive operators to model phenomena on irregular or curved domains with high fidelity, bridging the gap between numerical physics and data-driven learning. Similarly, curvature-aware models in high-energy physics and geometry-aware physics-informed neural networks have demonstrated that incorporating differential-geometric features or transformed domains can improve learning and physical modeling on non-Euclidean or geometrically complex spaces ^31,32^. In this context, glandular curvature in prostate tissue represents a biological analog of such physically grounded geometry: the morphological contours of glands can be converted into a quantitative curvature field. While our approach is grounded in physical principles, it is important to distinguish it from the formal domain of physics-informed artificial intelligence, such as physics-informed neural networks (PINNs), which explicitly embed governing equations into machine-learning architectures.

Multiple computational approaches exist to quantify curvature and its associated energetic landscape, ranging from differential-geometry–based estimators (e.g., polynomial fitting or finite differences) to field-based methods such as Laplacian curvature derived from binary masks. These geometric measures can then be physically interpreted through models like the Helfrich (bending) energy formulation, which relates local curvature to membrane tension and bending rigidity, enabling a direct link between morphology and underlying mechanical energetics ^33–35^ (**Supplementary Fig. 12**).

A key objective of our study is to quantitatively assess whether and where specific gland regions display elevated gene expression compared to others (**Supplementary Fig. 13a)**. High-curvature regions within glands display variable gene expression patterns, with some areas showing upregulation and others downregulation. We applied linear regression to scatter plots of individual cells within each gland, using curvature values as the x-axis and the expression of a selected gene as the y-axis. A common feature of transcriptomics data is that many cells exhibit zero counts for specific genes, and the expression values are inherently discrete. This sparsity and discreteness reduce the reliability of fitting continuous models such as linear regression, often leading to weaker correlations (**Supplementary Figs. 14-16**). By clustering cells according to their spatial proximity and correlation values (**see Methods**), and then aggregating expression within each cluster, we reduce both the total number of points on the scatter plot and the frequency of zero-expression values.

We first performed simulations of gland-like structures by generating random shapes and populating them with cells along the boundary. To each cell, we assigned a fictive RNA expression profile following a Zero-Inflated Negative Binomial (ZINB) distribution. We selected ZINB because, upon examining the distribution of RNA expression within actual glands, it most closely resembled a ZINB pattern (**Supplementary Fig. 17a**). We then selected random regions within the simulated glands and modified their expression levels by adding values drawn from a normal distribution, thereby creating localized areas of higher or lower expression (**Supplementary Fig. 17**). We then applied CurvSeq (**Fig. 1**) on the simulated glands. We simulated different scenarios in which only low-curvature regions or only high-curvature regions were selectively upregulated (**Supplementary Figs. 18 and 19**). We observed that the sign of the correlation depends on whether high- or low-curvature regions are upregulated. Importantly, we emphasize that this reflects a general trend rather than a strict linear relationship.

Our findings demonstrate the utility of CurvSeq/CurvSee as a multimodal framework for linking glandular morphology to molecular dynamics in prostate cancer. In our spatial transcriptomics datasets, we identified genes whose expression was associated with gland morphology and local microenvironmental features. These gene groups were associated with distinct biological functions, suggesting that differences in gland shape may be linked to specific molecular programs. In both the Xenium and Visium datasets, glands showed substantial inter-gland heterogeneity in their morphology–expression relationships. Genes enriched in pocket regions of one gland were not necessarily enriched in morphologically similar pockets of another gland, indicating that pocket-associated expression patterns were highly context-dependent. These results suggest that individual glands can exhibit distinct, context-dependent molecular patterns, with local gene-expression programs associated with gland-specific architecture and microenvironmental composition rather than by morphology alone. Across both spatial transcriptomics datasets, benign glands frequently displayed complex folded architectures. In the Xenium dataset, GG1 glands were generally smaller and more circular than benign glands, whereas in the Visium dataset, GG4 glands formed larger, densely cellular, and relatively circular glandular structures (**Fig. 4a; Supplementary Fig. 4**).

In the Xenium dataset, CCN1 showed notable pocket-associated heterogeneity. In some glands, such as Gland 22, CCN1 expression was elevated across multiple pocket regions (Fig. 2f), whereas other glands showed more restricted or variable pocket-level expression. CCN1, also known as CYR61, is a matricellular protein involved in tissue repair, extracellular matrix organization, and modulation of the local microenvironment. CCN1 has also been linked to extracellular matrix remodeling through regulation of matrix metalloproteinases, suggesting that CCN1-high pockets may represent localized remodeling-associated niches within folded glands. Other genes, including RHOC, EGFR, and YAP1, showed increased expression in benign glands and were significantly correlated with the morphomechanical score (Fig. 2e). RHOC is involved in cytoskeletal organization and cell contractility, EGFR is a receptor tyrosine kinase that regulates epithelial growth and signaling, and YAP1 is a mechanotransduction-associated transcriptional regulator involved in sensing tissue stiffness, cell adhesion, and mechanical stress. Together, these genes suggest that benign glands retain mechanosensitive programs linked to gland architecture. Interestingly, in the Visium dataset, glands continued to show substantial gland-to-gland heterogeneity, but MMP7 displayed a reproducible association with gland architecture and microenvironmental context. MMP7 expression was higher in glands that were larger, less circular, and more pocket-rich, and these glands also showed increased immune-cell proximity. Similarly, GG4 glands showed recurring boundary-associated patterns. Glands with neuroendocrine-like epithelial cells enriched along the outer gland layer exhibited increased MMP7 expression and were surrounded by higher immune-cell abundance.

Furthermore, we calculated the curvature–protein expression correlation matrix using an independent spatial proteomics dataset that included both Gleason-graded and TAN tissues. When projected into a latent space with UMAP, the resulting profiles separated glands from TAN tissues and those from Gleason-graded samples. This separation suggests that using only protein-curvature correlations, it may be possible to distinguish TAN from Gleason-graded glands, indicating a potential link between cellular mechanics, morphological properties, and underlying protein regulation. Using a random forest classifier, we predicted the tissue origin of each gland with 74% accuracy, highlighting the relationships between curvature, morphology, and glandular protein expression. The moderate classification accuracy likely reflects that this analysis used only curvature–protein expression correlations and did not incorporate additional morphologic or microenvironmental features, pocket architecture, stromal composition, or immune-cell proximity, which may further distinguish TAN and Gleason-graded glands. Proteins linked to mechanical functions appear prominently among the most important features. Tryptase has been implicated in the degradation of extracellular-matrix and basement-membrane proteins ^36^. Vimentin is an intermediate filament protein that regulates cytoskeletal organization, cell stiffness, and deformability ^37^. *Ecad* is a cell-cell adhesion molecule. In cell pairs, *Ecad* junctions maintain a relatively constant molecular tension and regulate force balance, linking adhesion to mechanical load distribution ^38^. These results show that curvature-associated protein organization alone captures meaningful differences between TAN and Gleason-graded glands, even without incorporating additional morphologic or microenvironmental features (**Fig. 8c)**.

Finally, we compared curvature–gene correlations in cohorts with *FOXA1* knockout. In these tissues, the glands exhibited disrupted morphology, characterized by reduced circularity. *FOXA1* has been shown to control the expression of cell adhesion and cytoskeletal genes. Loss of FOXA1 leads to changes in epithelial polarity and ductal morphology ^39^. Moreover, *SMAD* signaling is well known to drive ECM deposition, fibrosis, and stiffening in many tissues ^40^. Together, these findings suggest that transcriptional regulation of cell identity and extracellular signaling may contribute to the mechanical organization of glandular tissues. Our results indicate that FOXA1 loss is associated with altered coupling between tissue architecture, signaling activity, and local mechanical behavior in the prostate gland. In control glands, FOXA1 expression coincided with more organized glandular morphologies and coordinated pSMAD2 activity, consistent with a structured relationship between curvature, stiffness, and transcriptional regulation. In contrast, FOXA1-knockout tissues showed more irregular gland morphology, reduced circularity, and increased curvature, suggesting a disruption of glandular architecture. These morphological changes were accompanied by altered curvature–stiffness relationships. In fresh control tissues, high-curvature gland regions tended to show higher stiffness, whereas FOXA1-knockout glands showed increased stiffness in lower-curvature regions. Furthermore, whole-region stiffness maps revealed differences in gland contour organization between control and Foxa1-knockout tissues (**Fig. 8d**). This shift suggests that FOXA1 loss may alter how glandular tissues mechanically organize across regions of different curvature. While these data do not establish a direct causal mechanism, they are consistent with a role for FOXA1 in coordinating epithelial transcriptional state, tissue morphology, and local mechanical properties. Such disruption may provide one route by which altered transcriptional programs are associated with disorganized gland architecture in this tumor model.

Several limitations must be brought up. First, the overall sample size is modest, using public datasets and small patient cohorts. Larger-scale validation is required to confirm generalizability. Second, the gland segmentation was performed manually, which introduces subjectivity and bias. Future improvements in gland segmentation models could further reduce subjectivity in the pipeline by providing more consistent and precise gland boundary annotations. More accurate automated segmentation would improve contour extraction, curvature estimation, and pocket detection, thereby strengthening the reproducibility of downstream morphomechanical analyses. Third, our results show moderate to high correlation between gene expression and glandular morphology, but this does not prove causality. Gland folding may result from several processes, including differential epithelial growth, growth-induced buckling, stromal confinement, smooth-muscle contraction, extracellular-matrix remodeling, or the geometry of two-dimensional tissue sectioning. Curvature should therefore be interpreted as a geometric feature associated with tissue organization rather than as a direct measurement of mechanical stress or force. Further experiments are needed to validate the link between curvature and molecular changes. Finally, differences in spatial resolution across various datasets, including Xenium and Visium, as well as in lab immunofluorescence staining, may affect sensitivity in detecting patterns. Applying CurvSee to a larger cohort of patients would make it more robust. Extending this analysis to more comprehensively map mechanical stiffness across larger tissue areas would provide a deeper understanding of how mechanical properties vary throughout the gland. Since our current measurements were limited to selected regions, broader scanning would enable stronger connections between local stiffness patterns, gene and protein expression, and overall glandular morphology.

From a modeling perspective, CurvSeq computes a morphomechanical score analogous to pseudotime analysis, but using gland morphology and microenvironmental composition rather than transcriptional features.

In this study, interpretability was prioritized by using linear regression to relate glandular epithelial gene expression to nearby morphologic and microenvironmental features. However, these relationships are likely nonlinear and context-dependent, as shown by the strong gland-to-gland heterogeneity observed across datasets. Importantly, the associations reported here are correlative and do not establish direct causal relationships between gland morphology, microenvironmental composition, and gene expression. Future experimental validation, including perturbation studies and direct mechanical measurements, will be needed to determine whether specific morphologic or microenvironmental features actively drive the observed molecular programs. Future models that explicitly account for spatial structure may better capture how epithelial cells respond to local gland geometry and neighboring stromal or immune cells. For example, morphology-aware graph neural networks could model epithelial cells, gland boundaries, pockets, and surrounding cell types as connected spatial features, potentially improving prediction of gene-expression patterns while preserving the local tissue organization.

Importantly, this framework is not limited to prostate cancer. Applying CurvSeq to other glandular malignancies, including colon, pancreatic, and breast cancer, could determine whether curvature–gene expression coupling and pocket-associated microenvironmental niches are conserved across epithelial tumors. Extending the approach to three-dimensional imaging would also allow curvature and gland architecture to be measured in volumetric reconstructions, which may better reflect in vivo tissue stresses than two-dimensional sections. In future work, physics-informed neural networks and related geometric deep-learning approaches could further integrate curvature-derived physical constraints with spatial molecular data, enabling more mechanistic models of how tissue mechanics, morphology, and molecular state interact during cancer progression.

## Supporting information

Spatial Curve Seq Supplementary Materials

## Methods

### Dataset 1: FFPE Human Prostate Adenocarcinoma with 5K Human Pan Tissue and Pathways Panel

The first dataset is a spatial transcriptomics dataset produced using the 10x Genomics platform, containing 5,000 genes. Biomaterial specifications and tissue-preparation protocols are documented in the Xenium Prime FFPE Human Prostate dataset: https://www.10xgenomics.com/datasets/xenium-prime-ffpe-human-prostate

### Dataset 2: Visium HD Spatial Gene Expression Library, Human Prostate Cancer (FFPE)

The second dataset is also a spatial transcriptomics dataset, originating from the 10x Genomics platform, and contains over 18,000 genes. Biomaterial specifications and tissue-preparation protocols are documented here.: https://www.10xgenomics.com/datasets/visium-hd-cytassist-gene-expression-libraries-human-prostate-cancer-ffpe

### Dataset 3: Fluorescent Immunolabeled FFPE prostate sections

The third dataset is a multiplex immunofluorescence (mIF) spatial proteomics dataset generated from FFPE prostate cancer sections in our laboratory. The tissues were stained for: *AMACR/P504S, P63, PSA, CK HMW, H3ka9Ac, H3k4me2, H3k27me3, H4k12Ac, Concanavalin A, Phalloidins, WGA*. Slides were dewaxed and rehydrated, antigens were retrieved in sodium⍰citrate buffer (pH 6) using a pressure cooker (15 min), and tissues were blocked for 30 min at room temperature. Primary antibodies were incubated overnight at 4 °C, followed by species⍰matched secondary antibodies (e.g., Alexa Fluor dyes, 1:1000, 60 min, RT) and Hoechst nuclear counterstain before imaging. For iterative cycles, fluorophores were chemically bleached (4.5% H_2_O_2_/24 mM NaOH, 1 h, RT) and, when required, antibodies were stripped in a β⍰mercaptoethanol/SDS/Tris⍰HCl buffer at 56 °C for 30 min, after which slides were re-blocked and re-stained. Directly conjugated primaries (e.g., PSA, ERG, NKX3.1, H3K4me3) were applied without a secondary step (**Supplementary Table 1a**). Clinical details of the patients are summarized in the supplementary figures (**Supplementary Table 1b**).

### Dataset 4: Fluorescent Immunolabeled FFPE prostate of TAN and Tumor sections

The fourth dataset is a previously published spatial proteomics dataset of prostate cancer ^41^. The proteins included in the dataset are:

*AR, ChromA, CK5, Ecad, NCAM, Vim, NKX3*.*1, CK8, AMACR, H3K27ac, CD44, H3K4me3, Tryp, CD68, GZMB, Ki67, CD20, HLADRB1, CD3, CD11b, CD4, CD45, CD163, CD66b, PD1, CD90, FOXP3, and aSMA*.

The dataset was obtained directly from the supplementary materials of the original publication.

### Dataset 5: FOXA1 Knockout dataset

The fifth dataset is a spatial proteomics dataset that was stained in our lab. Mouse prostate tissues were obtained from Pten-null (P) and Pten/Foxa1 double-knockout (PF) mice at the indicated timepoints. The morphological characteristics of these models have been described previously ^42^. Formalin-fixed paraffin-embedded (FFPE), 10 µm thick, prostate cancer tissue sections were analyzed from samples in which the FOXA1 gene was genetically knocked out (PF) alongside controls retaining FOXA1 expression (P). Sections were subjected to antigen retrieval using citrate buffer. After retrieval, tissues were blocked with 5% bovine serum albumin (BSA).

### Dataset 6: PAVONE measurements

The sixth dataset comprised prostate tissue sections from P12, PF12, P19, and PF19 samples. During nanoindentation measurements, the tissue sections were maintained in phosphate-buffered saline (PBS) to preserve tissue hydration and minimize sample deterioration.

### Image Preprocessing

#### Dataset□1

No additional image processing or registration was required; cell segmentation and transcript quantification were provided by 10x□Genomics.

#### Dataset 2

No additional image processing or registration was required; cell segmentation and transcript quantification were provided by 10x□Genomics.

#### Dataset□3

Inter⍰cycle registration was performed using the Hoechst (nuclear) channel as the reference. We estimated transforms with FIJI/ImageJ’s Linear Stack Alignment with the SIFT plugin. We applied the resulting warp matrices—computed between nuclei images of successive cycles—to the corresponding fluorescence channels in each cycle.

#### Dataset□4

No further image processing or registration was performed, as the original authors already completed these steps.

#### Dataset 5

Inter⍰cycle registration was performed using the DAPI channel from the first cycle as a reference. The estimation of the transforms and the subsequent warping is the same as for Dataset 3. Regions with a high abundance of non-specific staining were cropped out of the images. Nuclear segmentation was done using Cellpose (Model: Cyto3, radius = 25). Cells were filtered based on the DAPI signal first. Cells that have none or very low DAPI signal were removed. Afterward, cells that had a very high signal for FOXA1 and pSMAD2 were considered outliers and were removed from the dataset.

### Segmentation of glandular structures

Gland masks were generated by manual annotation across all datasets to ensure accurate delineation of glandular structures. Manual gland annotations were guided by pathologist-provided annotations to maintain consistency with the histological organization of the tissue. All annotated glands underwent visual quality control, and glands with incomplete, ambiguous, or poorly defined boundaries were excluded from downstream analysis. To reduce small contour irregularities introduced by hand movement during annotation, the extracted gland boundaries were smoothed before curvature and pocket analysis. The final annotations were converted into instance masks in which each gland was assigned a unique integer label for downstream gland-specific analysis.

### Identification of Pocket regions

Pocket regions were identified from the segmented gland boundaries by comparing each segmented gland contour with its corresponding alpha-shape envelope. An alpha parameter of 0.001 was used to generate the alpha shape, which captured the overall gland outline while preserving the major boundary structure. Candidate pocket regions were defined as the geometric difference between the alpha-shape envelope and the original segmented gland contour, thereby identifying inward concave regions where the segmented boundary deviated from the overall gland shape. For each candidate pocket, we measured the pocket depth, defined as the maximum inward distance from the pocket opening to the deepest point of the concavity, and the pocket mouth length, defined as the distance across the opening of the pocket. To exclude broad and shallow indentations, candidate pockets with a depth-to-mouth-length ratio below 0.3 were removed. The remaining connected pocket regions were labeled individually and used for downstream analysis (**Supplementary Fig. 20**).

### Computation of the morphomechanical score

To summarize gland architecture along a continuous morphomechanical axis, we computed a gland-level morphomechanical score from geometric, curvature-based, pocket-associated, basal-layer, and peri-glandular microenvironmental features. For each segmented gland, we extracted area, perimeter, circularity, aspect ratio, boundary curvature variance, number of pockets, fraction of gland area occupied by pockets, mean pocket depth, and mean pocket depth-to-mouth-length ratio. Basal-layer organization was quantified from basal epithelial cells located inside each gland by measuring the shortest distance from each basal cell centroid to the gland boundary and summarizing these distances at the gland level. The peri-glandular microenvironment was quantified by expanding each gland boundary outward by a fixed buffer distance and measuring cell-type-specific densities within the resulting external band. A gland-level AnnData object was generated from the standardized morphomechanical feature matrix using Scanpy. Features were first scaled using sc.pp.scale, and a nearest-neighbor graph was constructed with sc.pp.neighbors using n_neighbors=5. Diffusion map embedding was then computed with sc.tl.diffmap. Diffusion pseudotime was then computed using sc.tl.dpt, and the resulting dpt_pseudotime values were used as the continuous morphomechanical score. This score was interpreted as a relative ordering of gland morphomechanical state rather than as a temporal lineage measurement

### Curvature Analysis

After segmenting the glands, we obtain a set of points representing the gland boundaries using **OpenCV**’s function ***findContours***, which are then used for curvature computation. Various papers have proposed different methods to compute curvature ^43^. One common approach involves selecting three neighboring points on the contour and computing the curvature by fitting a circle through them, using the circumradius to approximate local curvature ^44^ (**Supplementary Fig. 21a**). In this paper, we adopt a different approach by fitting a polynomial to a set of adjacent points and estimating the curvature from its first and second derivatives.

Let **Γ** be a closed contour consisting of N discrete points represented as an ordered sequence of points ***r*_*i*_** = (*x*_*i*_,*y*_*i*_) *for i* ∈{0,1,…,*N* −1}The points are arranged such that each point ***r*_*i*_** is adjacent to ***r*_*i*−1_** and ***r*_*i*+1_**, following the ordering:

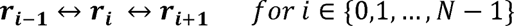

We define a local window centered at index **I** with a predefined window size **w**. The set of neighborhood points is indexed as:

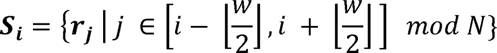

The modular arithmetic ensures that the window wraps around the contour for points near the endpoints. The local tangent direction at the central point *r*_*i*_ is approximated by

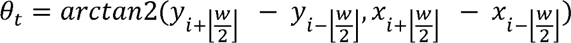

To simplify the curvature estimation and to ensure that the curvature is computed relative to the local tangent, the local neighborhood ***S*_*i*_** is translated such that ***r*_*i*_** becomes the origin, and then rotated by − *θ*_*t*_ to align the local x-axis with the tangent direction. The transformation is given by:

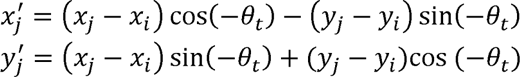

A third-degree polynomial p(x) is fitted to transformed neighborhood points 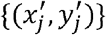,

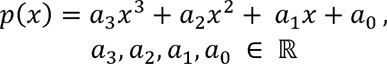

The first and second derivatives are given by:

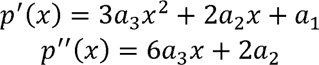

Evaluating these at x = 0, corresponding to the central point, we obtain (**Supplementary Fig. 21b**):

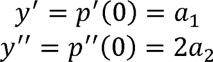

Using the standard curvature formula for a function y = f(x) ^45^,

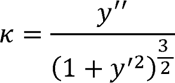

The curvature at the central point is computed as:

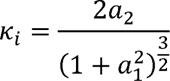

The radius of the best-fitting circle at the central point I is given by the inverse of the curvature:

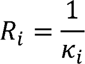

We keep the signed curvature value *K*_*i*_ because it shows whether the curvature bends inward or outward. The final curvature value is scaled by a factor of 100 to adjust for numerical scaling (**Supplementary Fig. 21c)**.

After computing the curvature of the contours, we aim to map these curvature values onto the cells. The projected values will indicate whether a cell is located in a region of the gland with high or low curvature and whether the curvature is directed inward or outward.

The glands overlayed with curvature values can be found in the supplementary figures (**Supplementary Figs. 22-26**)

### Projecting curvature value onto the cells

Let **Γ** be a closed contour consisting of N discrete points represented as an ordered sequence of points ***r*_*i*_** = (*x*_*i*_,*y*_*i*_) *for i* = 0,1,…,*N* −1 *and* ***c*_*j*_** = (*x*_*cj*_,*y*_*cj*_) *for j* = 0,1…,*M* with M being the number of cells in the dataset. We use **OpenCV**’s function ***pointPolygonTest*** to detect if a point ***c*_*j*_** is located inside of the contour **Γ**. We will note the subset of cells belonging inside of the contour **Γ** cell ***c*^Γ^**. Let 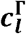 be a cell inside of **Γ**. We compute the Euclidean norm between 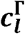 and all ***r*_*i*_**

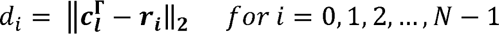

We keep the first k closest contour points to the cell, where k is a user-defined hyperparameter, and average their curvature values to get the projected curvature value in cell 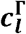 Depending on the number of points that constitute the boundary of the glands, the parameter k can be changed (**Supplementary Fig. 27**).

A summary of the total number of glands analyzed in each dataset is provided **(Supplementary Table 2). Cell clustering based on curvature**

Using the projected curvature as a per⍰cell feature, we first binarized cells into positive and negative curvature groups. We then clustered cells jointly by spatial proximity and curvature sign with HDBSCAN (Python; min_cluster_size = 5).

### Random Forest Classification

We implemented a Random Forest Classifier using the Scikit-Learn library to classify TAN versus Gleason-graded glands. The dataset was randomly partitioned into 80% for training and 20% for testing. Hyperparameter tuning was performed with the GridSearchCV function, which identified the following optimal parameters: max_depth = 2, min_samples_leaf = 6, min_samples_split = 10, n_estimators = 25.

#### Mechanical characterization of the prostate tissue sections

The stiffness characterization of the prostate tissues was done by indentation method with Pavone nanoindenter (Optics11 Life). The measurements were done in PBS solution at room temperature. The entire tissue sections were imaged with Pavone’s built-in microscopy, and then the areas of interests were selected to perform matrix scans where points are 20 um apart (dX=20 um and dY=20 um). Indentations were performed with a spherical probe of 3 µm tip radius and stiffness of 0.42 N m-1, which was pre-calibrated by the manufacturer. We used peak-load poking mode with a threshold at 0.5 uN for all the indentations, and the effective Young’s moduli were then calculated from Hertzian contact model and fit up to 0.2 uN of the force curve. All the data reported were selected under the criteria of R-squared value larger than 0.95 from Hertzian contact model fitting.

Local curvature stiffness measurements were conducted using an MFP-3D Atomic Force Microscope (Asylum Research). We first performed indentations on fixed and stained prostate tissues, followed by measurements on freshly prepared samples. For each sample, the AFM probe was positioned along the gland borders to quantify local mechanical stiffness. The AFM generated a stiffness heatmap, where each pixel represents the elastic modulus computed from force–indentation data (**Supplementary Figs. 28a, 28b, 29**). Deflection curves were extracted from each grid point in the heatmap, and the Hertzian contact model was manually fitted by identifying the contact point on each curve (**Supplementary Fig. 28c**). Curves with poor or ambiguous contact points were excluded, and an additional filtering step was applied based on the goodness of fit (R^2^ value) (**Supplementary Fig. 28d**). The images of the cantilever placement for measurement on the brightfield images of the glands can be found in the supplementary figures (**Supplementary Figs. 30, 31**).

For fresh tissues, measurements were collected over a smaller 25 µm × 25 µm region using a 4 × 4 grid, with cantilever stiffnesses of 80 × 10^−3^ N/m for PF12 and 21 × 10^−3^ N/m for P12. Fits showing R^2^ > 0.9 were retained for analysis.

## Data availability

All data generated or analyzed in this study are available from the corresponding author upon request.

## Code availability

The CurvSeq analysis code is available at https://github.com/coskunlab/spatialcurvseq.

## Acknowledgements

A.F.C. holds a Career Award at the Scientific Interface from Burroughs Welcome Fund and Bernie-Marcus Early-Career Professorship. A.F.C. was supported by start-up funds from the Georgia Institute of Technology and Emory University. Research reported in this study was supported by the National Institutes of Health under award number R35GM151028 and R33CA291197. This work was partially supported by the National Science Foundation CAREER under grant number 2338935. F.G.R.M. was supported by the NIH NIGMS T32 Training Program on Cell and Tissue Engineering (T32 GM145735)

## References

1. Siegel, R. L., Giaquinto, A. N. & Jemal, A. Cancer statistics, 2024. CA Cancer J Clin 74, 12–49 (2024).

2. Schrecengost, R. S. & Knudsen, K. E. Molecular Pathogenesis and Progression of Prostate Cancer. Semin Oncol 40, 244–258 (2013).

3. Epstein, J. I. et al. A Contemporary Prostate Cancer Grading System: A Validated Alternative to the Gleason Score. European Urology 69, 428–435 (2016).

4. Saranyutanon, S. et al. Cellular and Molecular Progression of Prostate Cancer: Models for Basic and Preclinical Research. Cancers 12, 2651 (2020).

5. Chen, R. C. et al. Recommended Patient-Reported Core Set of Symptoms to Measure in Prostate Cancer Treatment Trials. JNCI: Journal of the National Cancer Institute 106, dju132 (2014).

6. Haffner, M. C. et al. Genomic and phenotypic heterogeneity in prostate cancer. Nat Rev Urol 18, 79–92 (2021).

7. Golub, T. R. et al. Molecular Classification of Cancer: Class Discovery and Class Prediction by Gene Expression Monitoring. Science 286, 531–537 (1999).

8. Rubin, M. A., Girelli, G. & Demichelis, F. Genomic Correlates to the Newly Proposed Grading Prognostic Groups for Prostate Cancer. European Urology 69, 557–560 (2016).

9. Berglund, E. et al. Spatial maps of prostate cancer transcriptomes reveal an unexplored landscape of heterogeneity. Nat Commun 9, 2419 (2018).

10. Quan, Y., Zhang, H., Wang, M. & Ping, H. Visium spatial transcriptomics reveals intratumor heterogeneity and profiles of Gleason score progression in prostate cancer. iScience 26, 108429 (2023).

11. Xu, J., Wang, D., Di, X. & Liu, Y. Advances in spatial omics for the analysis of prostate cancer. Anal Bioanal Chem 418, 3865–3886 (2026).

12. Chelebian, E. et al. Morphological Features Extracted by AI Associated with Spatial Transcriptomics in Prostate Cancer. Cancers 13, 4837 (2021).

13. Chatterjee, A. et al. Changes in Epithelium, Stroma, and Lumen Space Correlate More Strongly with Gleason Pattern and Are Stronger Predictors of Prostate ADC Changes than Cellularity Metrics. Radiology 277, 751–762 (2015).

14. Maane, I. A. et al. Evaluation of Combined Quantification of PCA3 and AMACR Gene Expression for Molecular Diagnosis of Prostate Cancer in Moroccan Patients by RT-qPCR. Asian Pac J Cancer Prev 17, 5229–5235 (2016).

15. Singh, R. & Kyte, J. A. STEAP1: a promising target in prostate cancer therapy. Trends in Cancer 11, 722–725 (2025).

16. Liang, L. et al. HIF1α-PHD1-FOXA1 Axis Orchestrates Hypoxic Reprogramming and Androgen Signaling Suppression in Prostate Cancer. Cells 14, 1008 (2025).

17. Erokhin, M. M., Kozelchuk, N. Y., Ziganshin, R. H., Tatarskiy, V. V. & Chetverina, D. A. HOXB13 interactome in prostate cancer cells: biochemical and functional interactions between the transcription factors HOXB13 and TBX3. Vavilovskii Zhurnal Genet Selektsii 29, 744–752 (2025).

18. Weiner, A. B. et al. Molecular hallmarks of prostate-specific membrane antigen in treatment-naïve prostate cancer. Eur Urol 86, 579–587 (2024).

19. Thangapazham, R. et al. Loss of the NKX3.1 tumorsuppressor promotes the TMPRSS2-ERG fusion gene expression in prostate cancer. BMC Cancer 14, 16 (2014).

20. Koistinen, H. et al. The roles of proteases in prostate cancer. IUBMB Life 75, 493–513 (2023).

21. Sakamoto, S. et al. Increased expression of CYR61, an extracellular matrix signaling protein, in human benign prostatic hyperplasia and its regulation by lysophosphatidic acid. Endocrinology 145, 2929–2940 (2004).

22. Lynch, C. C. et al. MMP-7 promotes prostate cancer-induced osteolysis via the solubilization of RANKL. Cancer Cell 7, 485–496 (2005).

23. Khorshid Sokhangouy, S., Zeinali, M., Fathi, S. & Nazari, E. Deep learning assisted identification of SCUBE2 and SLC16A5 combination in RNA-sequencing data as a novel specific potential diagnostic biomarker in prostate cancer. Med Biol Eng Comput 63, 2955–2968 (2025).

24. Song, Q. et al. Decreased expression of SCUBE2 is associated with progression and prognosis in colorectal cancer. Oncol Rep 33, 1956–1964 (2015).

25. Taverna, G. et al. Mast Cells as a Potential Prognostic Marker in Prostate Cancer. Dis Markers 35, 711–720 (2013).

26. Li, R., Wheeler, T., Dai, H. & Ayala, G. Neural cell adhesion molecule is upregulated in nerves with prostate cancer invasion. Human Pathology 34, 457–461 (2003).

27. Romero, R. et al. The neuroendocrine transition in prostate cancer is dynamic and dependent on ASCL1. Nat Cancer 5, 1641–1659 (2024).

28. Song, B. et al. Targeting FOXA1-mediated repression of TGF-β signaling suppresses castration-resistant prostate cancer progression. J Clin Invest 129, 569–582 (2019).

29. Chen, C., Kim, H. J. & Yang, P. Evaluating spatially variable gene detection methods for spatial transcriptomics data. Genome Biology 25, 18 (2024).

30. Behanova, A., Klemm, A. & Wählby, C. Spatial Statistics for Understanding Tissue Organization. Front Physiol 13, 832417 (2022).

31. Muhammad, A. Curvature-Aware Deep Learning for Vector Boson Fusion: Differential Geometry, Physics-Inspired Features, and Quantum Method Limitations. Preprint at 10.48550/arXiv.2510.04887 (2025).

32. Burbulla, S. Physics-informed neural networks for transformed geometries and manifolds. Preprint at 10.48550/arXiv.2311.15940 (2023).

33. Kim, J. A Review of Continuum Mechanics for Mechanical Deformation of Lipid Membranes. Membranes (Basel) 13, 493 (2023).

34. Du, C.-J., Hawkins, P. T., Stephens, L. R. & Bretschneider, T. 3D time series analysis of cell shape using Laplacian approaches. BMC Bioinformatics 14, 296 (2013).

35. Rangamani, P., Behzadan, A. & Holst, M. Local sensitivity analysis of the “membrane shape equation” derived from the Helfrich energy. Mathematics and Mechanics of Solids 26, 356–385 (2021).

36. Iddamalgoda, A. et al. Mast cell tryptase and photoaging: possible involvement in the degradation of extracellular matrix and basement membrane proteins. Arch Dermatol Res 300 Suppl 1, S69–76 (2008).

37. Alisafaei, F. et al. Vimentin is a key regulator of cell mechanosensing through opposite actions on actomyosin and microtubule networks. Commun Biol 7, 658 (2024).

38. Sim, J. Y. et al. Spatial distribution of cell-cell and cell-ECM adhesions regulates force balance while maintaining E-cadherin molecular tension in cell pairs. Mol Biol Cell 26, 2456–2465 (2015).

39. De Felice, D. et al. Rarγ-Foxa1 signaling promotes luminal identity in prostate progenitors and is disrupted in prostate cancer. EMBO Rep 26, 443–469 (2025).

40. Massagué, J. & Sheppard, D. TGF-β signaling in health and disease. Cell 186, 4007–4037 (2023).

41. Ak, Ç. et al. Multiplex imaging of localized prostate tumors reveals altered spatial organization of AR-positive cells in the microenvironment. iScience 27, 110668 (2024).

42. Brea, L. et al. FOXA1 loss drives basal/squamous de-differentiation of prostate cancer and induces an immunosuppressive tumor microenvironment. Nat Commun 17, 4572 (2026).

43. Lin, W.-Y., Chiu, Y.-L., Widder, K., Hu, Y. & Boston, N. Robust and Accurate Curvature Estimation Using Adaptive Line Integrals. EURASIP J. Adv. Signal Process. 2010, 240309 (2010).

44. Driscoll, M. K. et al. Cell Shape Dynamics: From Waves to Migration. PLoS Comput Biol 8, e1002392 (2012).

45. Chrystie, R. S. M., Burns, I. S., Hult, J. & Kaminski, C. F. On the improvement of two-dimensional curvature computation and its application to turbulent premixed flame correlations. Meas. Sci. Technol. 19, 125503 (2008).

