## Supplementary material for "Morphomechanically-Informed Spatial Curvature Sequencing in Prostate Cancer": Spatial Curve Seq Supplementary Materials

### Supplementary Materials for Morphomechanically-Informed Spatial Curvature Sequencing in Prostate Cancer

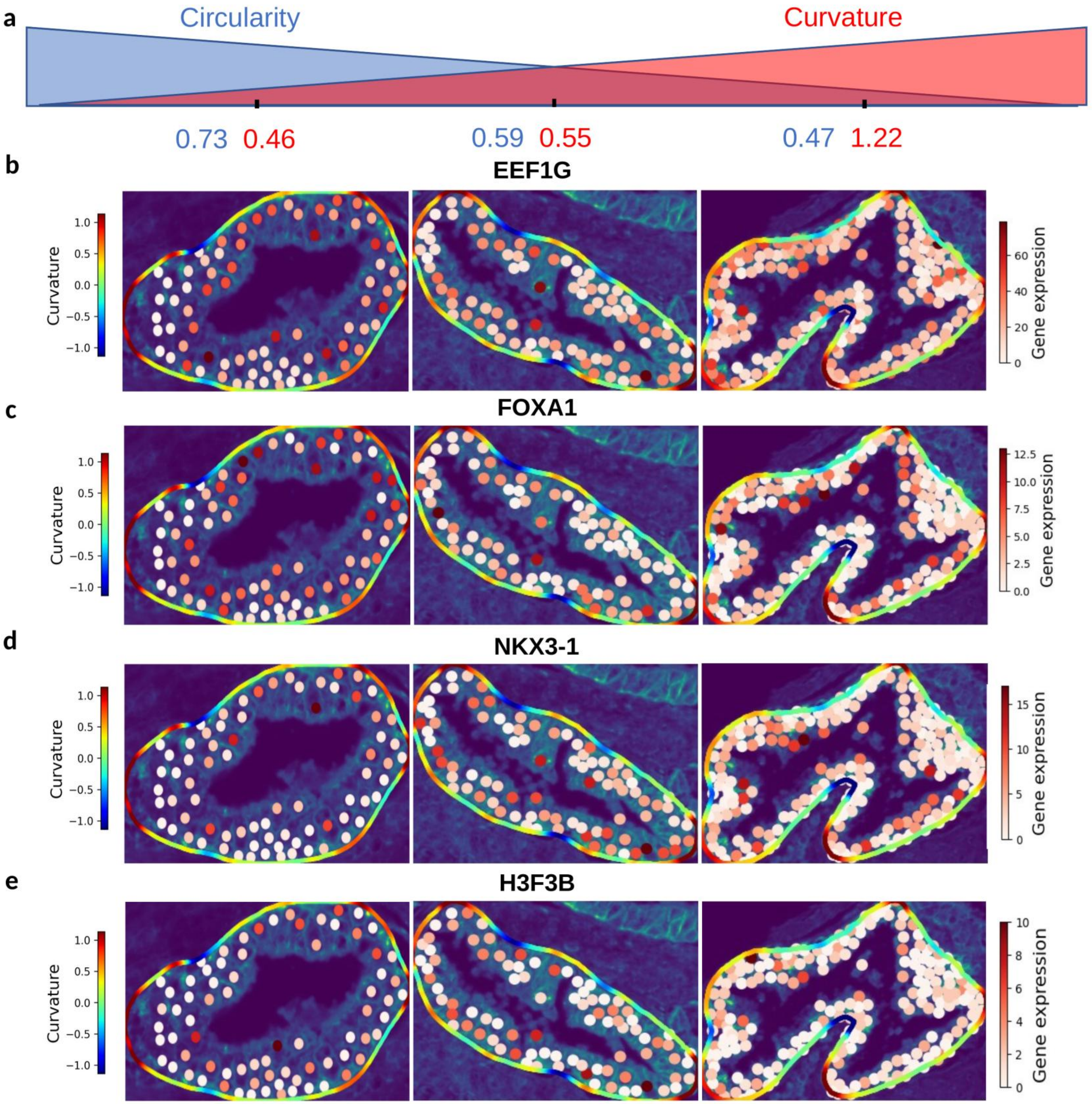

a
Circularity
Curvature
0.73 0.46
0.59 0.55
0.47 1.22
b
EEF1G
c
FOXA1
d
NKX3-1
e
H3F3B
Curvature
Gene expression

**Supplementary Figure 1. Representative curvature-associated epithelial gene-expression patterns across glands with increasing morphologic complexity.**
(a) Schematic representation of the gland morphology axis used to select representative glands, ranging from higher circularity to higher curvature. Numerical values indicate circularity and curvature measurements for the selected examples. (b–e) Representative glands showing spatial expression of EEF1G, FOXA1, NKX3-1, and H3F3B overlaid with gland contours and projected curvature values. White points denote epithelial cells within each gland, and the contour color represents local boundary curvature. Across selected glands, gene expression is visualized relative to changes in gland shape, circularity, and curvature, illustrating heterogeneous spatial expression patterns along the gland boundary and within epithelial regions.

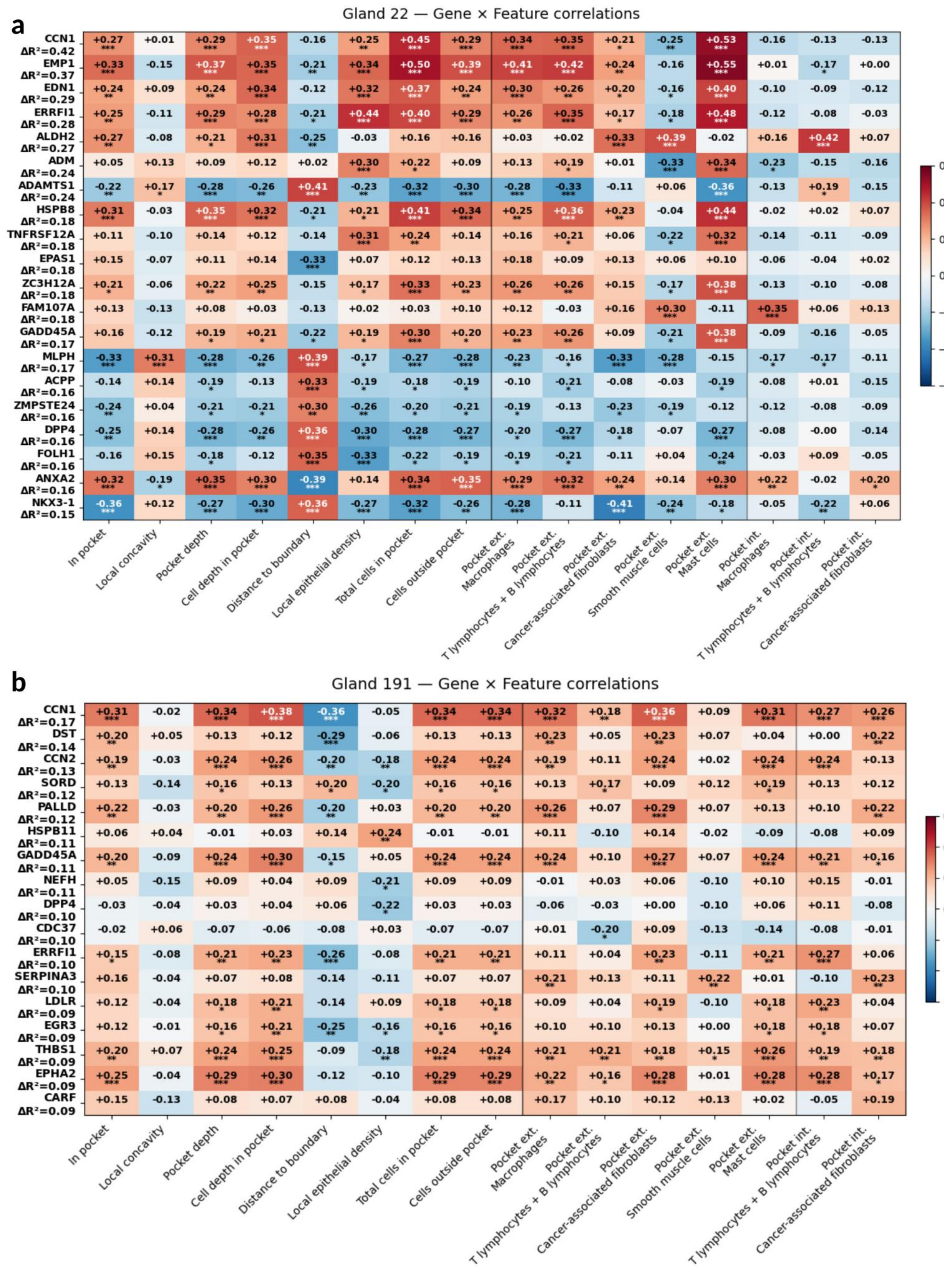
a
Gland 22 — Gene × Feature correlations
CCN1 ΔR²=0.42
EMP1 ΔR²=0.37
EDN1 ΔR²=0.29
ERRFI1 ΔR²=0.28
ALDH2 ΔR²=0.27
ADM ΔR²=0.24
ADAMTS1 ΔR²=0.24
HSPB8 ΔR²=0.18
TNFRSF12A ΔR²=0.18
EPAS1 ΔR²=0.18
ZC3H12A ΔR²=0.18
FAM107A ΔR²=0.18
GADD45A ΔR²=0.17
MLPH ΔR²=0.17
ACPP ΔR²=0.16
ZMPSTE24 ΔR²=0.16
DPP4 ΔR²=0.16
FOLH1 ΔR²=0.16
ANXA2 ΔR²=0.16
NKX3-1 ΔR²=0.15
In pocket
Local concavity
Pocket depth
Cell depth in pocket
Distance to boundary
Local epithelial density
Total cells in pocket
Cells outside pocket
Pocket ext. Macrophages
Pocket ext. T lymphocytes + B lymphocytes
Pocket ext. Cancer-associated fibroblasts
Pocket ext. Smooth muscle cells
Pocket ext. Mast cells
Pocket int. Macrophages
Pocket int. T lymphocytes + B lymphocytes
Pocket int. Cancer-associated fibroblasts
Spearman ρ
0.6
0.4
0.2
0.0
−0.2
−0.4
−0.6
b
Gland 191 — Gene × Feature correlations
CCN1 ΔR²=0.17
DST ΔR²=0.14
CCN2 ΔR²=0.13
SORD ΔR²=0.12
PALLD ΔR²=0.12
HSPB11 ΔR²=0.11
GADD45A ΔR²=0.11
NEFH ΔR²=0.11
DPP4 ΔR²=0.10
CDC37 ΔR²=0.10
ERRFI1 ΔR²=0.10
SERPINA3 ΔR²=0.10
LDLR ΔR²=0.09
EGR3 ΔR²=0.09
THBS1 ΔR²=0.09
EPHA2 ΔR²=0.09
CARF ΔR²=0.09
In pocket
Local concavity
Pocket depth
Cell depth in pocket
Distance to boundary
Local epithelial density
Total cells in pocket
Cells outside pocket
Pocket ext. Macrophages
Pocket ext. T lymphocytes + B lymphocytes
Pocket ext. Cancer-associated fibroblasts
Pocket ext. Smooth muscle cells
Pocket ext. Mast cells
Pocket int. T lymphocytes + B lymphocytes
Pocket int. Cancer-associated fibroblasts
Spearman ρ
0.6
0.4
0.2
0.0
−0.2
−0.4
−0.6

**Supplementary Figure 2. Gene–feature correlation analysis reveals gland-specific associations between epithelial gene expression, pocket architecture, and local microenvironmental composition.**

(a) Heatmap of Spearman correlations between epithelial gene expression and morphologic or microenvironmental features in gland 22. Rows correspond to selected genes and columns correspond to pocket-level or local gland features, including pocket location, local concavity, pocket depth, cell depth within pockets, distance to boundary, epithelial density, pocket size, stromal composition, smooth muscle, fibroblasts, macrophages, lymphocytes, and cancer-associated fibroblast features. (b) Heatmap of Spearman correlations between genes and feature classes in gland 191. Correlation coefficients are shown within each heatmap cell, with red indicating positive associations and blue indicating negative associations. Asterisks indicate statistical significance.

a

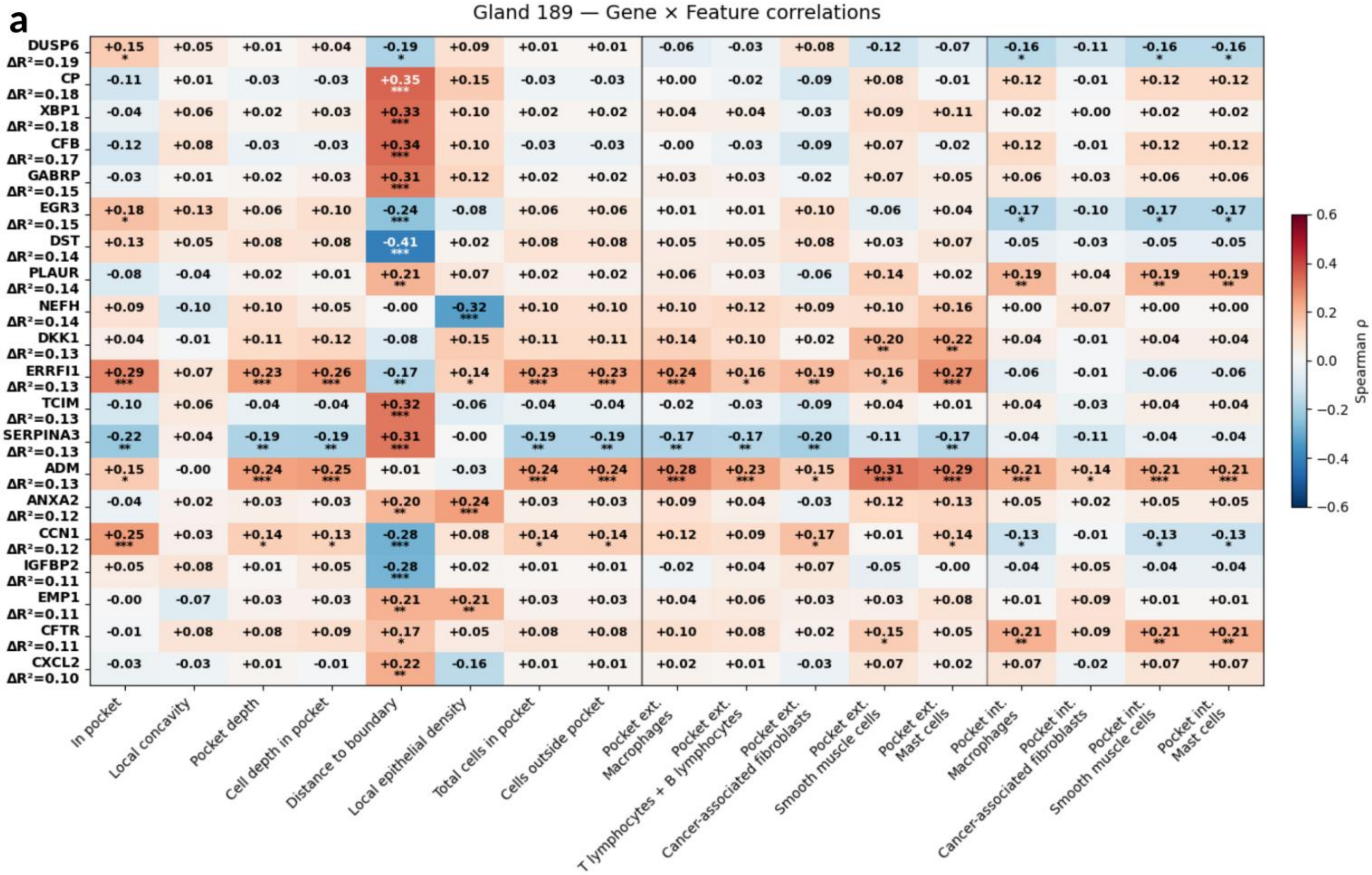

Gland 189 — Gene × Feature correlations
DUSP6 ΔR²=0.19
CP ΔR²=0.18
XBP1 ΔR²=0.18
CFB ΔR²=0.17
GABRP ΔR²=0.15
EGR3 ΔR²=0.15
DST ΔR²=0.14
PLAUR ΔR²=0.14
NEFH ΔR²=0.14
DKK1 ΔR²=0.13
ERRFI1 ΔR²=0.13
TCIM ΔR²=0.13
SERPINA3 ΔR²=0.13
ADM ΔR²=0.13
ANXA2 ΔR²=0.12
CCN1 ΔR²=0.12
IGFBP2 ΔR²=0.11
EMP1 ΔR²=0.11
CFTR ΔR²=0.11
CXCL2 ΔR²=0.10
In pocket
Local concavity
Pocket depth
Cell depth in pocket
Distance to boundary
Local epithelial density
Total cells in pocket
Cells outside pocket
Pocket ext. Macrophages
Pocket ext. T lymphocytes + B lymphocytes
Pocket ext. Cancer-associated fibroblasts
Pocket ext. Smooth muscle cells
Pocket ext. Mast cells
Pocket int. Macrophages
Pocket int. Cancer-associated fibroblasts
Pocket int. Smooth muscle cells
Pocket int. Mast cells
Spearman ρ
0.6
0.4
0.2
0.0
−0.2
−0.4
−0.6


b

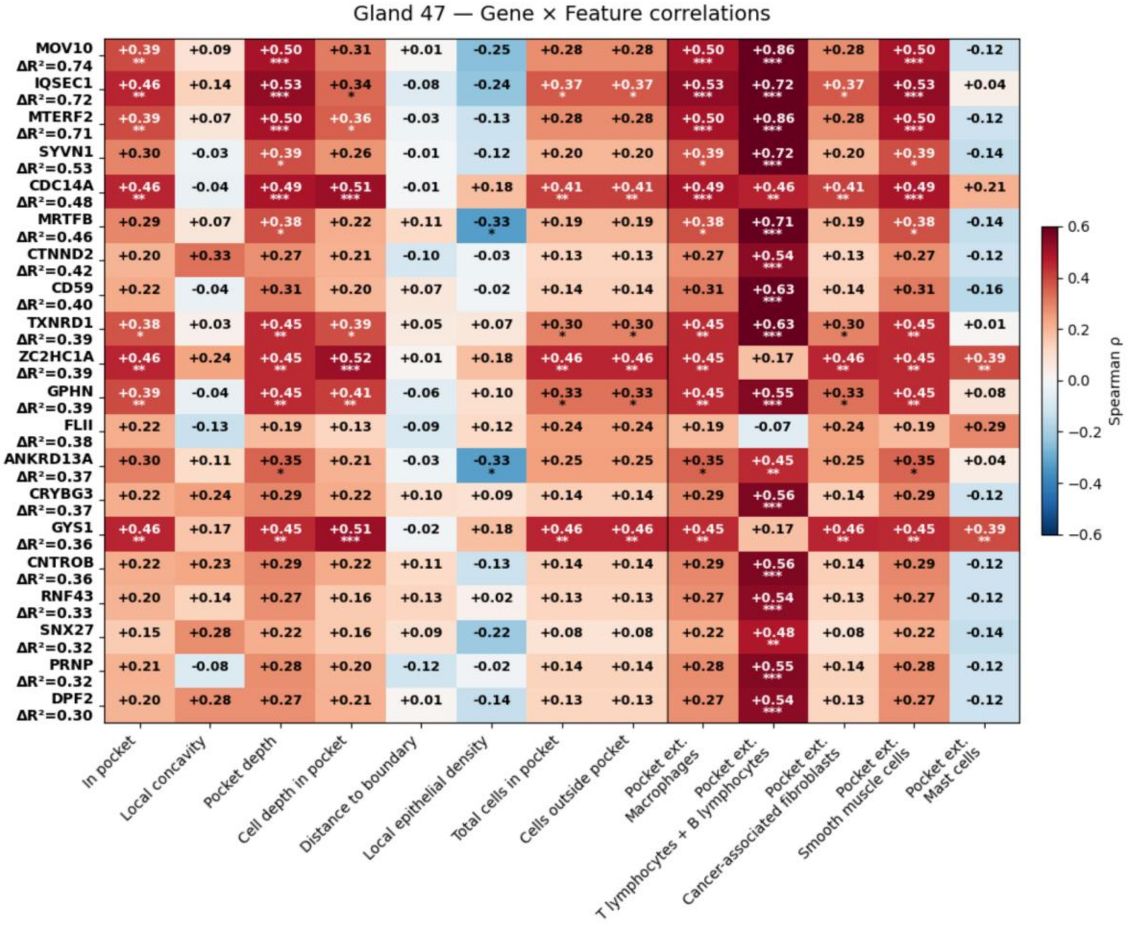

Gland 47 — Gene × Feature correlations
MOV10 ΔR²=0.74
IQSEC1 ΔR²=0.72
MTERF2 ΔR²=0.71
SYVN1 ΔR²=0.53
CDC14A ΔR²=0.48
MRTFB ΔR²=0.46
CTNND2 ΔR²=0.42
CD59 ΔR²=0.40
TXNRD1 ΔR²=0.39
ZC2HC1A ΔR²=0.39
GPHN ΔR²=0.39
FLII ΔR²=0.38
ANKRD13A ΔR²=0.37
CRYBG3 ΔR²=0.37
GYS1 ΔR²=0.36
CNTROB ΔR²=0.36
RNF43 ΔR²=0.33
SNX27 ΔR²=0.32
PRNP ΔR²=0.32
DPF2 ΔR²=0.30
In pocket
Local concavity
Pocket depth
Cell depth in pocket
Distance to boundary
Local epithelial density
Total cells in pocket
Cells outside pocket
Pocket ext. Macrophages
Pocket ext. T lymphocytes + B lymphocytes
Pocket ext. Cancer-associated fibroblasts
Pocket ext. Smooth muscle cells
Pocket ext. Mast cells
Spearman ρ
0.6
0.4
0.2
0.0
−0.2
−0.4
−0.6

**Supplementary Figure 3. Additional gland-level gene–feature correlations demonstrate heterogeneous pocket-associated transcriptional programs across individual glands.** (a) Heatmap of Spearman correlations between epithelial gene expression and pocket-level morphologic or microenvironmental features in gland 189. Features include pocket localization, local concavity, pocket depth, cell depth within pockets, distance to boundary, epithelial density, pocket size, nearby stromal and immune cell densities, and cell-type composition outside pocket regions. (b) Heatmap of Spearman correlations between epithelial gene expression and the same feature classes in gland 47. Correlation coefficients are displayed within each cell, with red indicating positive associations and blue indicating negative associations. Asterisks indicate statistical significance.

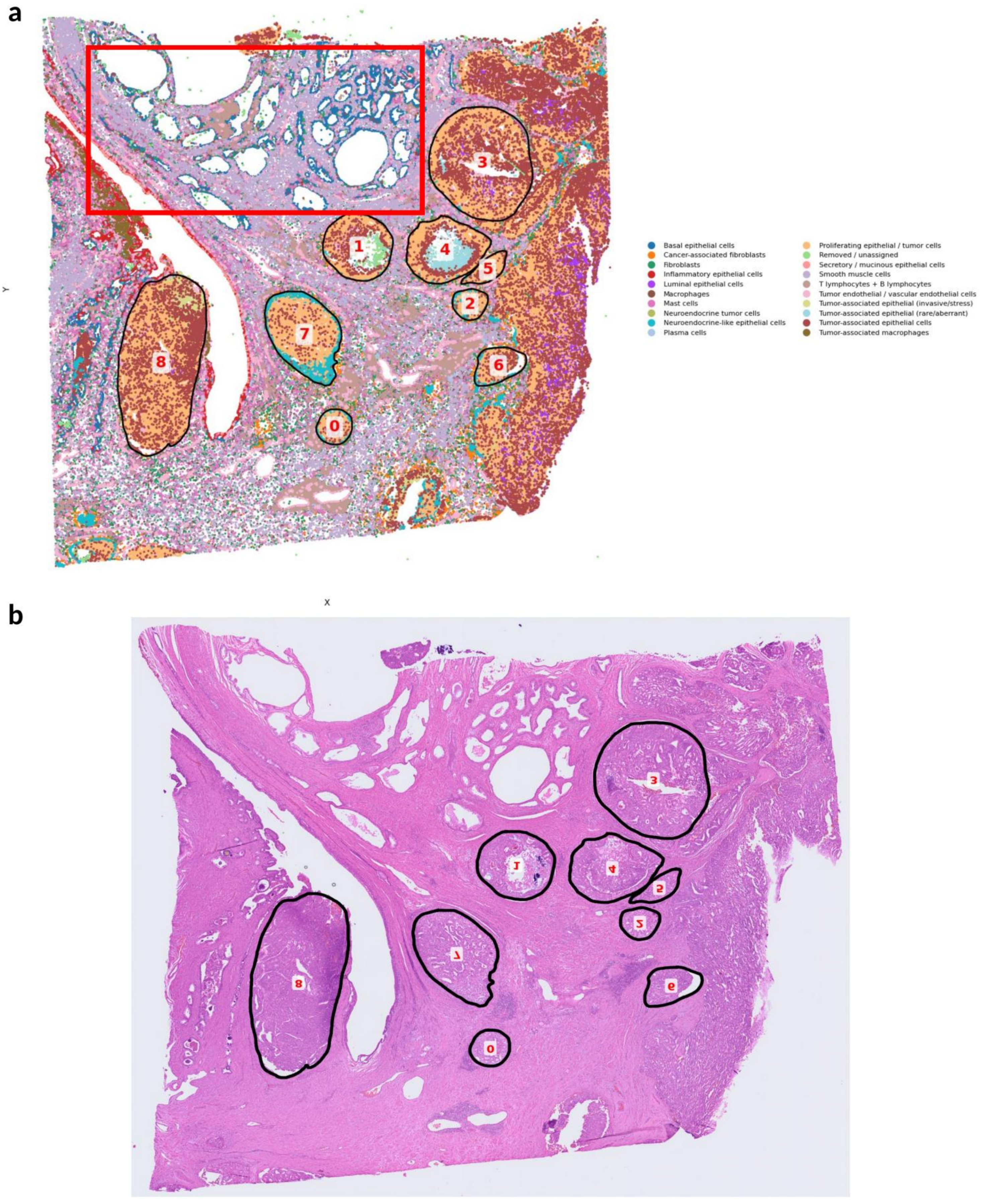
a
b
Y
X
3
1
4
5
2
7
6
8
0
Basal epithelial cells
Cancer-associated fibroblasts
Fibroblasts
Inflammatory epithelial cells
Luminal epithelial cells
Macrophages
Mast cells
Neuroendocrine tumor cells
Neuroendocrine-like epithelial cells
Plasma cells
Proliferating epithelial / tumor cells
Removed / unassigned
Secretory / mucinous epithelial cells
Smooth muscle cells
T lymphocytes + B lymphocytes
Tumor endothelial / vascular endothelial cells
Tumor-associated epithelial (invasive/stress)
Tumor-associated epithelial (rare/aberrant)
Tumor-associated epithelial cells
Tumor-associated macrophages

**Supplementary Figure 4. Spatial annotation of GG4 tumor glands and adjacent benign regions in prostate tissue.**

(a) Whole-section spatial cell-type map showing annotated gland regions used for high-grade prostate cancer analysis. Numbered circular annotations indicate individual GG4 tumor glands, while the red boxed region highlights adjacent benign gland regions used for comparison. Cell-type annotations are shown across the tissue section, including epithelial, stromal, immune, endothelial, and tumor-associated cell populations. (b) Corresponding H&E image of the same tissue section with annotated GG4 tumor glands outlined and numbered. Together, these panels show the spatial context and histologic regions selected for downstream gland-level morphology, radial organization, and inside-versus-outside boundary analyses.

**a**

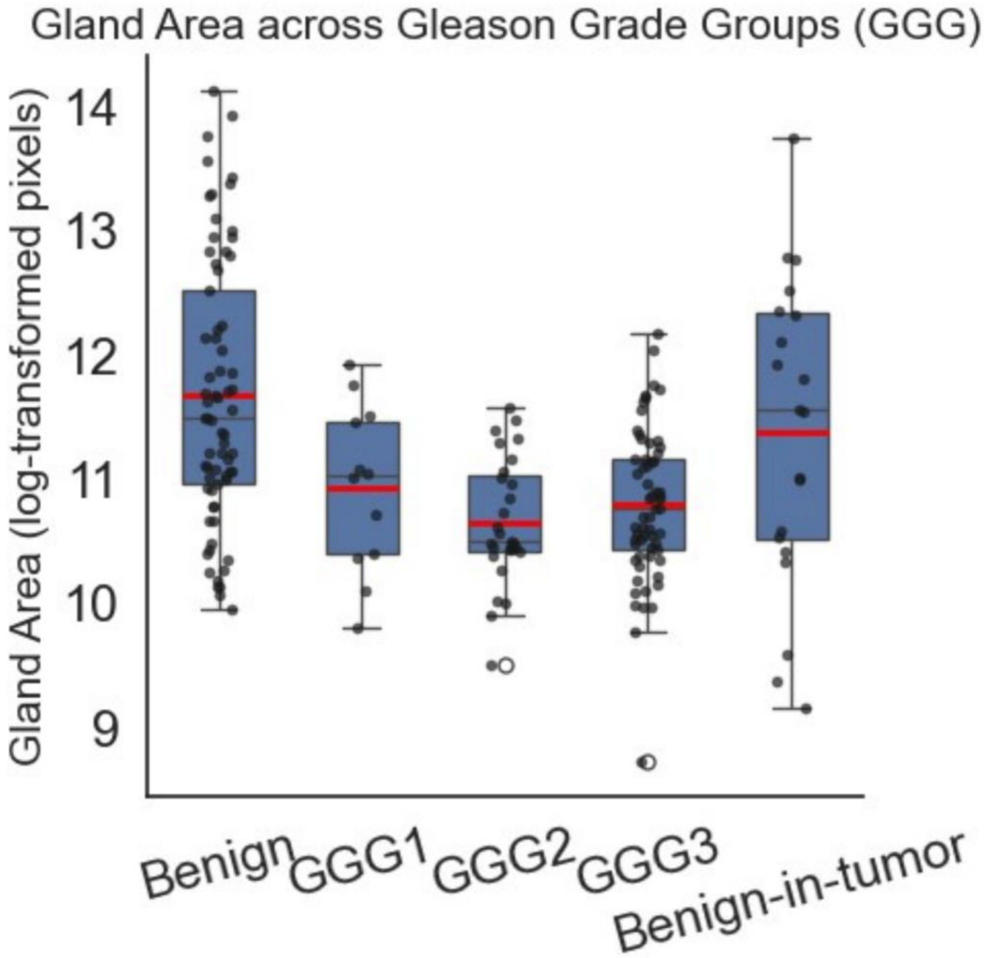

Gland Area across Gleason Grade Groups (GGG)
Gland Area (log-transformed pixels)
14
13
12
11
10
9
Benign
GGG1
GGG2
GGG3
Benign-in-tumor


**b**

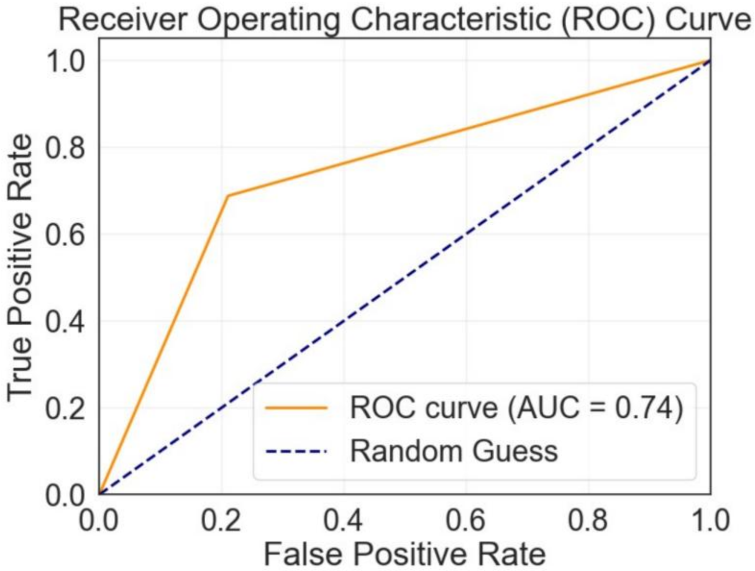

Receiver Operating Characteristic (ROC) Curve
True Positive Rate
False Positive Rate
0.0
0.2
0.4
0.6
0.8
1.0
ROC curve (AUC = 0.74)
Random Guess


**c**

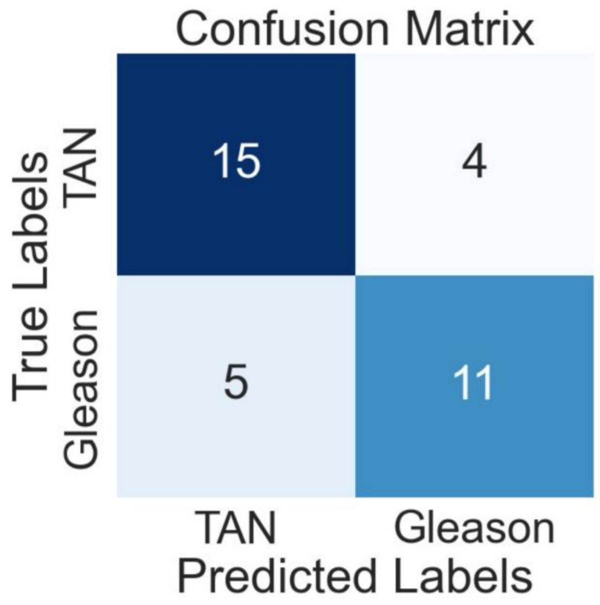

Confusion Matrix
True Labels
TAN
Gleason
15
4
5
11
TAN
Gleason
Predicted Labels

**Supplementary Figure 5. Gland area varies across prostate gland states and morphology-based classification partially separates TAN and Gleason glands.**

(a) Boxplot showing log-transformed gland area across benign, GG1, GG2, GG3, and benign-in-tumor gland categories. Individual points represent glands, boxes indicate the interquartile range, and red horizontal lines indicate group means. (b) Receiver operating characteristic curve for classification of tumor-adjacent normal (TAN) versus Gleason glands using gland-level features. (c) Corresponding confusion matrix showing classification performance. The classifier achieved an AUC of 0.74, with 15 TAN glands and 11 Gleason glands correctly classified, and 4 TAN glands and 5 Gleason glands misclassified. Together, these analyses show that gland-level morphology provides partial separation of TAN and tumor-associated gland states.

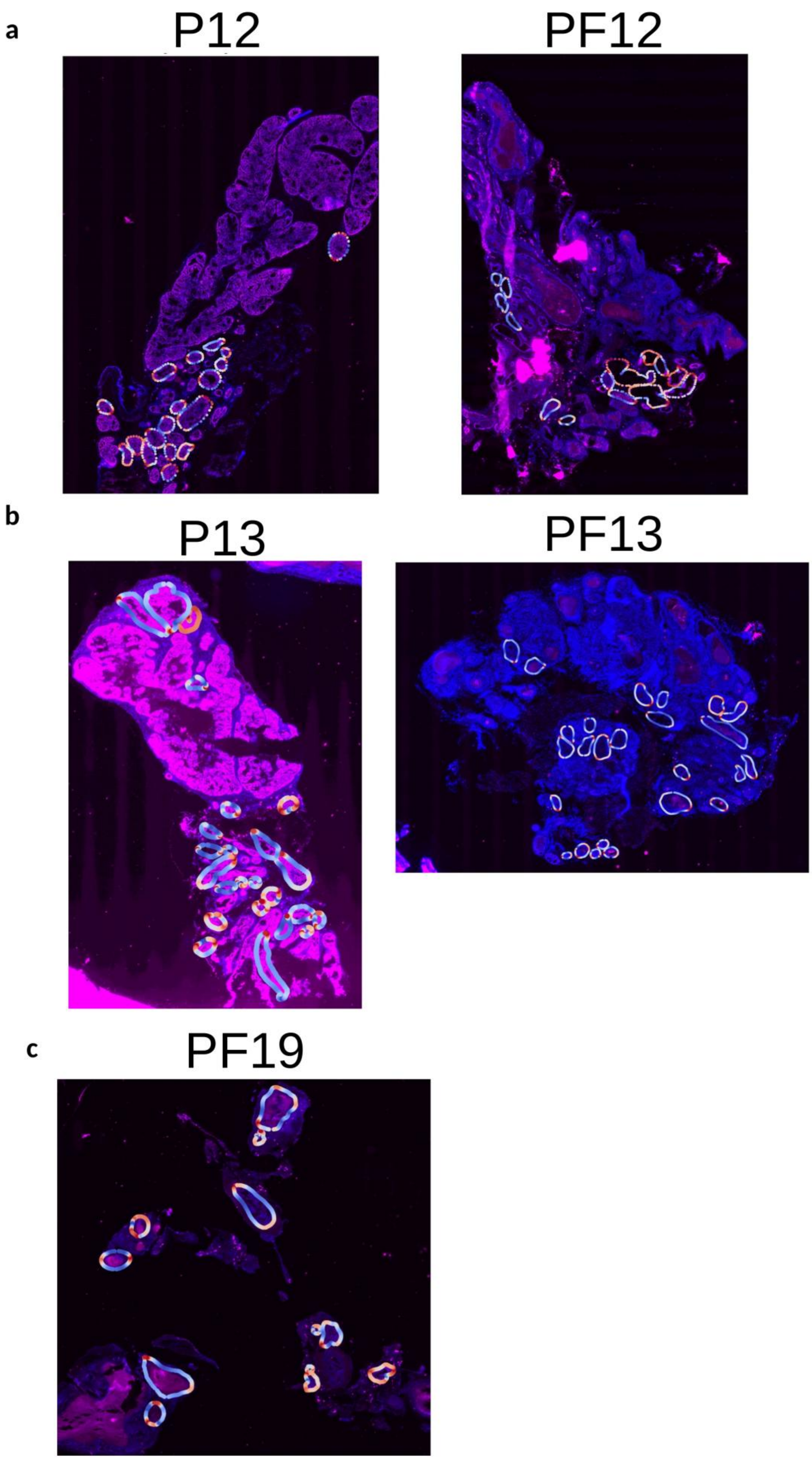
a
P12
PF12
b
P13
PF13
c
PF19

**Supplementary Figure 6. FOXA1 expression and gland morphology across control and FOXA1-knockout prostate tissues.**

Representative fluorescent images showing FOXA1 (magenta) and DAPI (blue) staining in control (P) and Foxa1-knockout (PF) mouse prostate tissues at weeks 12 (a), 13 (b), and 19 (c). The images were overlaid with the segmented glands and curvature computed.

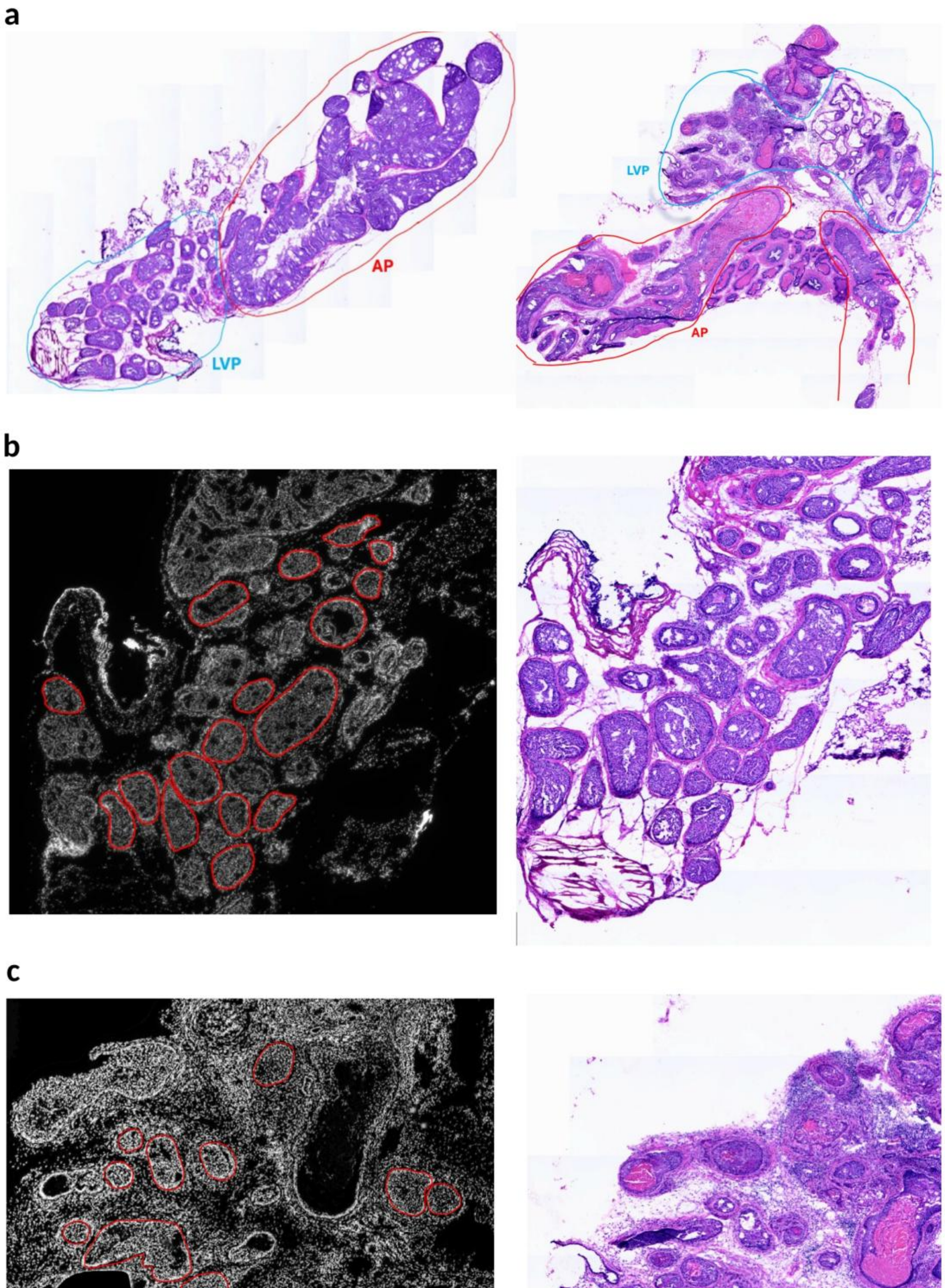
a
AP
LVP
LVP
AP
b
c

**Supplementary Figure 7: Overview of prostate gland segmentation and regional annotation across anterior prostate (AP) and lateral-ventral prostate (LVP) lobes.**

(a)Histological overview of prostate gland sections highlighting the anterior prostate (AP, red) and lateral-ventral prostate (LVP, blue) lobes (P12: left, PF12: right)

(b) Representative fluorescence and corresponding H&E images of the P12 sample showing segmented gland boundaries (red outlines) used for morphological and curvature analyses across different regions of the prostate.

(c) Representative fluorescence and corresponding H&E images of the PF12 sample showing segmented gland boundaries (red outlines)

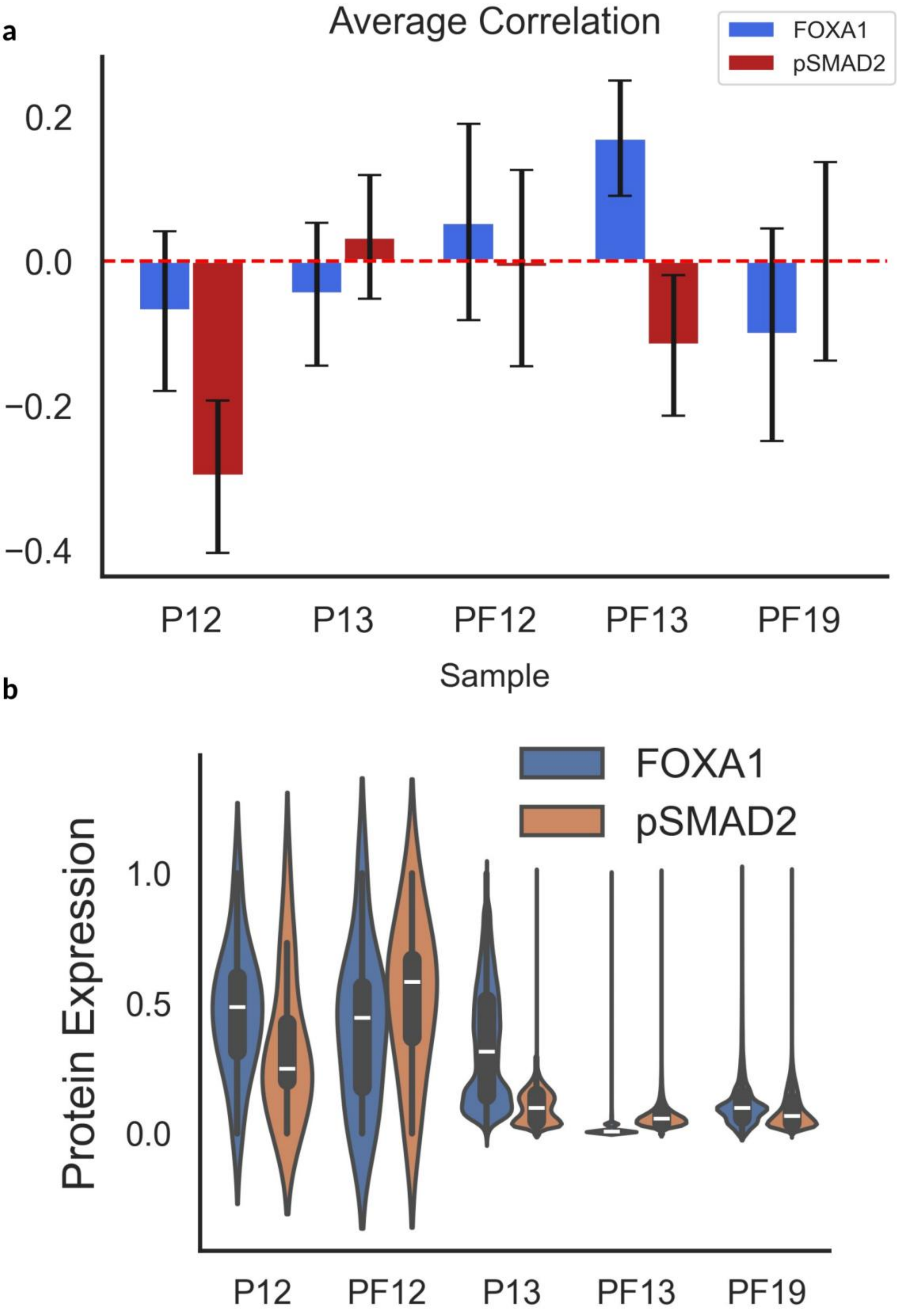
a
Average Correlation
FOXA1
pSMAD2
0.2
0.0
−0.2
−0.4
P12
P13
PF12
PF13
PF19
Sample
b
FOXA1
pSMAD2
Protein Expression
1.0
0.5
0.0
P12
PF12
P13
PF13
PF19

**Supplementary Figure 8. Average correlation between gland curvature and protein expression across samples.**

(a) Bar plot showing the mean curvature correlation values for FOXA1 (blue) and pSMAD2 (red) across control (P) and FOXA1-knockout (PF) mice at weeks 12, 13, and 19. Error bars represent the standard error of the mean across all glands within each sample. Positive correlations indicate higher protein expression in regions of high curvature, whereas negative correlations reflect higher expression in flatter or inward-bending regions.
(b) Violin plots showing the distribution of normalized protein expression levels for FOXA1 (blue) and pSMAD2 (orange) across control (P) and FOXA1-knockout (PF) mice at weeks 12, 13, and 19. FOXA1 expression is markedly reduced in all PF samples, confirming effective knockout, while pSMAD2 expression remains detectable across both cohorts. Interestingly, pSMAD2 expression appears slightly reduced in PF samples compared to their corresponding P samples.

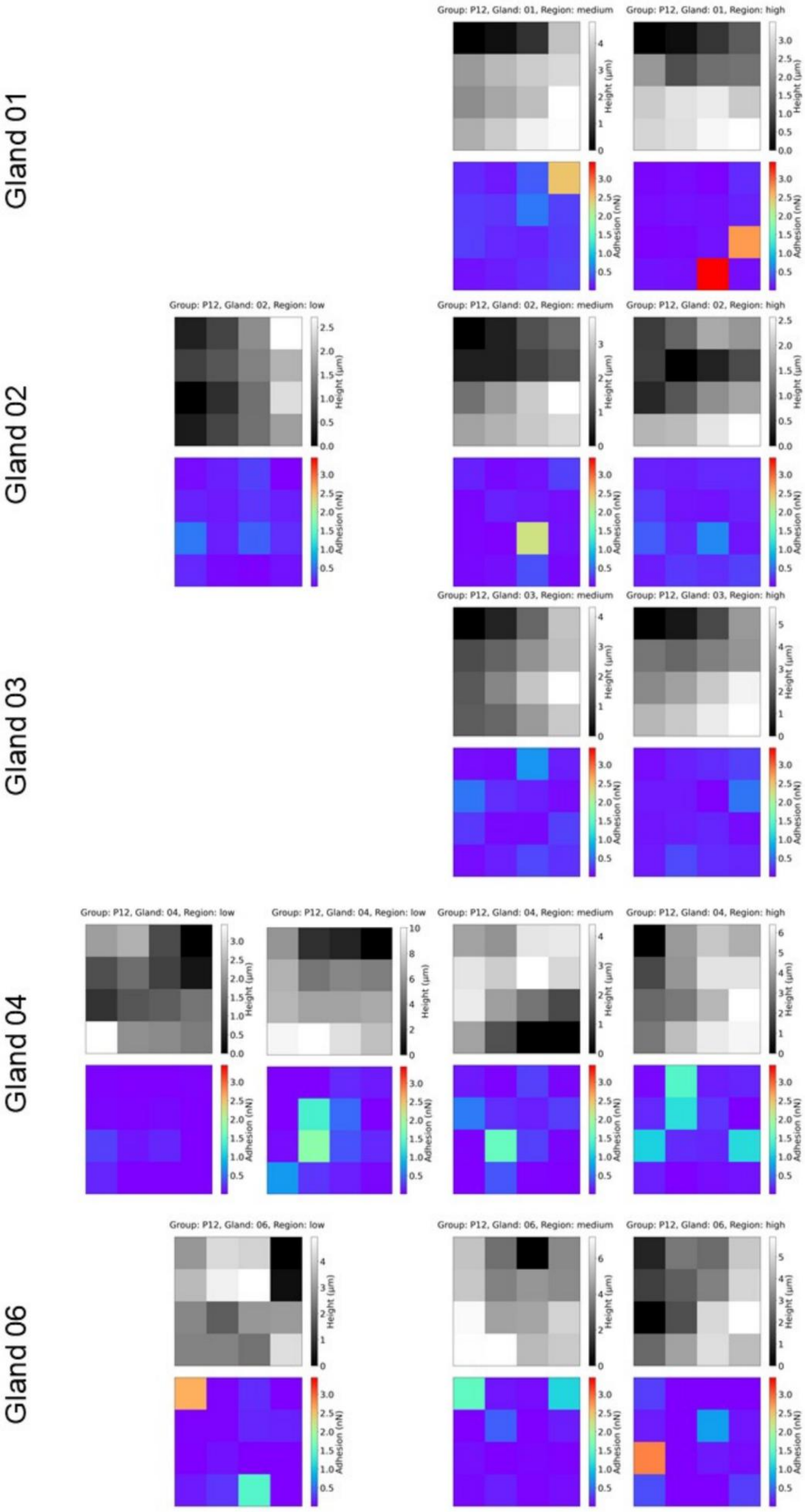

Gland 01
Group: P12, Gland: 01, Region: medium
Group: P12, Gland: 01, Region: high
Gland 02
Group: P12, Gland: 02, Region: low
Group: P12, Gland: 02, Region: medium
Group: P12, Gland: 02, Region: high
Gland 03
Group: P12, Gland: 03, Region: medium
Group: P12, Gland: 03, Region: high
Gland 04
Group: P12, Gland: 04, Region: low
Group: P12, Gland: 04, Region: low
Group: P12, Gland: 04, Region: medium
Group: P12, Gland: 04, Region: high
Gland 06
Group: P12, Gland: 06, Region: low
Group: P12, Gland: 06, Region: medium
Group: P12, Gland: 06, Region: high
Height (μm)
Adhesion (nN)

**Supplementary Figure 9. AFM adhesion and height maps across curvature regions in P12 fresh tissue.**

Representative adhesion (blue–red) and height (grayscale) AFM heatmaps from multiple glands of the P12 sample. Each gland (01–06) includes measurements from regions of low, medium, and high curvature. The adhesion maps reveal localized variability in tip–sample interaction strength, while the height maps illustrate the topographical differences across gland borders.

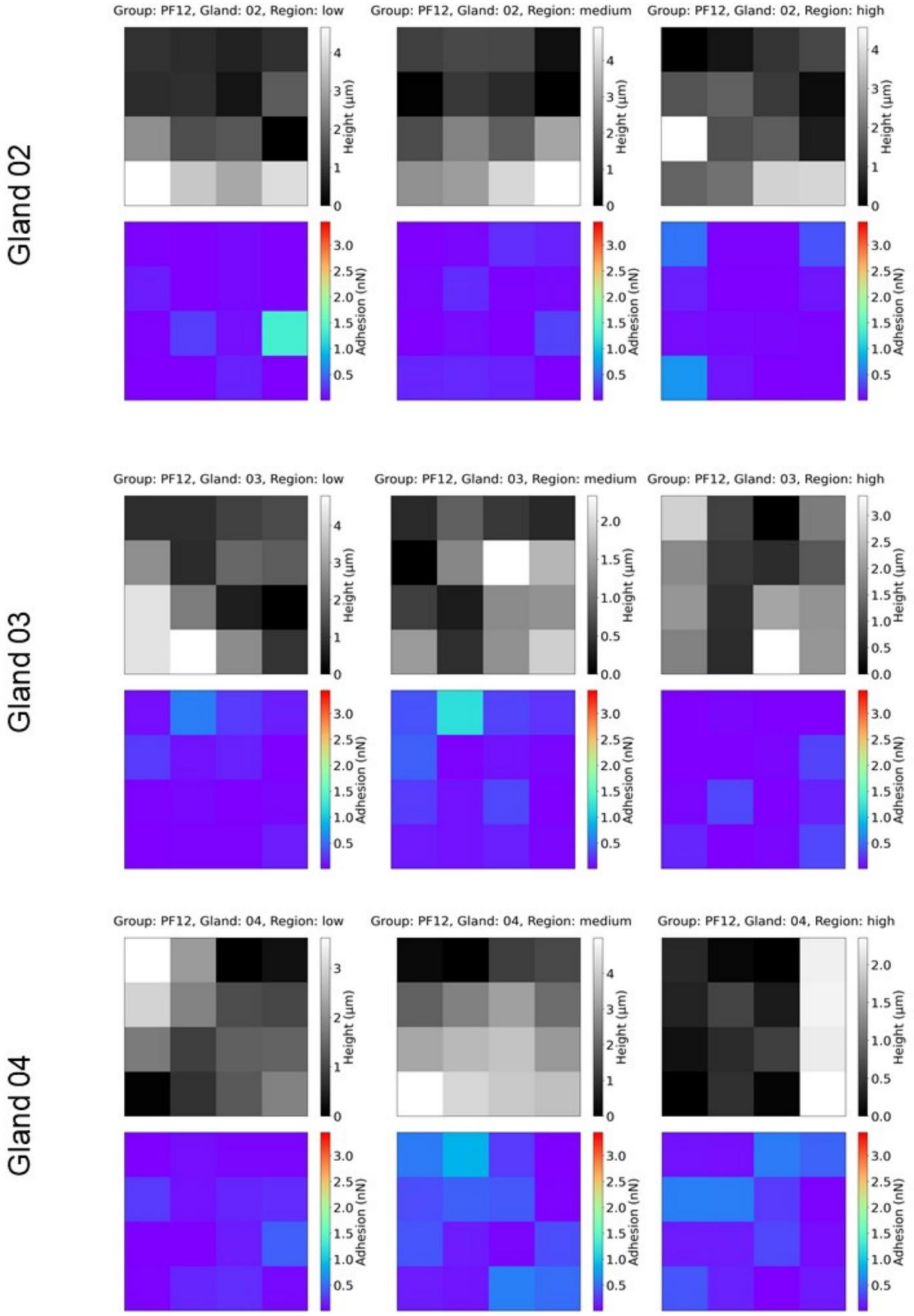

Gland 02
Group: PF12, Gland: 02, Region: low
Group: PF12, Gland: 02, Region: medium
Group: PF12, Gland: 02, Region: high
Gland 03
Group: PF12, Gland: 03, Region: low
Group: PF12, Gland: 03, Region: medium
Group: PF12, Gland: 03, Region: high
Gland 04
Group: PF12, Gland: 04, Region: low
Group: PF12, Gland: 04, Region: medium
Group: PF12, Gland: 04, Region: high
Height (µm)
Adhesion (nN)

**Supplementary Figure 10. AFM adhesion and height maps across curvature regions in PF12 fresh tissue.**

Representative AFM-derived adhesion (blue–red) and height (grayscale) heatmaps for multiple glands (02–04) from the PF12 sample. Each gland includes measurements from low, medium, and high curvature regions. Compared to P12, PF12 tissues exhibit greater variability in adhesion across curvature regions, indicating altered
surface mechanics in the FOXA1-knockout sample.

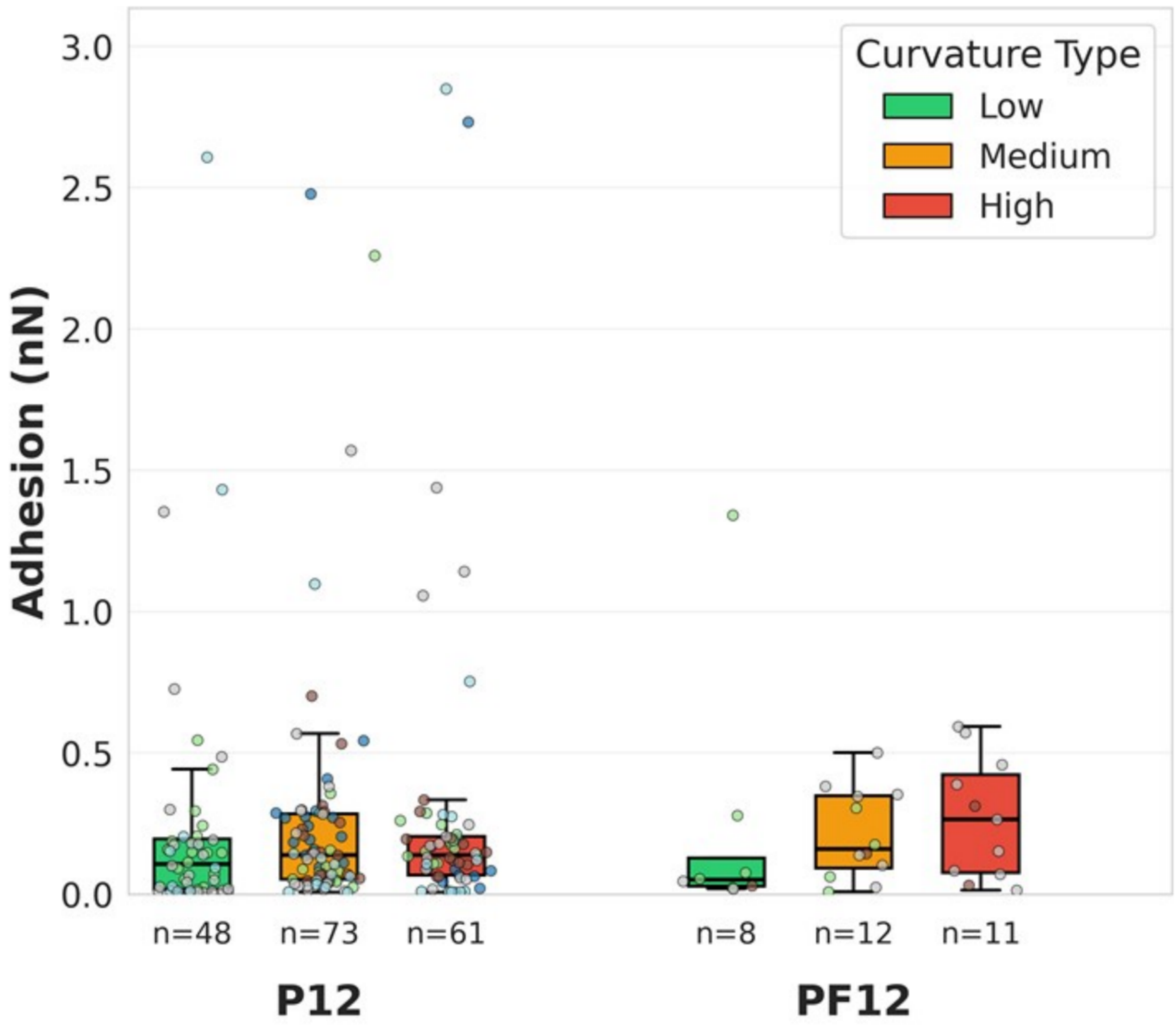

Curvature Type
Low
Medium
High
Adhesion (nN)
3.0
2.5
2.0
1.5
1.0
0.5
0.0
n=48
n=73
n=61
n=8
n=12
n=11
P12
PF12

**Supplementary Figure 11. Quantification of AFM adhesion across curvature regions in fresh tissues.**

Boxplots showing adhesion forces (in nN) measured in P12 and PF12 fresh tissue samples across regions of low (green), medium (orange), and high (red) curvature. P12 tissues exhibit relatively uniform adhesion across curvature types, whereas PF12 tissues display higher variability, particularly in high-curvature regions, indicating altered surface mechanics following FOXA1 knockout.

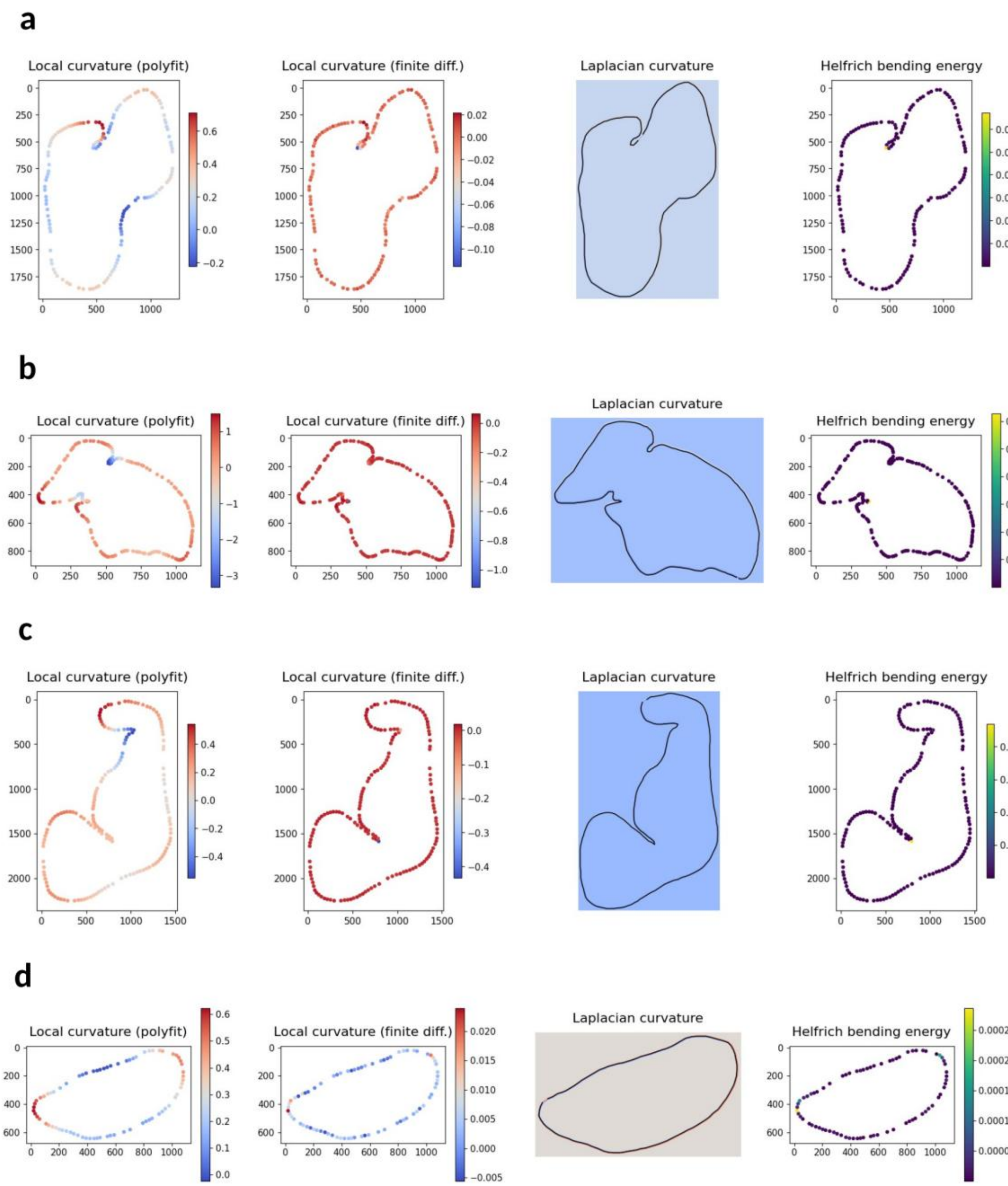
a
Local curvature (polyfit)
Local curvature (finite diff.)
Laplacian curvature
Helfrich bending energy
b
Local curvature (polyfit)
Local curvature (finite diff.)
Laplacian curvature
Helfrich bending energy
c
Local curvature (polyfit)
Local curvature (finite diff.)
Laplacian curvature
Helfrich bending energy
d
Local curvature (polyfit)
Local curvature (finite diff.)
Laplacian curvature
Helfrich bending energy

**Supplementary Figure 12. Comparison of curvature estimation methods.**

Examples of prostate gland contours analyzed using different curvature computation approaches, including polynomial fitting, finite differences, Laplacian curvature, and Helfrich bending energy.

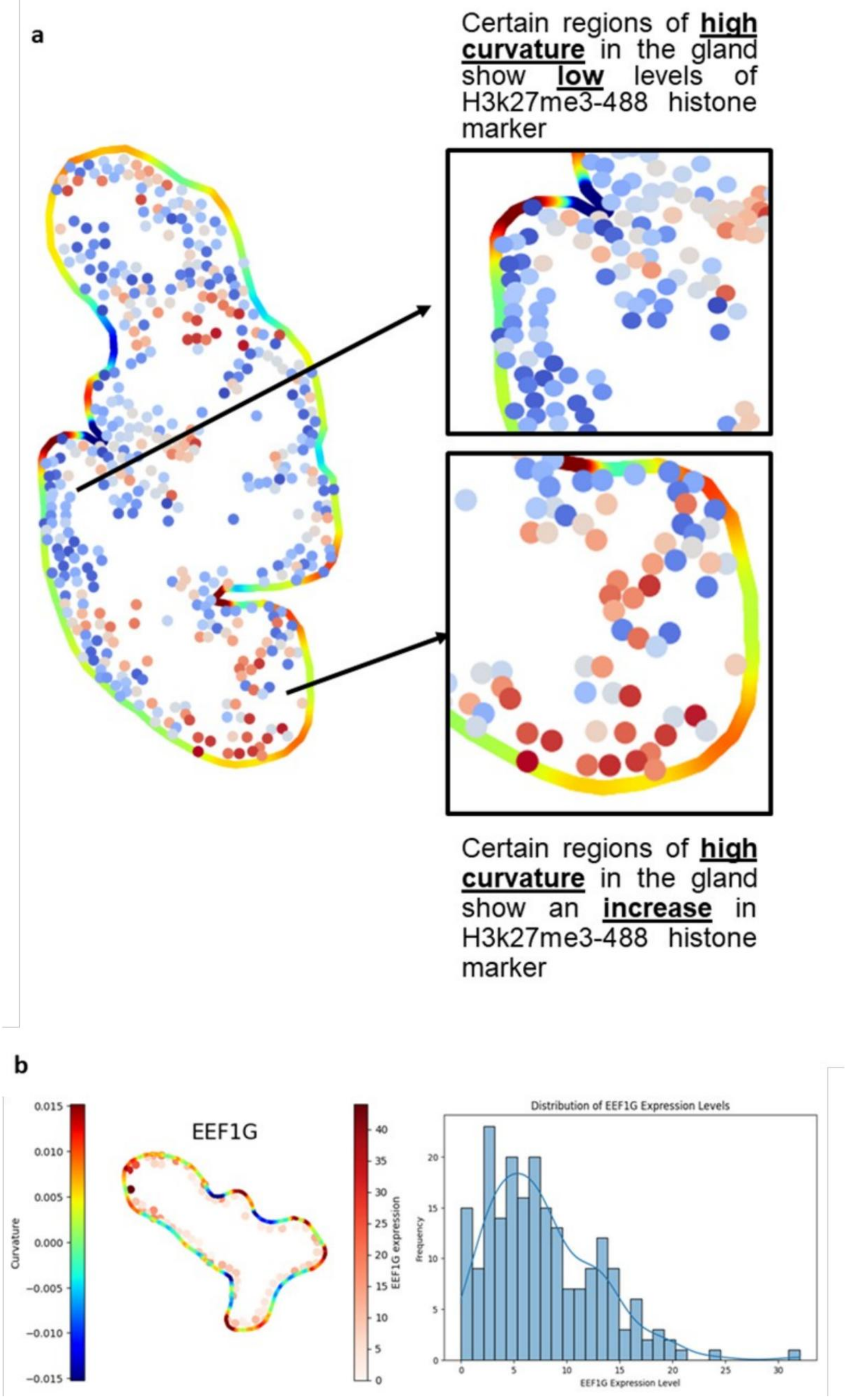
a
Certain regions of high curvature in the gland show low levels of H3k27me3-488 histone marker
Certain regions of high curvature in the gland show an increase in H3k27me3-488 histone marker
b
EEF1G
Curvature
EEF1G expression
Distribution of EEF1G Expression Levels
Frequency
EEF1G Expression Level

**Supplementary Figure 13. Regional curvature-dependent variation in histone marker and gene expression.**

(a) High-curvature regions within a representative gland exhibit differential levels of the histone modification marker H3K27me3. In some areas, high curvature coincides with reduced H3K27me3 signal (top inset), whereas in other regions, high curvature is associated with increased H3K27me3 signal (bottom inset).

(b) Example gland showing curvature mapping (left) and spatial distribution of EEF1G expression projected onto the gland boundary (middle), with corresponding histogram of EEF1G expression levels across cells (right).

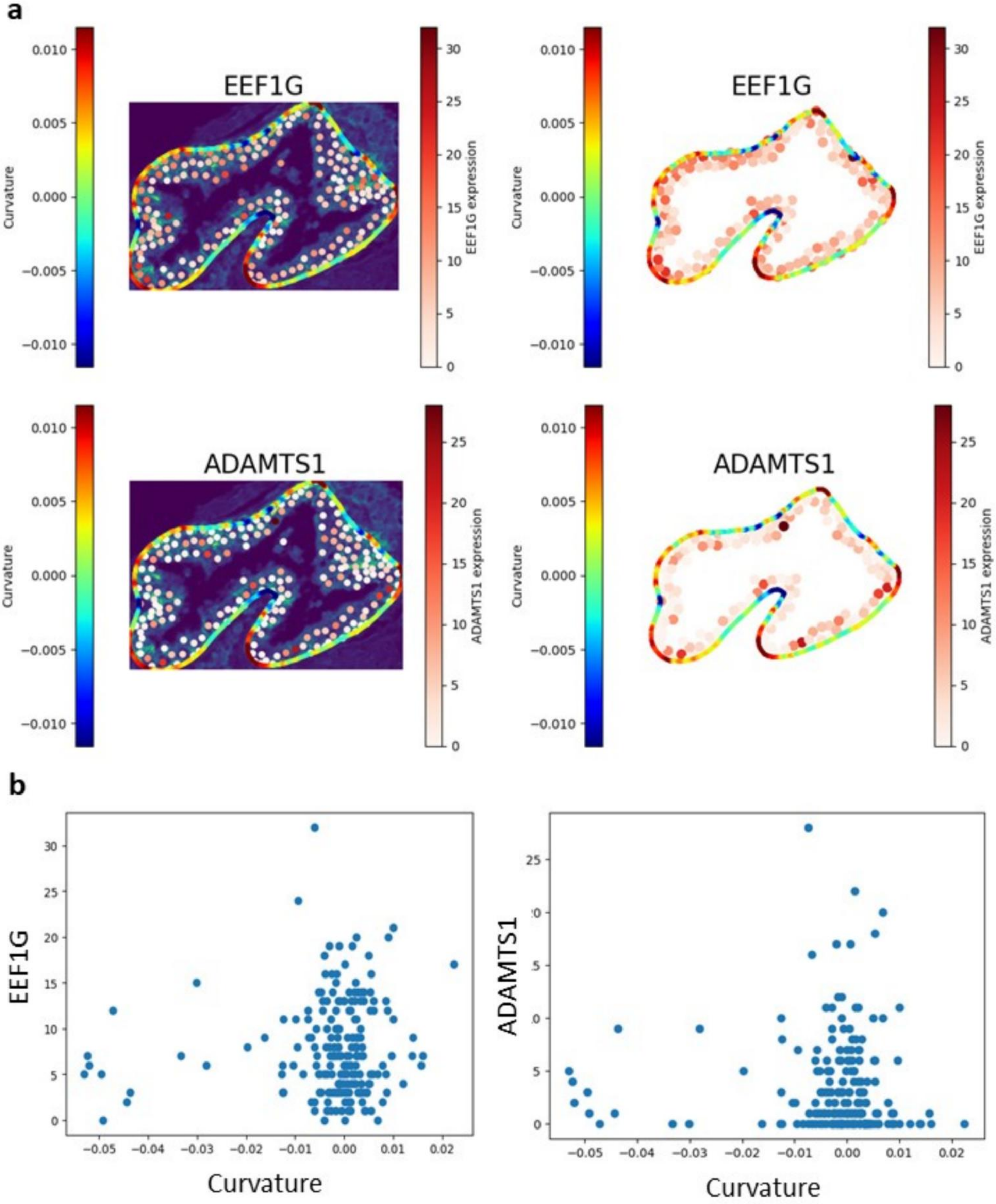

a
EEF1G
Curvature
EEF1G expression
ADAMTS1
ADAMTS1 expression
0.010
0.005
0.000
−0.005
−0.010
30
25
20
15
10
5
0
b
EEF1G
ADAMTS1
Curvature
−0.05
−0.04
−0.03
−0.02
−0.01
0.00
0.01
0.02

**Supplementary Figure 14. Relationship between gland curvature and gene expression at the single-cell level (Gland 1).**

(a) Representative glands with local curvature values mapped along the boundaries (color scale: blue = inward/concave, red = outward/convex). Expression levels of EEF1G (top) and ADAMTS1 (bottom) are projected onto individual cells along the gland contour, shown on background tissue (left) and isolated boundary plots (right).

(b) Scatter plots of curvature values (x-axis) versus gene expression (y-axis) for individual cells within the glands.

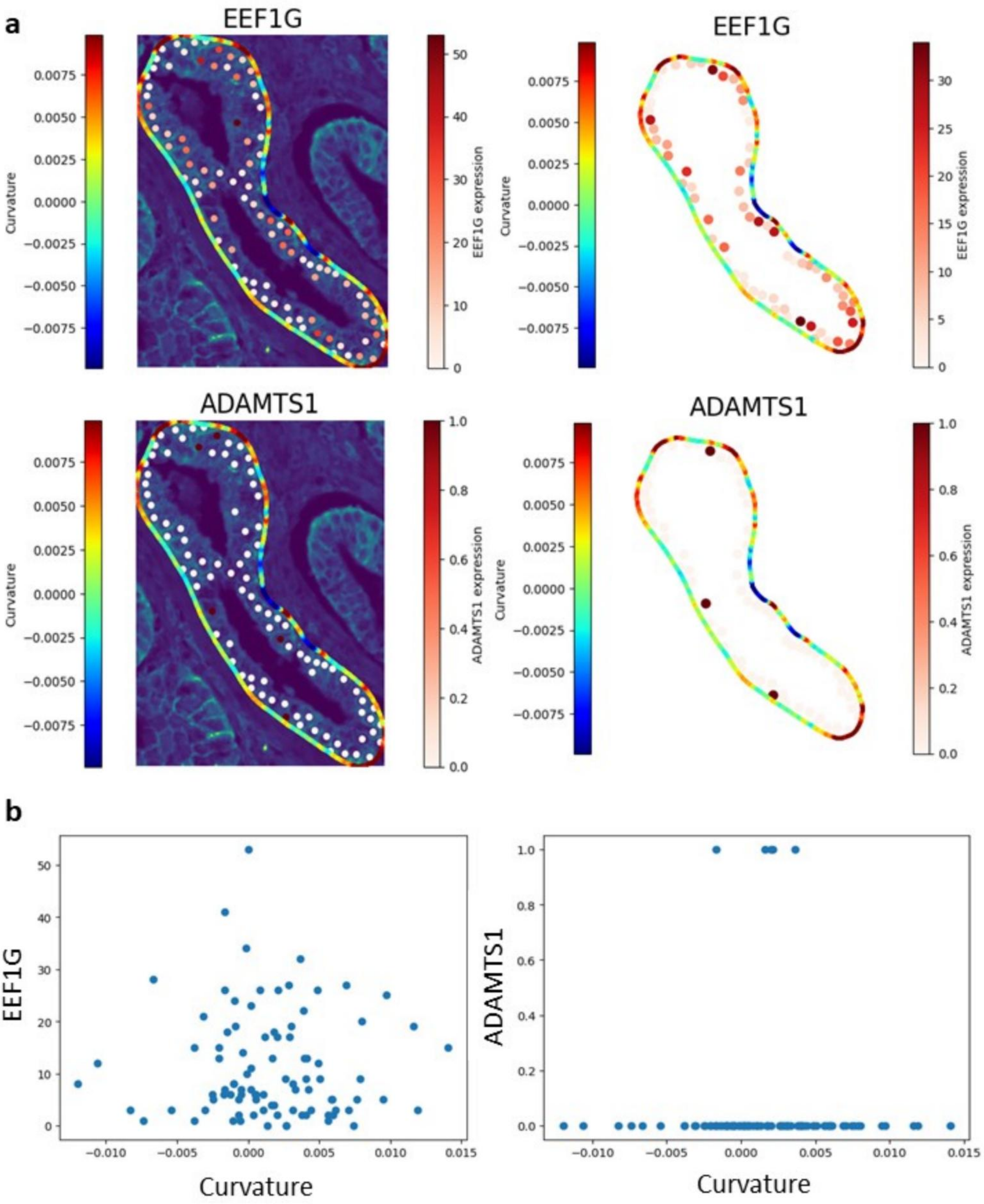

a
EEF1G
Curvature
EEF1G expression
ADAMTS1
ADAMTS1 expression
b
EEF1G
ADAMTS1
Curvature

**Supplementary Figure 15. Relationship between gland curvature and gene expression at the single-cell level (Gland 2).**

(a) Representative glands with curvature values mapped along the boundary (color scale: red = outward/convex, blue = inward/concave). Expression of EEF1G (top) and ADAMTS1 (bottom) is projected onto individual cells, shown both on the tissue background (left) and along the isolated gland contour (right).

(b) Scatter plots showing the relationship between local curvature (x-axis) and gene expression (y-axis) for individual cells. Each point represents a single cell located along the gland boundary. EEF1G exhibits a broad dynamic range of expression across curvature values, whereas ADAMTS1 expression is more restricted, with most cells displaying low expression regardless of curvature.

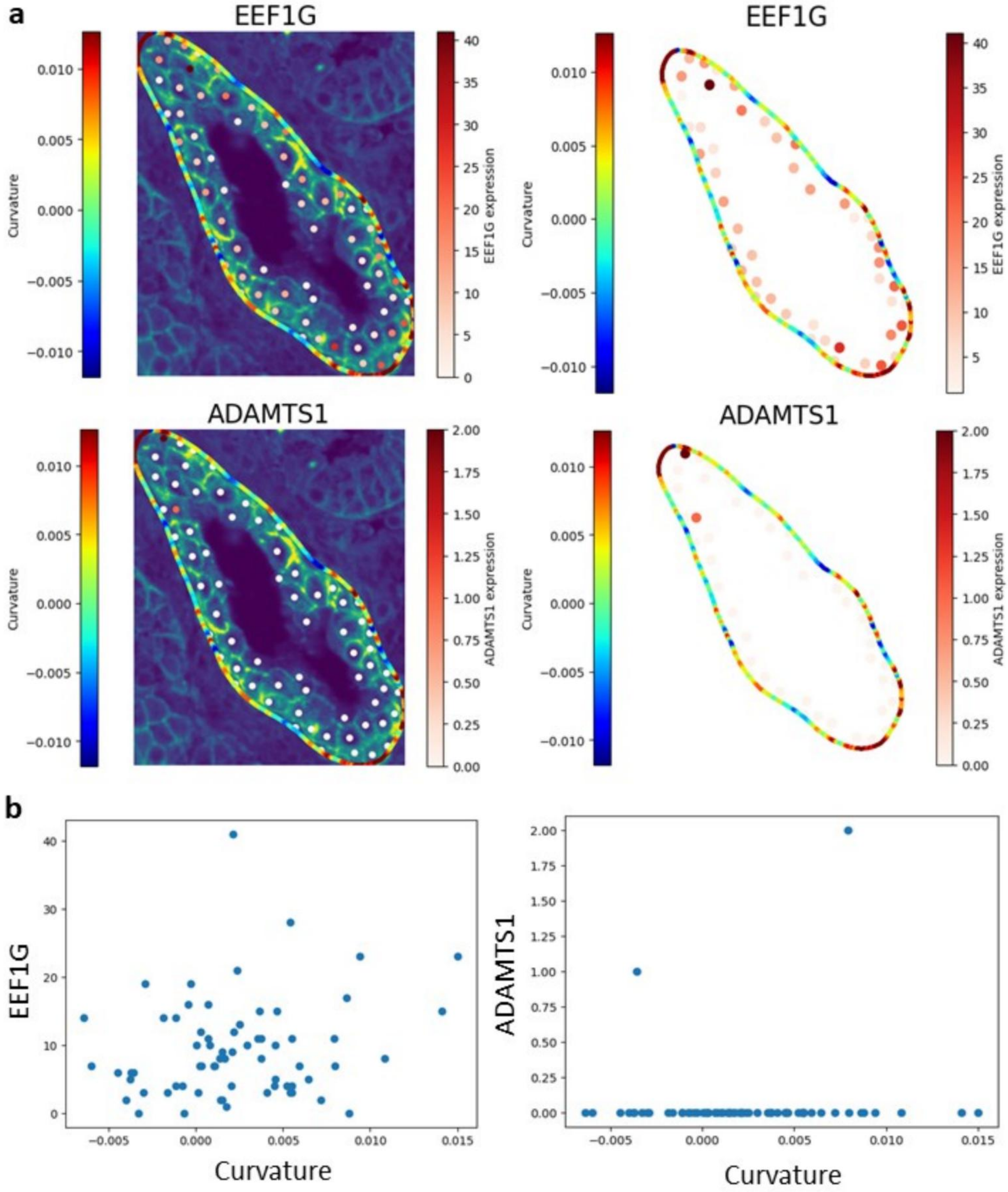
a
EEF1G
Curvature
EEF1G expression
EEF1G
Curvature
EEF1G expression
ADAMTS1
Curvature
ADAMTS1 expression
ADAMTS1
Curvature
ADAMTS1 expression
b
EEF1G
Curvature
ADAMTS1
Curvature

**Supplementary Figure 16. Relationship between gland curvature and gene expression at the single-cell level (Gland 3).**

(a) Representative glands showing curvature mapped along the boundary (left) and expression levels of EEF1G (top) and ADAMTS1 (bottom) projected onto individual cells (color intensity reflects gene expression levels). The right panels display the same cells mapped along the isolated gland contour, highlighting spatial variation in expression with respect to curvature.

(b) Scatter plots showing the association between local curvature values (x-axis) and gene expression (y-axis) for EEF1G (left) and ADAMTS1 (right). EEF1G displays a wide range of expression values distributed across curvature levels, while ADAMTS1 expression remains low in most cells with occasional outliers of higher expression.

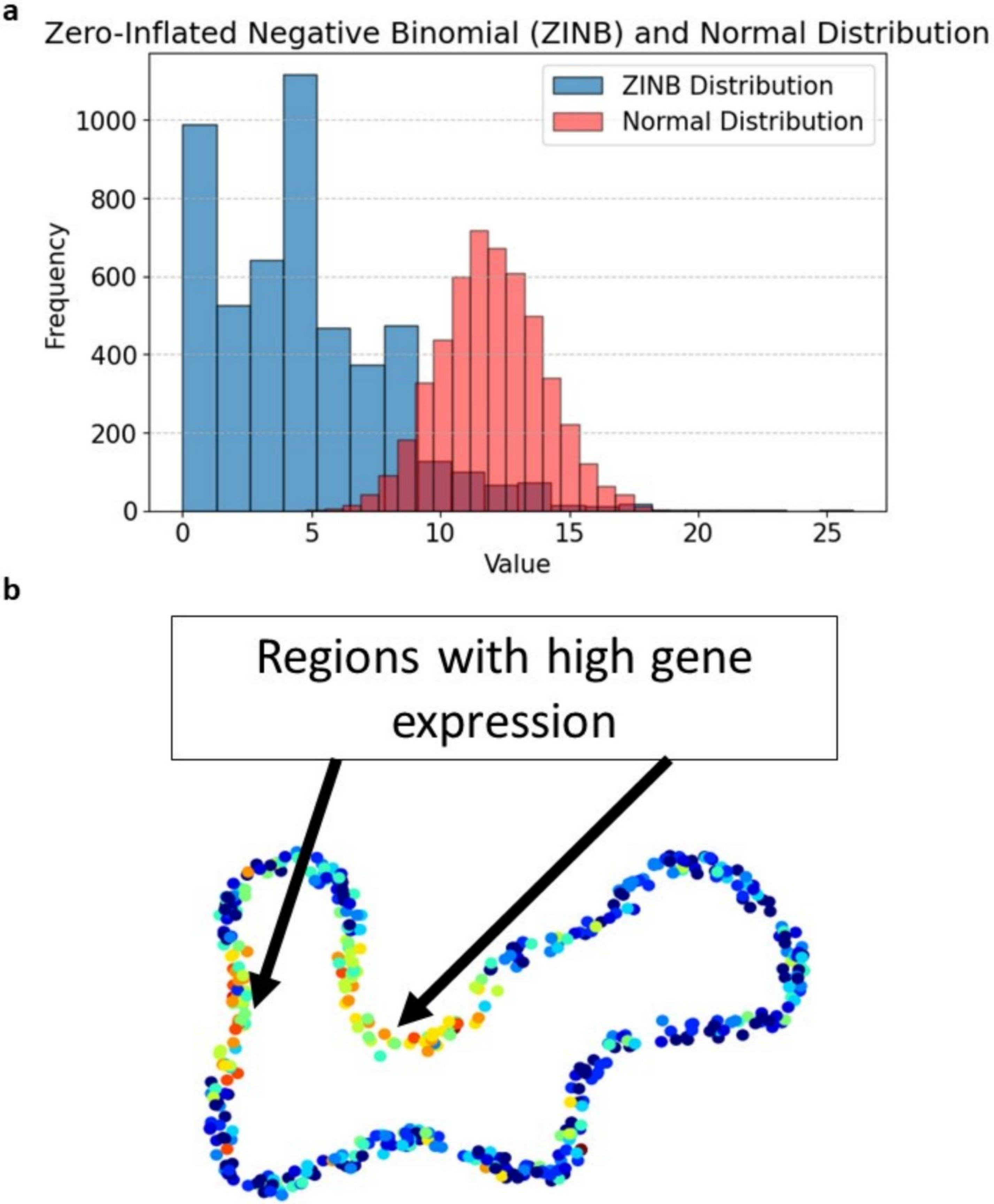
a
Zero-Inflated Negative Binomial (ZINB) and Normal Distribution
ZINB Distribution
Normal Distribution
Frequency
0
200
400
600
800
1000
0
5
10
15
20
25
Value
b
Regions with high gene expression

**Supplementary Figure 17. Characteristics of transcriptomics data distribution and spatial gene expression patterns.**

(a) Comparison of a Zero-Inflated Negative Binomial (ZINB) distribution (blue) with a Normal distribution (red).

(b) Example gland showing spatial gene expression projected onto individual cells. Regions of high expression are highlighted along the gland boundary, illustrating localized variability in expression patterns relative to tissue morphology.

**a**

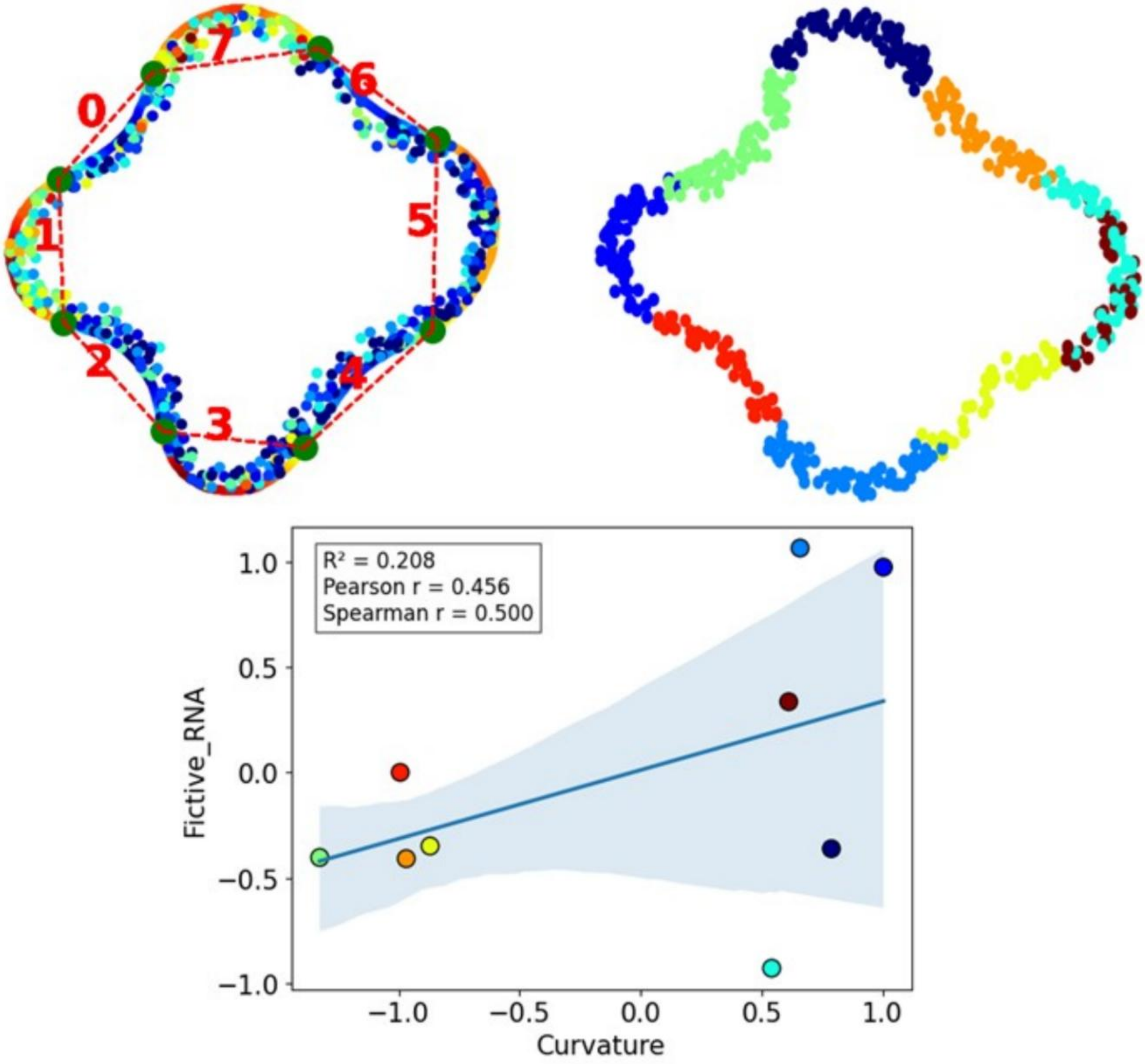

0
1
2
3
4
5
6
7
R² = 0.208
Pearson r = 0.456
Spearman r = 0.500
Fictive_RNA
Curvature


**b**

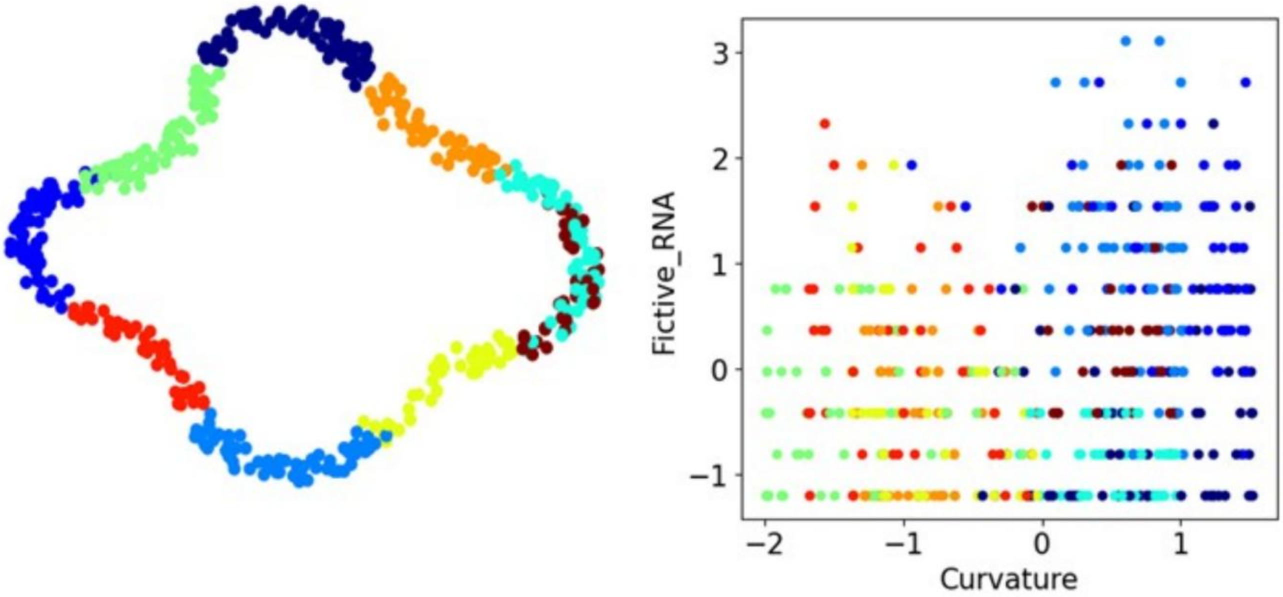

Fictive_RNA
Curvature

**Supplementary Figure 18. Simulation of curvature–RNA relationships in synthetic glands.**
(a) Simulated gland structure with cells arranged along the border. Left: example gland annotated with numbered regions (red) and curvature mapping (color scale). Right: Cells color-coded by clustering from spatial proximity and projected curvature. Bottom: aggregated fictive RNA expression (y-axis) plotted against local curvature (x-axis), showing a moderate positive correlation (Pearson r = 0.456, Spearman r = 0.500).
(b) scatter plot of curvature values versus fictive RNA expression for individual cells, illustrating the heterogeneous distribution of expression relative to curvature. The colors from the scatter plot correspond to the colors of the clusters.

**a**

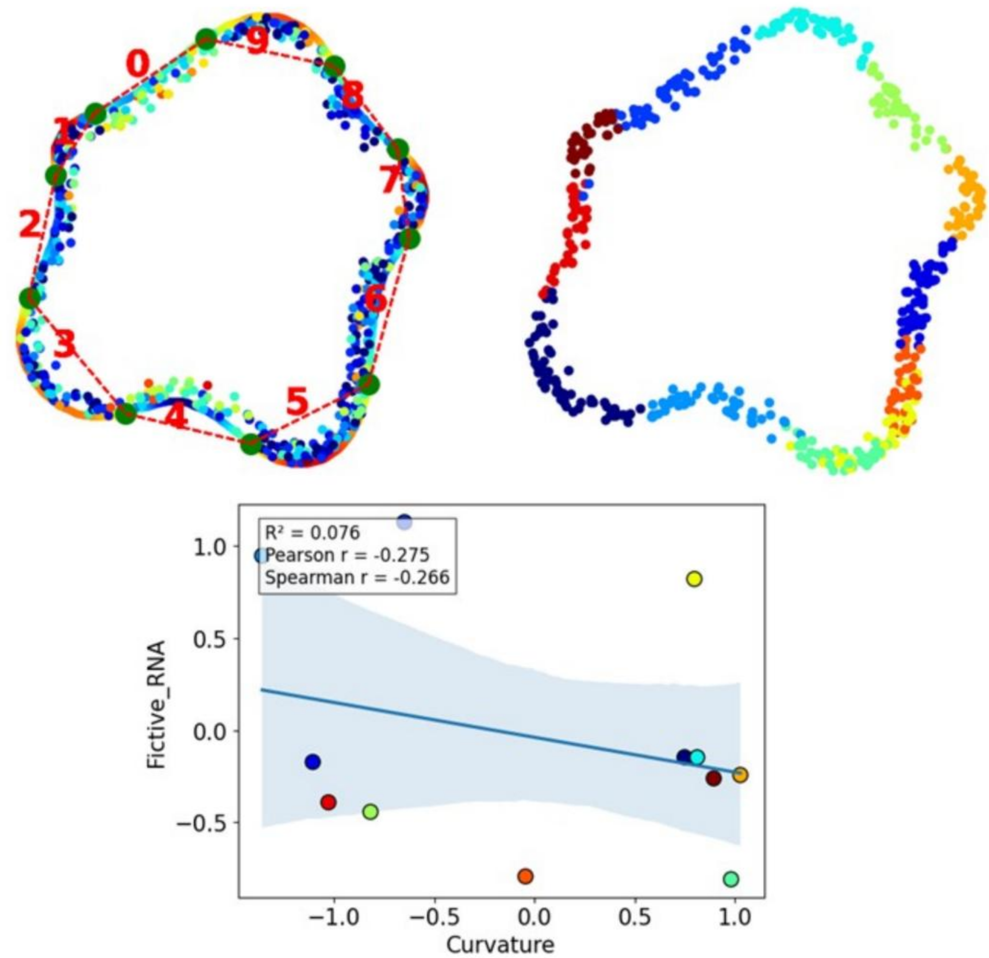
0
1
2
3
4
5
6
7
8
9
R² = 0.076
Pearson r = -0.275
Spearman r = -0.266
Fictive_RNA
Curvature
1.0
0.5
0.0
−0.5
−1.0
−0.5
0.0
0.5
1.0

**b**

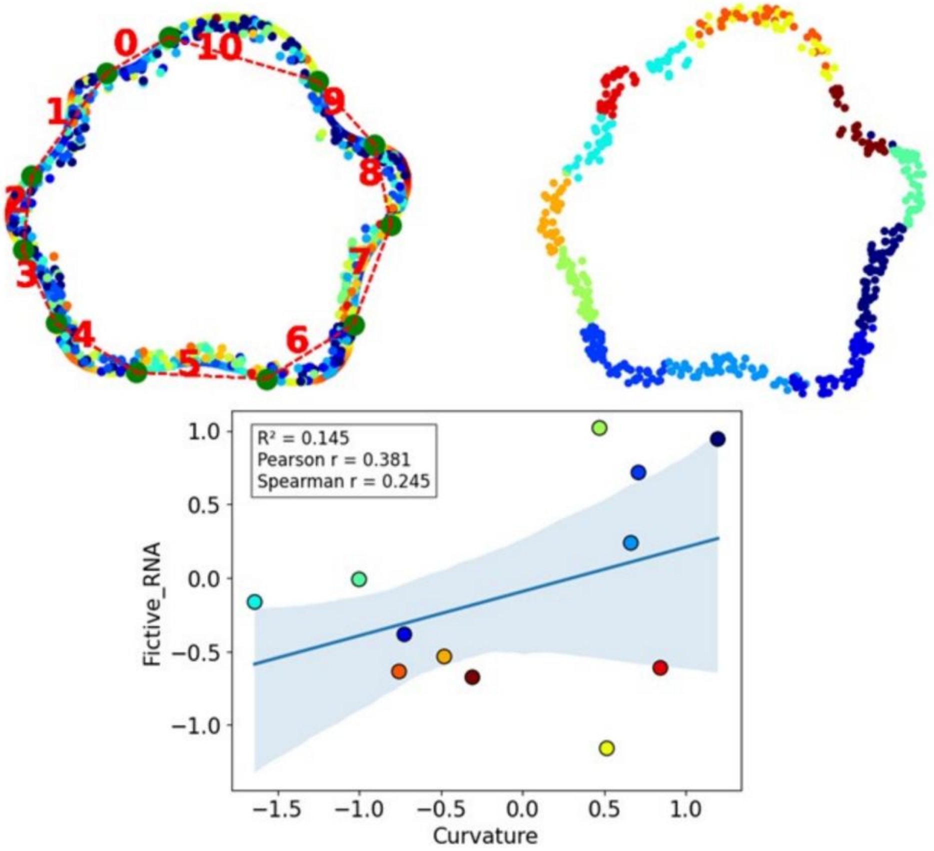
0
1
2
3
4
5
6
7
8
9
10
R² = 0.145
Pearson r = 0.381
Spearman r = 0.245
Fictive_RNA
Curvature
1.0
0.5
0.0
−0.5
−1.0
−1.5
−1.0
−0.5
0.0
0.5
1.0

**Supplementary Figure 19. Simulated examples of curvature–RNA relationships showing positive and negative trends.**

(a) Example synthetic gland where fictive RNA expression was randomly assigned to specific regions. Left: annotated gland contour with numbered segments. Right: Cells color-coded by clustering from spatial proximity and projected curvature. Bottom: aggregated fictive RNA expression plotted against curvature, showing a weak negative correlation (Pearson r = –0.275, Spearman r = –0.266).

(b) Another synthetic gland simulation with localized fictive RNA expression. Left: annotated contour and segmentation. Right: Cells color-coded by clustering from spatial proximity and projected curvature. Bottom: scatter plot of curvature values versus fictive RNA expression, showing a moderate positive correlation (Pearson r= 0.381, Spearman r= 0.245).

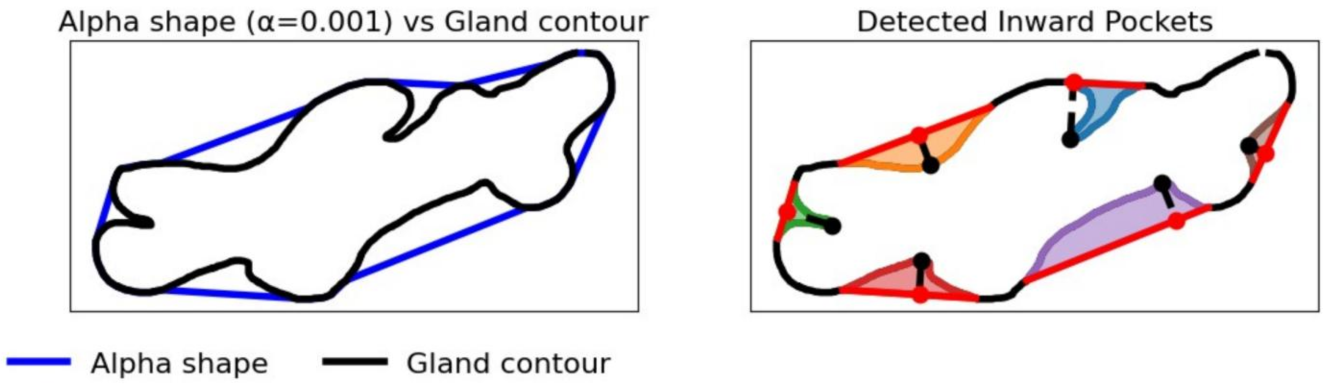

Alpha shape (α=0.001) vs Gland contour
Detected Inward Pockets
Alpha shape
Gland contour

**Supplementary Figure 20. Alpha-shape-based detection of inward pocket regions from gland contours.**

Representative example illustrating the computational approach used to identify inward pocket regions along gland boundaries. The left panel shows the original gland contour in black and the corresponding alpha shape generated using $\alpha = 0.001$ in blue. The right panel shows detected inward pockets defined from deviations between the gland contour and the alpha-shape boundary. Colored pocket regions indicate candidate inward invaginations, while connecting lines mark the corresponding pocket openings used to define pocket geometry.

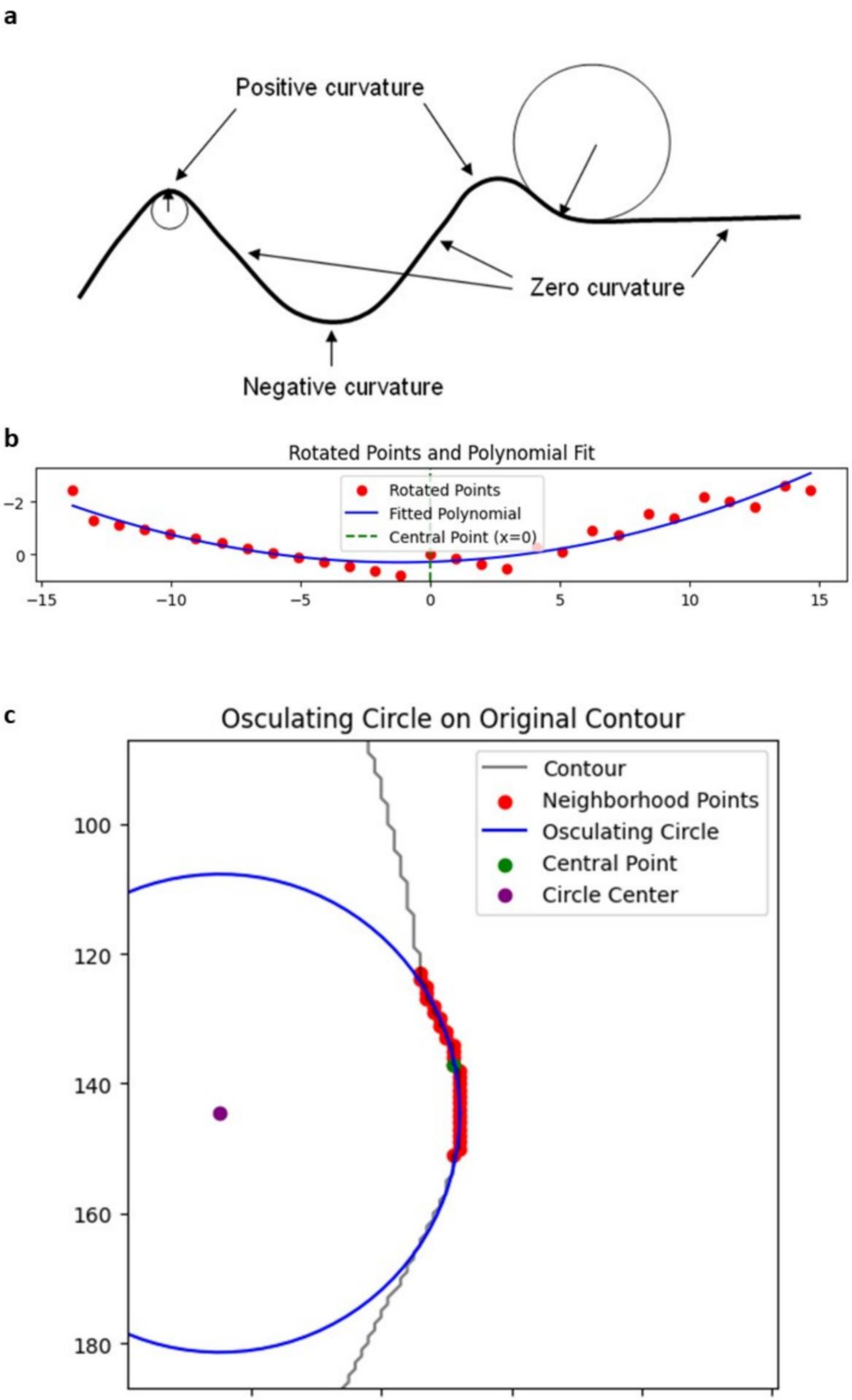

a
Positive curvature
Zero curvature
Negative curvature
b
Rotated Points and Polynomial Fit
Rotated Points
Fitted Polynomial
Central Point (x=0)
−2
0
−15
−10
−5
0
5
10
15
c
Osculating Circle on Original Contour
Contour
Neighborhood Points
Osculating Circle
Central Point
Circle Center
100
120
140
160
180
620
640
660
680
700

**Supplementary Figure 21. Illustration of curvature computation using polynomial fitting and osculating circles.**
(a) Conceptual diagram showing regions of positive, negative, and zero curvature along a curve. Positive curvature corresponds to outward bending, negative curvature to inward bending, and zero curvature to flat regions.

https://csegrecorder.com/articles/view/seismic-curvature-attributes-for-mapping-faults-fractures-and-other

(b) Example of curvature estimation: local contour points (red) are rotated and fit with a second-order polynomial (blue line). The curvature is computed at the central point (green).
(c) Geometric interpretation of curvature via the osculating circle. Neighborhood points along the gland contour (red) define the local shape, and the osculating circle (blue) approximates curvature at the central point (green) with the circle center shown in purple.

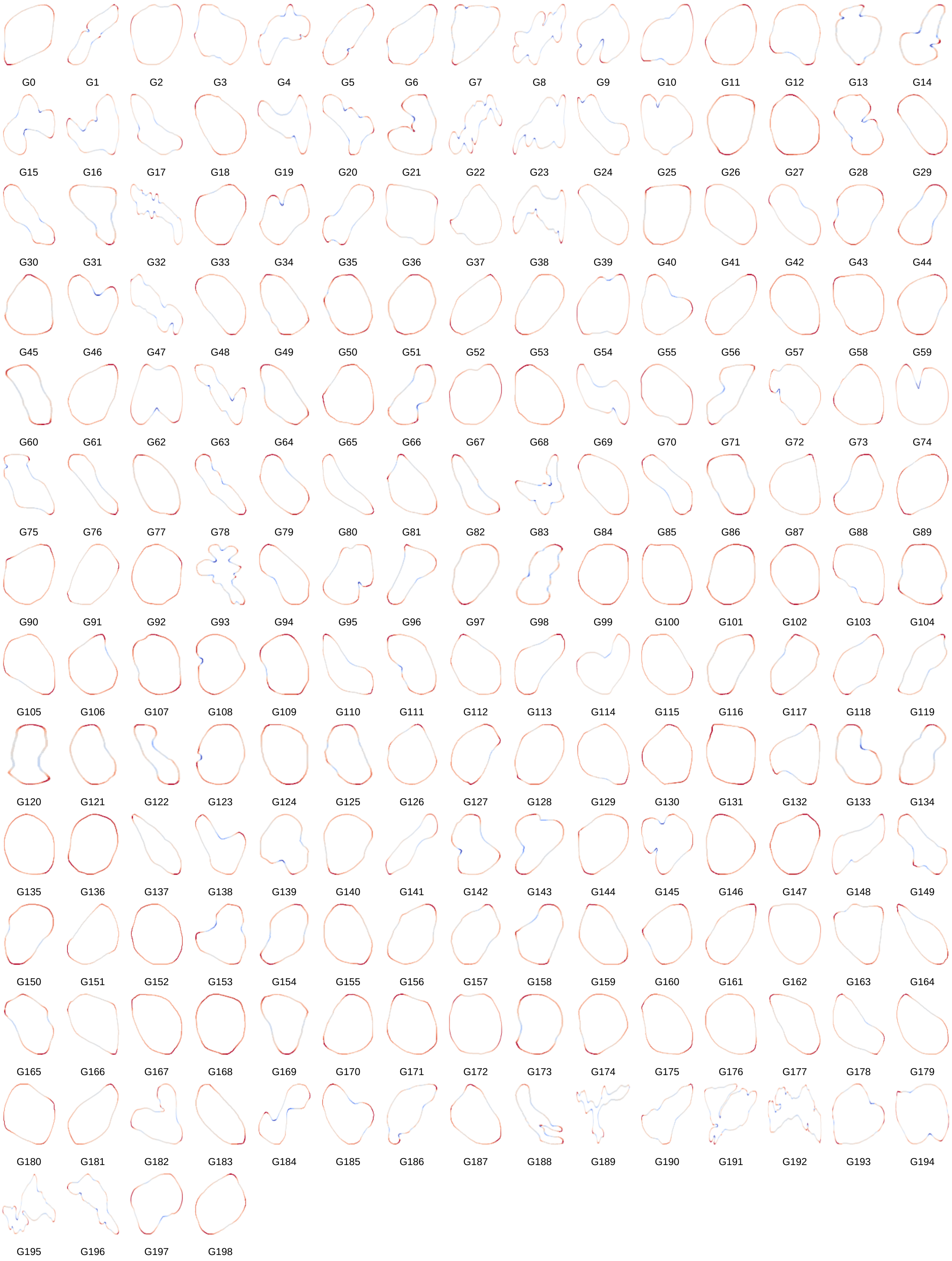
G0 G1 G2 G3 G4 G5 G6 G7 G8 G9 G10 G11 G12 G13 G14
G15 G16 G17 G18 G19 G20 G21 G22 G23 G24 G25 G26 G27 G28 G29
G30 G31 G32 G33 G34 G35 G36 G37 G38 G39 G40 G41 G42 G43 G44
G45 G46 G47 G48 G49 G50 G51 G52 G53 G54 G55 G56 G57 G58 G59
G60 G61 G62 G63 G64 G65 G66 G67 G68 G69 G70 G71 G72 G73 G74
G75 G76 G77 G78 G79 G80 G81 G82 G83 G84 G85 G86 G87 G88 G89
G90 G91 G92 G93 G94 G95 G96 G97 G98 G99 G100 G101 G102 G103 G104
G105 G106 G107 G108 G109 G110 G111 G112 G113 G114 G115 G116 G117 G118 G119
G120 G121 G122 G123 G124 G125 G126 G127 G128 G129 G130 G131 G132 G133 G134
G135 G136 G137 G138 G139 G140 G141 G142 G143 G144 G145 G146 G147 G148 G149
G150 G151 G152 G153 G154 G155 G156 G157 G158 G159 G160 G161 G162 G163 G164
G165 G166 G167 G168 G169 G170 G171 G172 G173 G174 G175 G176 G177 G178 G179
G180 G181 G182 G183 G184 G185 G186 G187 G188 G189 G190 G191 G192 G193 G194
G195 G196 G197 G198

**Supplementary Figure 22. Overview of all segmented Xenium glands used for curvature analysis (n = 194).**

Individual gland contours from the Xenium prostate dataset are shown separately and arranged in a grid for visualization. Boundary segments are colored by local curvature, with red indicating outward curvature and blue indicating inward curvature. This overview highlights the broad heterogeneity in gland size, shape, and boundary folding across the Xenium gland set, including both benign and GG1 glands used in the downstream morphomechanical analyses.

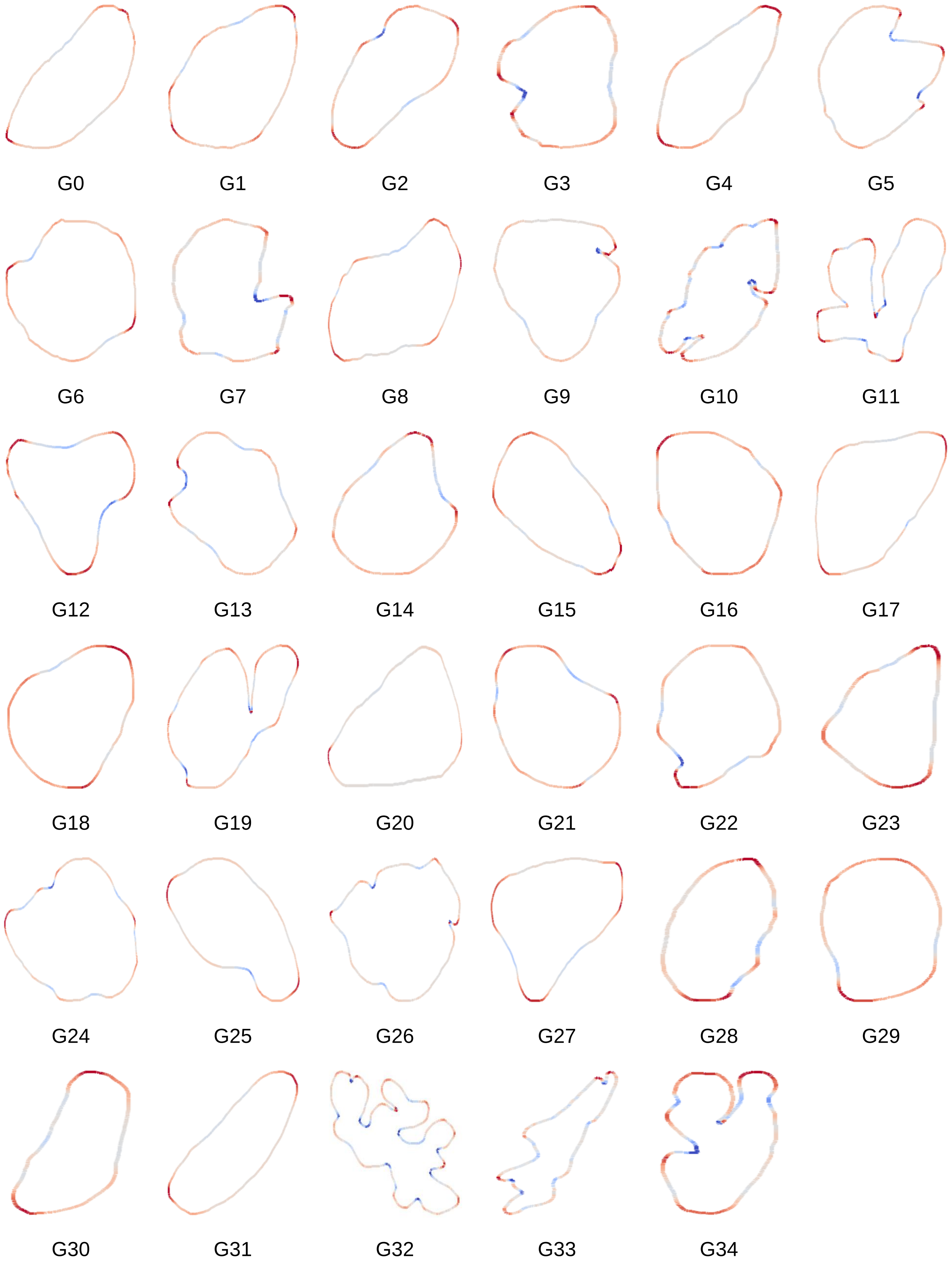
G0
G1
G2
G3
G4
G5
G6
G7
G8
G9
G10
G11
G12
G13
G14
G15
G16
G17
G18
G19
G20
G21
G22
G23
G24
G25
G26
G27
G28
G29
G30
G31
G32
G33
G34

**Supplementary Figure 23. Overview of all segmented benign Visium glands used for curvature analysis (n = 35).**

Individual benign gland contours from the Visium prostate dataset are shown separately and arranged in a grid for visualization. Boundary segments are colored by local curvature, with red indicating outward curvature and blue indicating inward curvature. This overview illustrates the diversity in gland size, shape, and boundary folding across the benign Visium gland set used for downstream morphomechanical analyses.

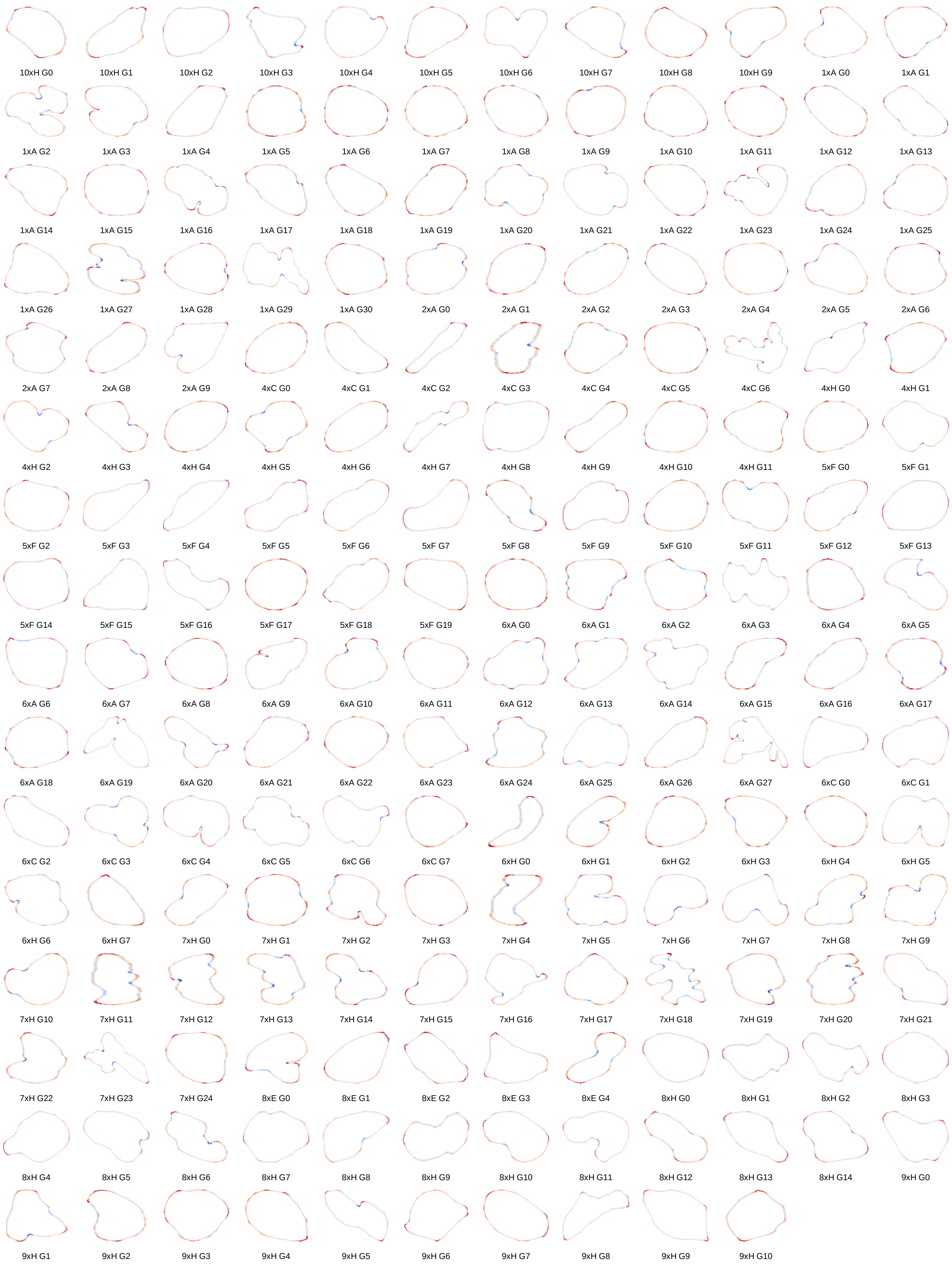
10xH G0
10xH G1
10xH G2
10xH G3
10xH G4
10xH G5
10xH G6
10xH G7
10xH G8
10xH G9
1xA G0
1xA G1
1xA G2
1xA G3
1xA G4
1xA G5
1xA G6
1xA G7
1xA G8
1xA G9
1xA G10
1xA G11
1xA G12
1xA G13
1xA G14
1xA G15
1xA G16
1xA G17
1xA G18
1xA G19
1xA G20
1xA G21
1xA G22
1xA G23
1xA G24
1xA G25
1xA G26
1xA G27
1xA G28
1xA G29
1xA G30
2xA G0
2xA G1
2xA G2
2xA G3
2xA G4
2xA G5
2xA G6
2xA G7
2xA G8
2xA G9
4xC G0
4xC G1
4xC G2
4xC G3
4xC G4
4xC G5
4xC G6
4xH G0
4xH G1
4xH G2
4xH G3
4xH G4
4xH G5
4xH G6
4xH G7
4xH G8
4xH G9
4xH G10
4xH G11
5xF G0
5xF G1
5xF G2
5xF G3
5xF G4
5xF G5
5xF G6
5xF G7
5xF G8
5xF G9
5xF G10
5xF G11
5xF G12
5xF G13
5xF G14
5xF G15
5xF G16
5xF G17
5xF G18
5xF G19
6xA G0
6xA G1
6xA G2
6xA G3
6xA G4
6xA G5
6xA G6
6xA G7
6xA G8
6xA G9
6xA G10
6xA G11
6xA G12
6xA G13
6xA G14
6xA G15
6xA G16
6xA G17
6xA G18
6xA G19
6xA G20
6xA G21
6xA G22
6xA G23
6xA G24
6xA G25
6xA G26
6xA G27
6xC G0
6xC G1
6xC G2
6xC G3
6xC G4
6xC G5
6xC G6
6xC G7
6xH G0
6xH G1
6xH G2
6xH G3
6xH G4
6xH G5
6xH G6
6xH G7
7xH G0
7xH G1
7xH G2
7xH G3
7xH G4
7xH G5
7xH G6
7xH G7
7xH G8
7xH G9
7xH G10
7xH G11
7xH G12
7xH G13
7xH G14
7xH G15
7xH G16
7xH G17
7xH G18
7xH G19
7xH G20
7xH G21
7xH G22
7xH G23
7xH G24
8xE G0
8xE G1
8xE G2
8xE G3
8xE G4
8xH G0
8xH G1
8xH G2
8xH G3
8xH G4
8xH G5
8xH G6
8xH G7
8xH G8
8xH G9
8xH G10
8xH G11
8xH G12
8xH G13
8xH G14
9xH G0
9xH G1
9xH G2
9xH G3
9xH G4
9xH G5
9xH G6
9xH G7
9xH G8
9xH G9
9xH G10

**Supplementary Figure 24. Overview of all segmented glands from spatial proteomics dataset 3 used for curvature analysis (n=160).**

Individual gland contours from spatial proteomics dataset 3 are shown separately and arranged in a grid for visualization. Boundary segments are colored by local curvature, with red indicating outward curvature and blue indicating inward curvature. This overview illustrates the heterogeneity in gland size, shape, and boundary folding across the dataset 3 gland set.

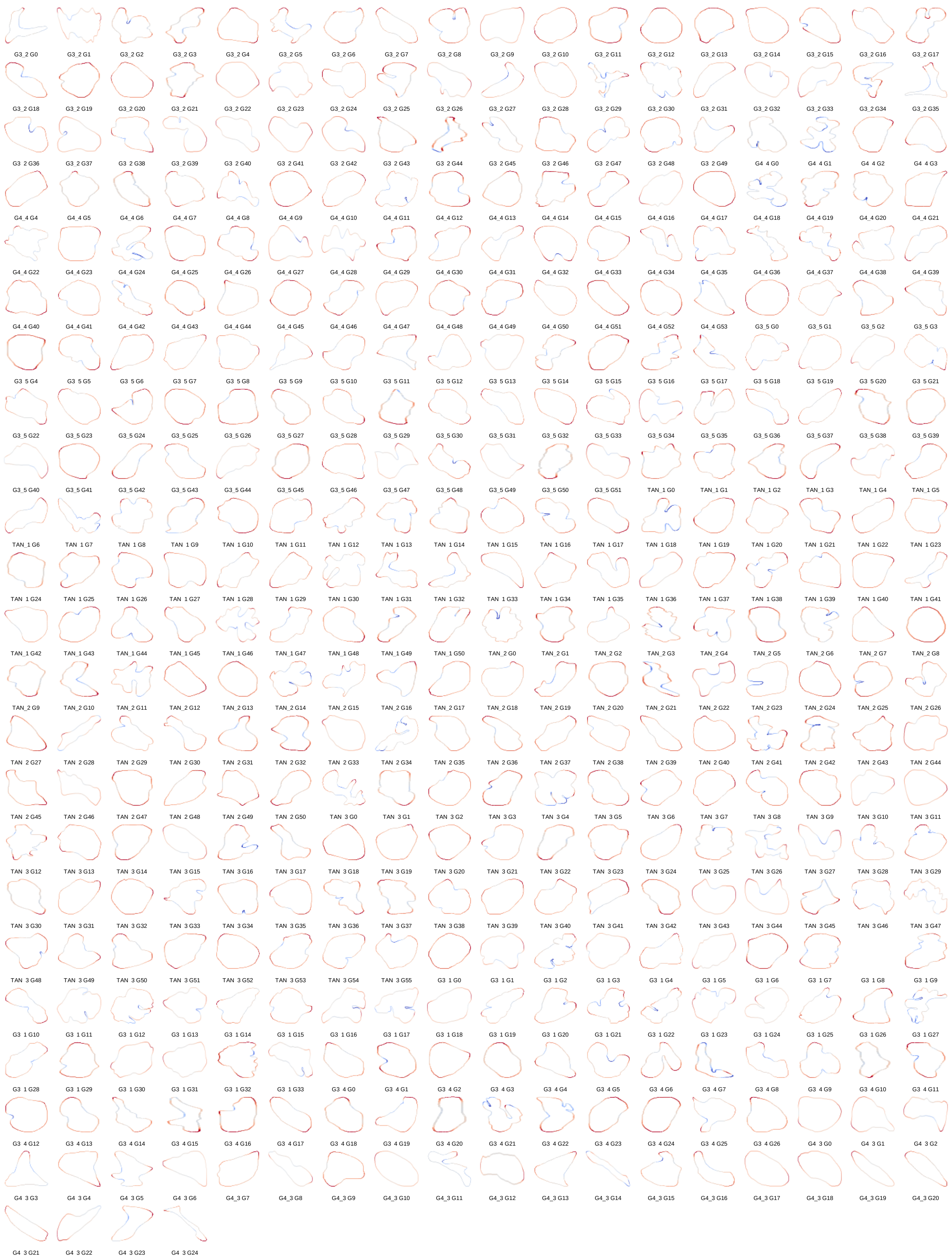
G3_2 G0
G3_2 G1
G3_2 G2
G3_2 G3
G3_2 G4
G3_2 G5
G3_2 G6
G3_2 G7
G3_2 G8
G3_2 G9
G3_2 G10
G3_2 G11
G3_2 G12
G3_2 G13
G3_2 G14
G3_2 G15
G3_2 G16
G3_2 G17
G3_2 G18
G3_2 G19
G3_2 G20
G3_2 G21
G3_2 G22
G3_2 G23
G3_2 G24
G3_2 G25
G3_2 G26
G3_2 G27
G3_2 G28
G3_2 G29
G3_2 G30
G3_2 G31
G3_2 G32
G3_2 G33
G3_2 G34
G3_2 G35
G3_2 G36
G3_2 G37
G3_2 G38
G3_2 G39
G3_2 G40
G3_2 G41
G3_2 G42
G3_2 G43
G3_2 G44
G3_2 G45
G3_2 G46
G3_2 G47
G3_2 G48
G3_2 G49
G4_4 G0
G4_4 G1
G4_4 G2
G4_4 G3
G4_4 G4
G4_4 G5
G4_4 G6
G4_4 G7
G4_4 G8
G4_4 G9
G4_4 G10
G4_4 G11
G4_4 G12
G4_4 G13
G4_4 G14
G4_4 G15
G4_4 G16
G4_4 G17
G4_4 G18
G4_4 G19
G4_4 G20
G4_4 G21
G4_4 G22
G4_4 G23
G4_4 G24
G4_4 G25
G4_4 G26
G4_4 G27
G4_4 G28
G4_4 G29
G4_4 G30
G4_4 G31
G4_4 G32
G4_4 G33
G4_4 G34
G4_4 G35
G4_4 G36
G4_4 G37
G4_4 G38
G4_4 G39
G4_4 G40
G4_4 G41
G4_4 G42
G4_4 G43
G4_4 G44
G4_4 G45
G4_4 G46
G4_4 G47
G4_4 G48
G4_4 G49
G4_4 G50
G4_4 G51
G4_4 G52
G4_4 G53
G3_5 G0
G3_5 G1
G3_5 G2
G3_5 G3
G3_5 G4
G3_5 G5
G3_5 G6
G3_5 G7
G3_5 G8
G3_5 G9
G3_5 G10
G3_5 G11
G3_5 G12
G3_5 G13
G3_5 G14
G3_5 G15
G3_5 G16
G3_5 G17
G3_5 G18
G3_5 G19
G3_5 G20
G3_5 G21
G3_5 G22
G3_5 G23
G3_5 G24
G3_5 G25
G3_5 G26
G3_5 G27
G3_5 G28
G3_5 G29
G3_5 G30
G3_5 G31
G3_5 G32
G3_5 G33
G3_5 G34
G3_5 G35
G3_5 G36
G3_5 G37
G3_5 G38
G3_5 G39
G3_5 G40
G3_5 G41
G3_5 G42
G3_5 G43
G3_5 G44
G3_5 G45
G3_5 G46
G3_5 G47
G3_5 G48
G3_5 G49
G3_5 G50
G3_5 G51
TAN_1 G0
TAN_1 G1
TAN_1 G2
TAN_1 G3
TAN_1 G4
TAN_1 G5
TAN_1 G6
TAN_1 G7
TAN_1 G8
TAN_1 G9
TAN_1 G10
TAN_1 G11
TAN_1 G12
TAN_1 G13
TAN_1 G14
TAN_1 G15
TAN_1 G16
TAN_1 G17
TAN_1 G18
TAN_1 G19
TAN_1 G20
TAN_1 G21
TAN_1 G22
TAN_1 G23
TAN_1 G24
TAN_1 G25
TAN_1 G26
TAN_1 G27
TAN_1 G28
TAN_1 G29
TAN_1 G30
TAN_1 G31
TAN_1 G32
TAN_1 G33
TAN_1 G34
TAN_1 G35
TAN_1 G36
TAN_1 G37
TAN_1 G38
TAN_1 G39
TAN_1 G40
TAN_1 G41
TAN_1 G42
TAN_1 G43
TAN_1 G44
TAN_1 G45
TAN_1 G46
TAN_1 G47
TAN_1 G48
TAN_1 G49
TAN_1 G50
TAN_2 G0
TAN_2 G1
TAN_2 G2
TAN_2 G3
TAN_2 G4
TAN_2 G5
TAN_2 G6
TAN_2 G7
TAN_2 G8
TAN_2 G9
TAN_2 G10
TAN_2 G11
TAN_2 G12
TAN_2 G13
TAN_2 G14
TAN_2 G15
TAN_2 G16
TAN_2 G17
TAN_2 G18
TAN_2 G19
TAN_2 G20
TAN_2 G21
TAN_2 G22
TAN_2 G23
TAN_2 G24
TAN_2 G25
TAN_2 G26
TAN_2 G27
TAN_2 G28
TAN_2 G29
TAN_2 G30
TAN_2 G31
TAN_2 G32
TAN_2 G33
TAN_2 G34
TAN_2 G35
TAN_2 G36
TAN_2 G37
TAN_2 G38
TAN_2 G39
TAN_2 G40
TAN_2 G41
TAN_2 G42
TAN_2 G43
TAN_2 G44
TAN_2 G45
TAN_2 G46
TAN_2 G47
TAN_2 G48
TAN_2 G49
TAN_2 G50
TAN_3 G0
TAN_3 G1
TAN_3 G2
TAN_3 G3
TAN_3 G4
TAN_3 G5
TAN_3 G6
TAN_3 G7
TAN_3 G8
TAN_3 G9
TAN_3 G10
TAN_3 G11
TAN_3 G12
TAN_3 G13
TAN_3 G14
TAN_3 G15
TAN_3 G16
TAN_3 G17
TAN_3 G18
TAN_3 G19
TAN_3 G20
TAN_3 G21
TAN_3 G22
TAN_3 G23
TAN_3 G24
TAN_3 G25
TAN_3 G26
TAN_3 G27
TAN_3 G28
TAN_3 G29
TAN_3 G30
TAN_3 G31
TAN_3 G32
TAN_3 G33
TAN_3 G34
TAN_3 G35
TAN_3 G36
TAN_3 G37
TAN_3 G38
TAN_3 G39
TAN_3 G40
TAN_3 G41
TAN_3 G42
TAN_3 G43
TAN_3 G44
TAN_3 G45
TAN_3 G46
TAN_3 G47
TAN_3 G48
TAN_3 G49
TAN_3 G50
TAN_3 G51
TAN_3 G52
TAN_3 G53
TAN_3 G54
TAN_3 G55
G3_1 G0
G3_1 G1
G3_1 G2
G3_1 G3
G3_1 G4
G3_1 G5
G3_1 G6
G3_1 G7
G3_1 G8
G3_1 G9
G3_1 G10
G3_1 G11
G3_1 G12
G3_1 G13
G3_1 G14
G3_1 G15
G3_1 G16
G3_1 G17
G3_1 G18
G3_1 G19
G3_1 G20
G3_1 G21
G3_1 G22
G3_1 G23
G3_1 G24
G3_1 G25
G3_1 G26
G3_1 G27
G3_1 G28
G3_1 G29
G3_1 G30
G3_1 G31
G3_1 G32
G3_1 G33
G3_4 G0
G3_4 G1
G3_4 G2
G3_4 G3
G3_4 G4
G3_4 G5
G3_4 G6
G3_4 G7
G3_4 G8
G3_4 G9
G3_4 G10
G3_4 G11
G3_4 G12
G3_4 G13
G3_4 G14
G3_4 G15
G3_4 G16
G3_4 G17
G3_4 G18
G3_4 G19
G3_4 G20
G3_4 G21
G3_4 G22
G3_4 G23
G3_4 G24
G3_4 G25
G3_4 G26
G4_3 G0
G4_3 G1
G4_3 G2
G4_3 G3
G4_3 G4
G4_3 G5
G4_3 G6
G4_3 G7
G4_3 G8
G4_3 G9
G4_3 G10
G4_3 G11
G4_3 G12
G4_3 G13
G4_3 G14
G4_3 G15
G4_3 G16
G4_3 G17
G4_3 G18
G4_3 G19
G4_3 G20
G4_3 G21
G4_3 G22
G4_3 G23
G4_3 G24

**Supplementary Figure 25. Overview of all segmented glands from spatial proteomics dataset 4 used for curvature analysis (n=199).**

Individual gland contours from spatial proteomics dataset 4 are shown separately and arranged in a grid for visualization. Boundary segments are colored by local curvature, with red indicating outward curvature and blue indicating inward curvature.

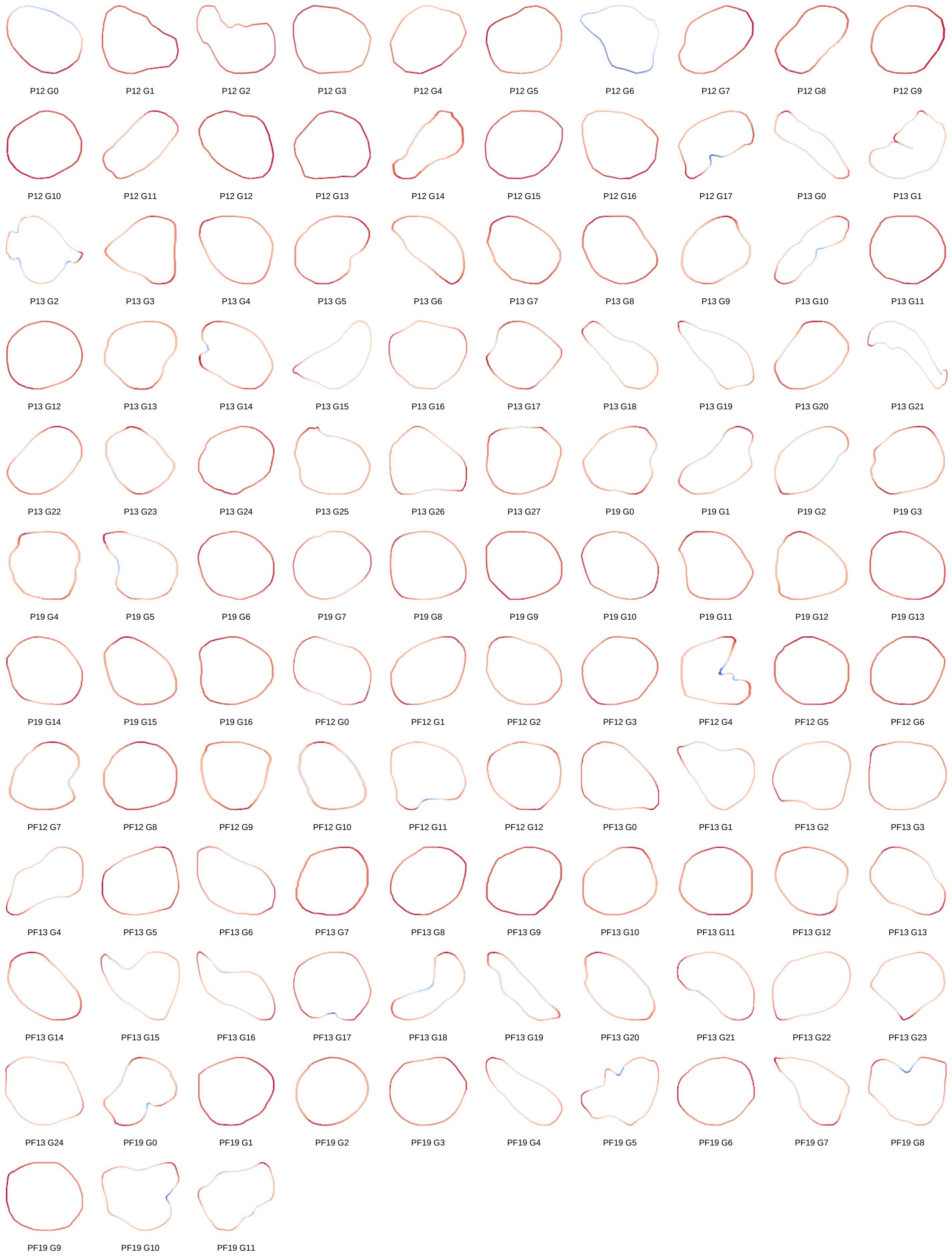
P12 G0
P12 G1
P12 G2
P12 G3
P12 G4
P12 G5
P12 G6
P12 G7
P12 G8
P12 G9
P12 G10
P12 G11
P12 G12
P12 G13
P12 G14
P12 G15
P12 G16
P12 G17
P13 G0
P13 G1
P13 G2
P13 G3
P13 G4
P13 G5
P13 G6
P13 G7
P13 G8
P13 G9
P13 G10
P13 G11
P13 G12
P13 G13
P13 G14
P13 G15
P13 G16
P13 G17
P13 G18
P13 G19
P13 G20
P13 G21
P13 G22
P13 G23
P13 G24
P13 G25
P13 G26
P13 G27
P19 G0
P19 G1
P19 G2
P19 G3
P19 G4
P19 G5
P19 G6
P19 G7
P19 G8
P19 G9
P19 G10
P19 G11
P19 G12
P19 G13
P19 G14
P19 G15
P19 G16
PF12 G0
PF12 G1
PF12 G2
PF12 G3
PF12 G4
PF12 G5
PF12 G6
PF12 G7
PF12 G8
PF12 G9
PF12 G10
PF12 G11
PF12 G12
PF13 G0
PF13 G1
PF13 G2
PF13 G3
PF13 G4
PF13 G5
PF13 G6
PF13 G7
PF13 G8
PF13 G9
PF13 G10
PF13 G11
PF13 G12
PF13 G13
PF13 G14
PF13 G15
PF13 G16
PF13 G17
PF13 G18
PF13 G19
PF13 G20
PF13 G21
PF13 G22
PF13 G23
PF13 G24
PF19 G0
PF19 G1
PF19 G2
PF19 G3
PF19 G4
PF19 G5
PF19 G6
PF19 G7
PF19 G8
PF19 G9
PF19 G10
PF19 G11

**Supplementary Figure 26. Overview of all segmented glands from spatial proteomics dataset 5 used for curvature analysis (n=96).**

Individual gland contours from spatial proteomics dataset 5 are shown separately and arranged in a grid for visualization. Boundary segments are colored by local curvature, with red indicating outward curvature and blue indicating inward curvature.

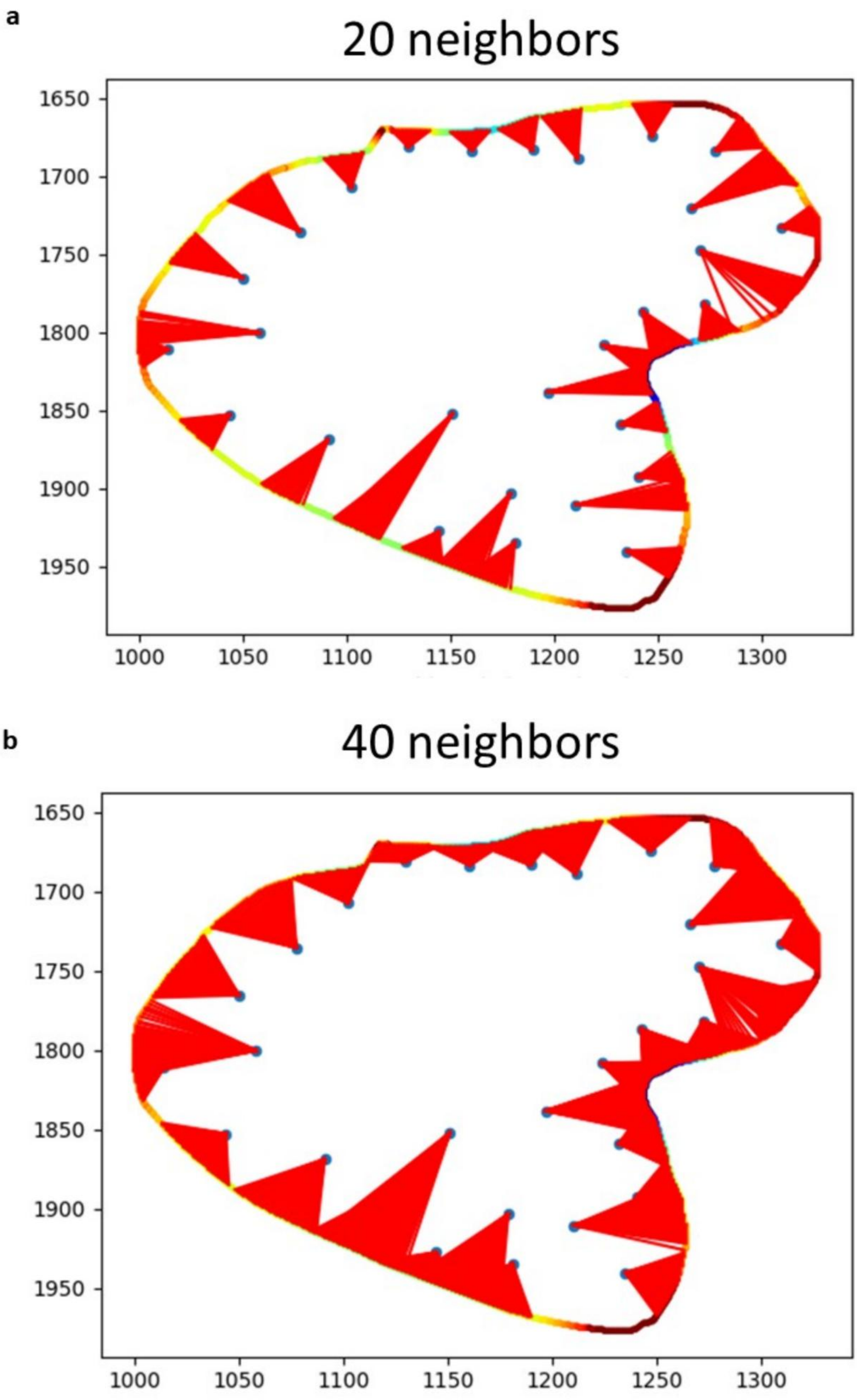
a
20 neighbors
1650
1700
1750
1800
1850
1900
1950
1000
1050
1100
1150
1200
1250
1300
b
40 neighbors
1650
1700
1750
1800
1850
1900
1950
1000
1050
1100
1150
1200
1250
1300

**Supplementary Figure 27. Effect of neighborhood size on curvature-based clustering.**
(a) Gland segmentation showing connections between cells when using 20 nearest neighbors.
(b) The same gland with 40 nearest neighbors. Increasing the number of neighbors results in broader connections across the gland boundary, smoothing local clustering patterns, and reducing sensitivity to small-scale variations.

**a**

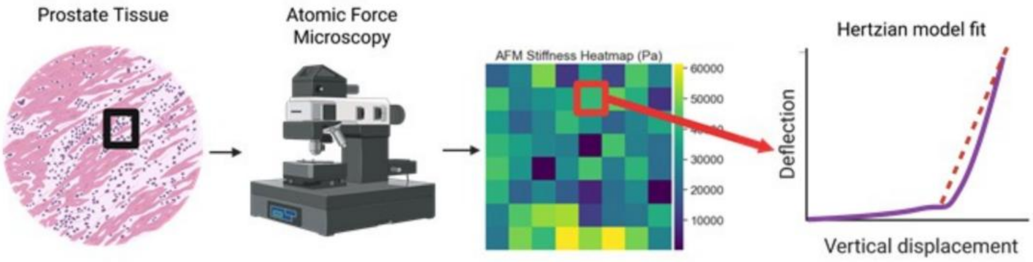


**b**

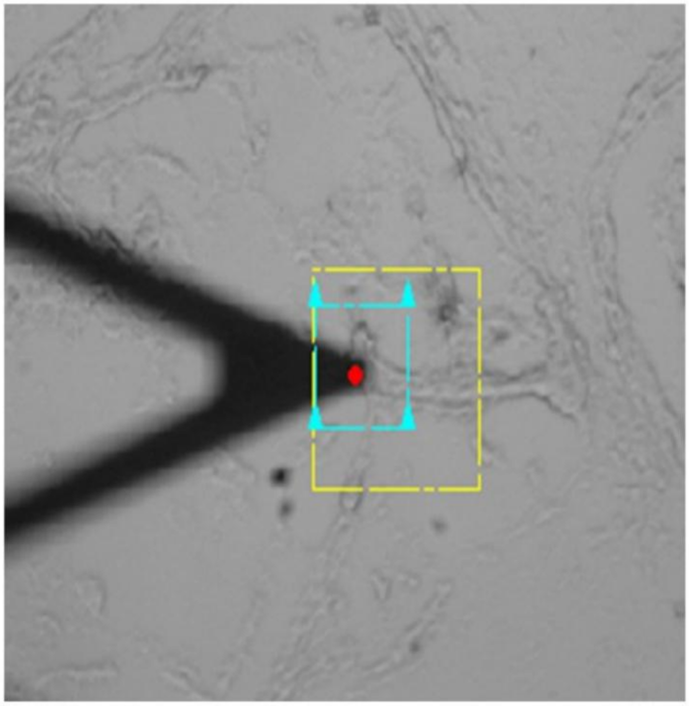

**c**

$$F = \frac{4}{3} E^* \sqrt{R}\, \delta^{3/2}$$

$F$: applied force (N)

$E^*$: reduced modulus (Pa)

$R$: tip radius (m)

$\delta$: indentation depth (m)

**d**

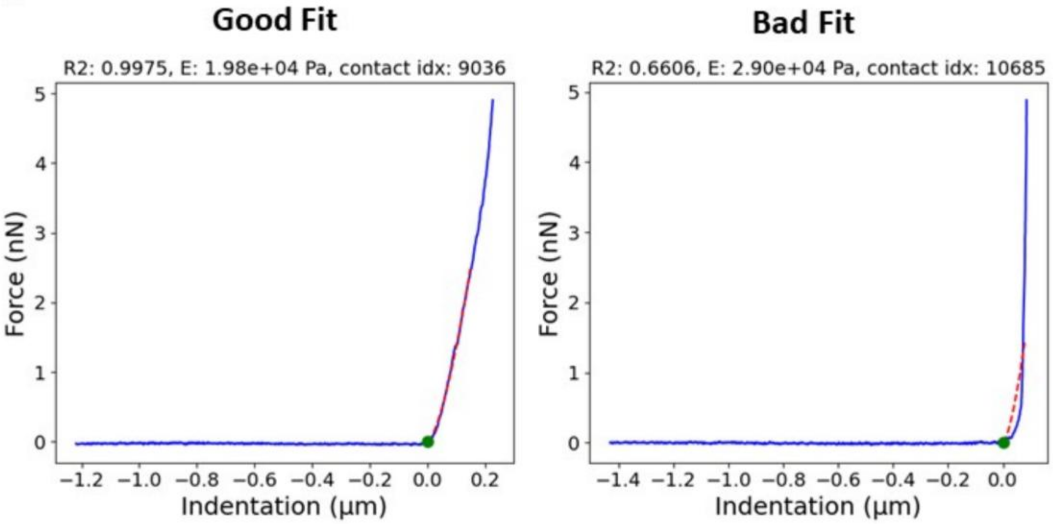

**Supplementary Figure 28: Measurement of tissue stiffness using Atomic Force Microscopy**

(a) Schematic overview of the workflow. Prostate tissue sections were selected and indented using AFM to obtain force–displacement curves. The data were fitted using the Hertzian contact model to calculate the reduced elastic modulus ($E*$).

(b) Optical image of the AFM probe in contact with the tissue surface. The yellow box indicates the scanning region, and the red dot marks the indentation site.

(c) The Hertzian model equation used for stiffness estimation, where F is the applied force, $E*$ is the reduced modulus, R is the tip radius, and δ is the indentation depth.

(d) Representative force–indentation curves from two measurement sites showing model fits (dashed red lines) and corresponding stiffness values. Some curves were kept (left), and some curves were discarded (right) because the R2 values were too low, or locating the contact point on the deflection curves was too ambiguous.

**a**

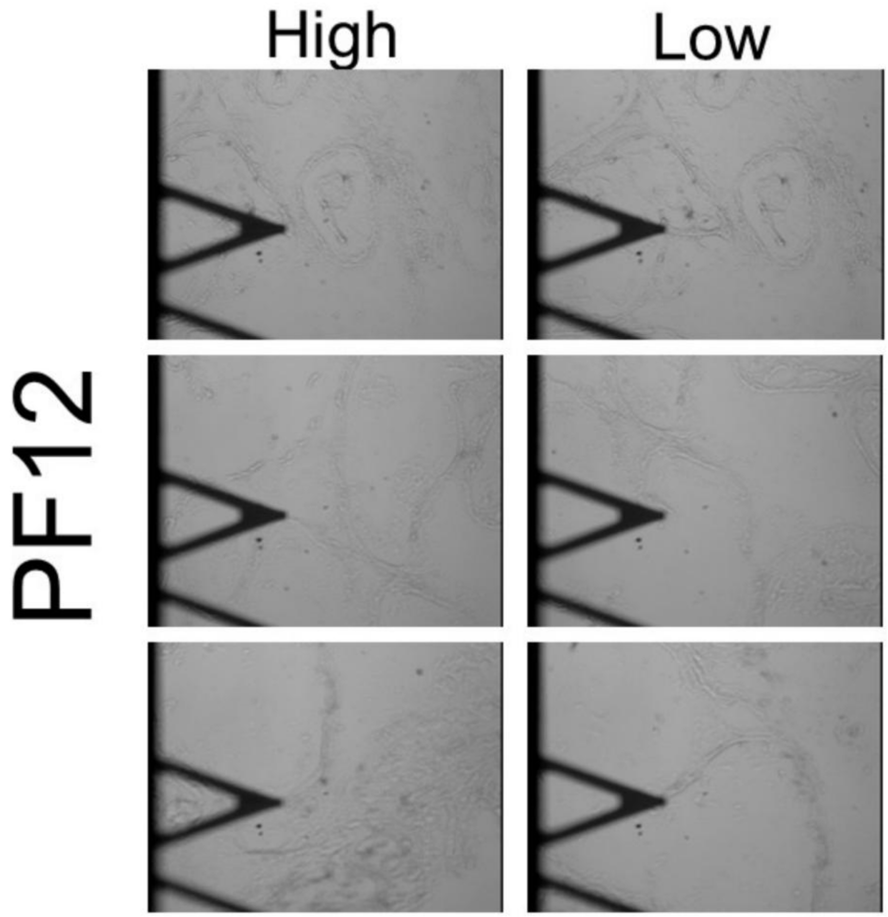


**b**

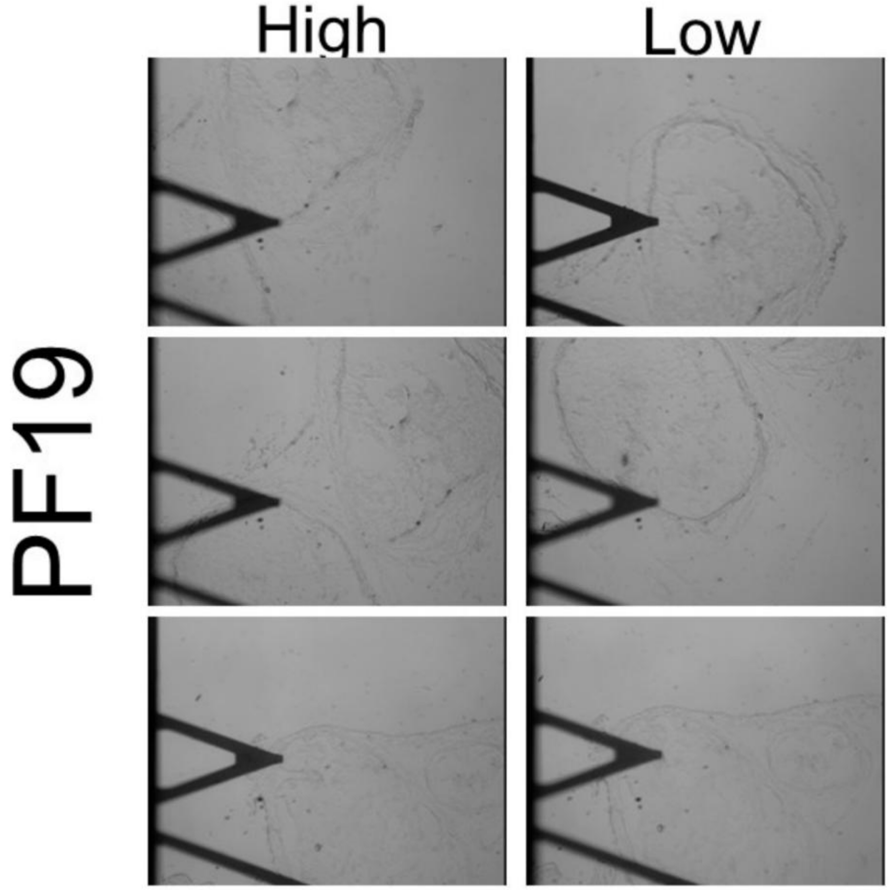

**Supplementary Figure 29. AFM measurement locations along gland borders in fixed FOXA1 control and knockout tissues.**

(a) Corresponding AFM images for FOXA1-knockout tissue (PF12). The AFM probe was positioned on or near the gland boundary for each measurement to capture local stiffness variations between high- and low-curvature regions.

(b) Representative brightfield AFM images showing indentation points along regions of high and low curvature in PF19.

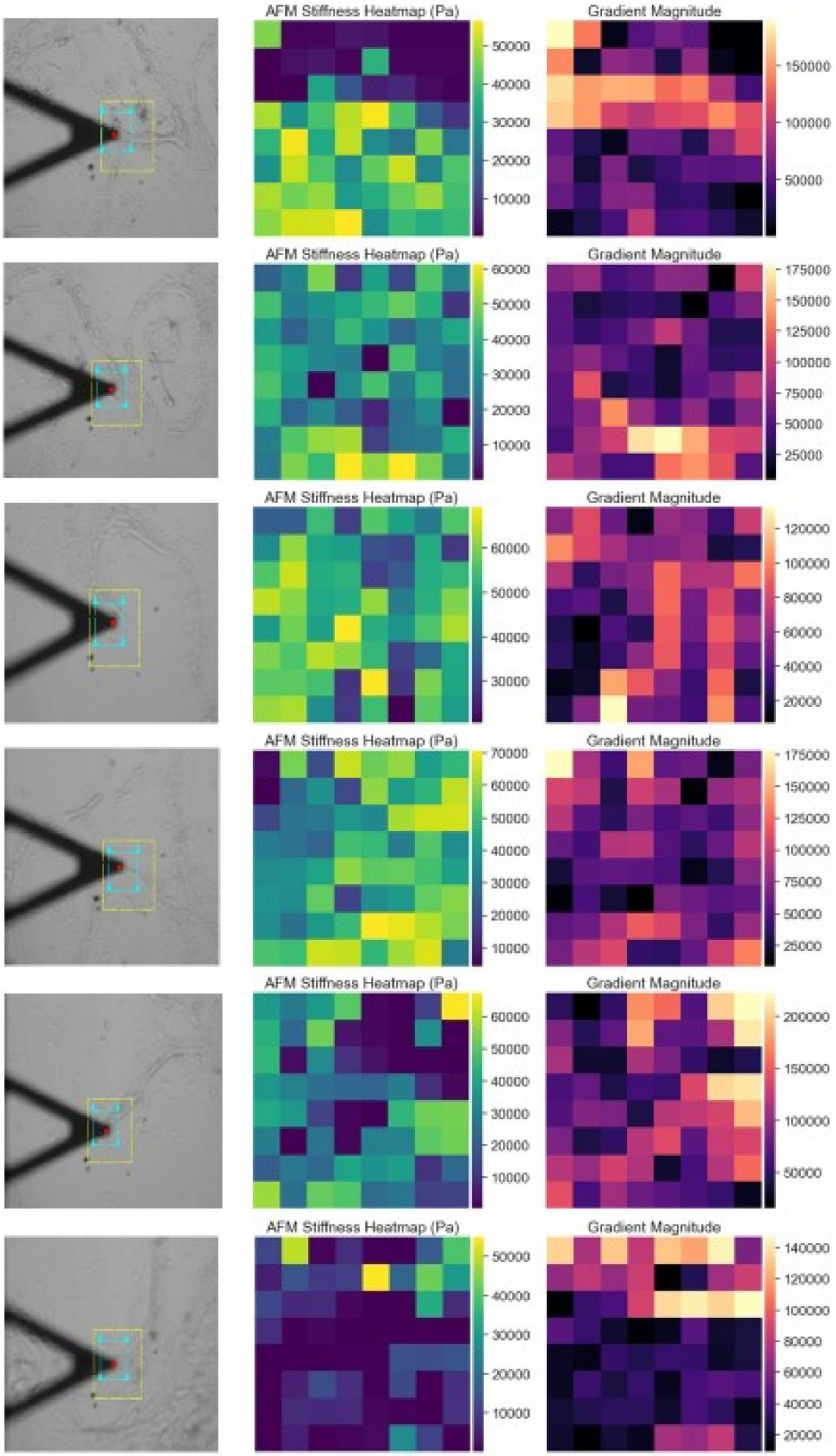

AFM Stiffness Heatmap (Pa)
50000
40000
30000
20000
10000
Gradient Magnitude
150000
100000
50000
AFM Stiffness Heatmap (Pa)
60000
50000
40000
30000
20000
10000
Gradient Magnitude
175000
150000
125000
100000
75000
50000
25000
AFM Stiffness Heatmap (Pa)
60000
50000
40000
30000
Gradient Magnitude
120000
100000
80000
60000
40000
20000
AFM Stiffness Heatmap (Pa)
70000
60000
50000
40000
30000
20000
10000
Gradient Magnitude
175000
150000
125000
100000
75000
50000
25000
AFM Stiffness Heatmap (Pa)
60000
50000
40000
30000
20000
10000
Gradient Magnitude
200000
150000
100000
50000
AFM Stiffness Heatmap (Pa)
50000
40000
30000
20000
10000
Gradient Magnitude
140000
120000
100000
80000
60000
40000
20000

**Supplementary Figure 30. AFM stiffness heatmaps and curve-fitting optimization for Hertzian model analysis.**

Representative AFM-derived stiffness heatmaps (middle) and corresponding gradient magnitude maps(right) from fixed prostate tissue regions imaged near the gland border (left). Each heatmap represents the raw stiffness distribution obtained from AFM scans, with the cyan box indicating the full scanned area. For quantitative analysis, we extracted a smaller 3×3 subregion centered on the AFM tip position to focus on the precise area of contact. Force–indentation curves within this localized region were manually examined, and contact points were refined to optimize Hertzian model fitting, yielding improved $R^2$ values and more reliable stiffness estimations.

a

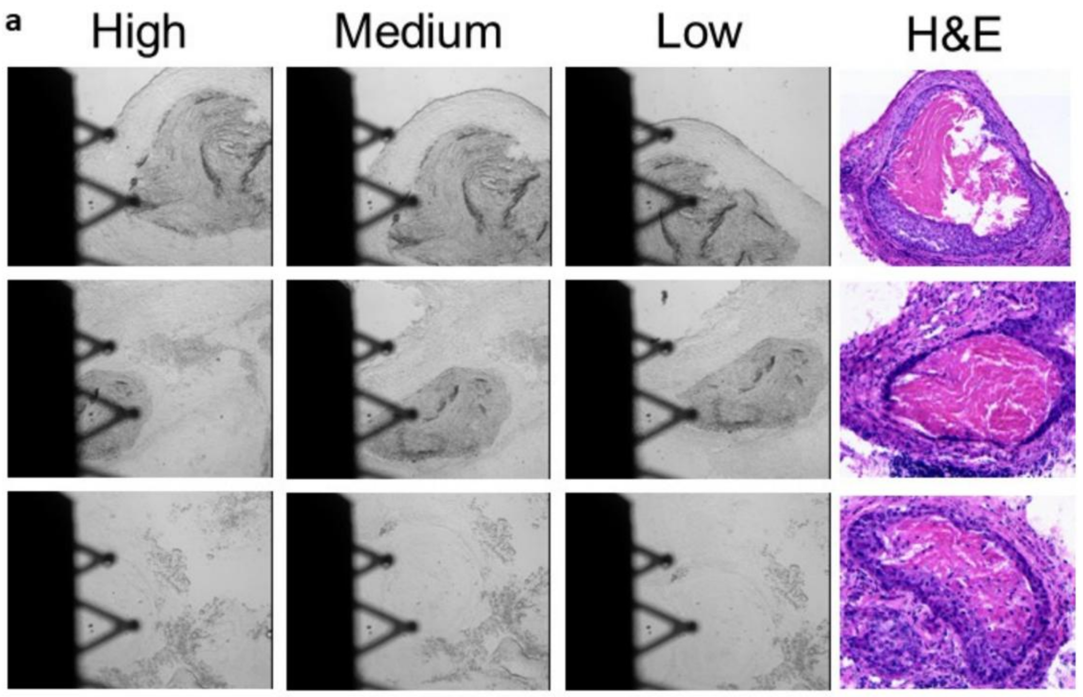


b

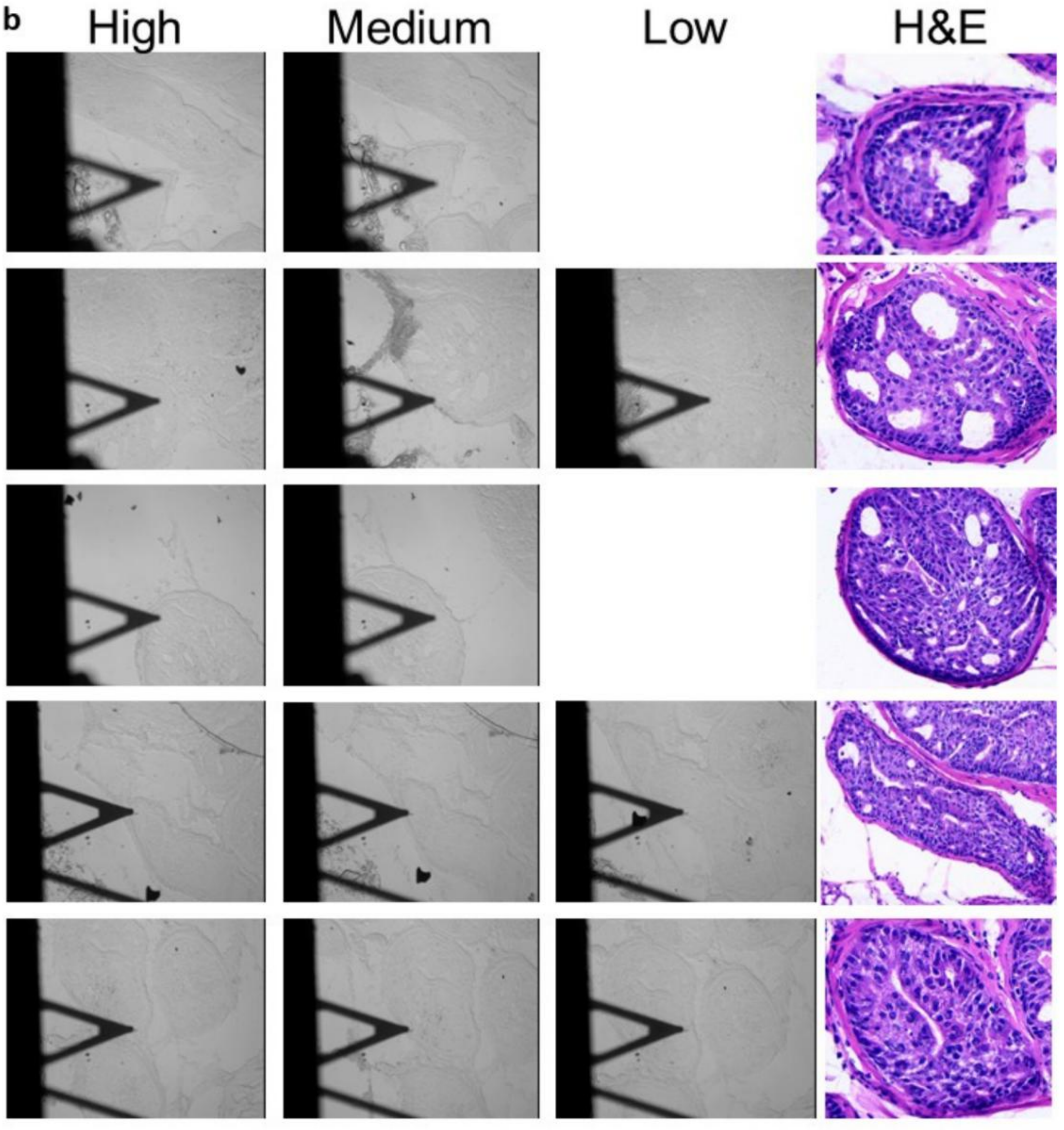

**Supplementary Figure 31. AFM measurement locations along gland borders in fresh FOXA1 control and knockout tissues.**

(a) Representative brightfield AFM images from fresh FOXA1-knockout tissue (PF12) showing indentation sites at regions of high, medium, and low curvature.

(b) Corresponding AFM images from fresh FOXA1-positive control tissue (P12) acquired under the same conditions. The AFM probe was positioned on or near the gland boundary for each indentation to measure local stiffness variations across curvature levels.

**a**

| Cycle | Marker | Species | Channel | Primary Dilution | Secondary dilution |
|---|---|---|---|---|---|
| Cycle 1 | Hoechst | N/A | Ch1 | 1:2000 | N/A |
| | AMACR/ P504S (P) | Rb | Ch2 | 1:100 | 1:1000 |
| | N/A | N/A | Ch3 | N/A | N/A |
| | P63 | M | Ch4 | 1:100 | 1:1000 |
| Cycle 2 | Hoechst | N/A | Ch1 | 1:2000 | N/A |
| | PSA | Rb | Ch2 | 1:100 | 1:1000 |
| | N/A | N/A | Ch3 | N/A | N/A |
| | CK HMW | M | Ch4 | 1:100 | 1:1000 |
| Cycle 5 | Hoechst | N/A | Ch1 | 1:2000 | N/A |
| | N/A | N/A | Ch2 | N/A | N/A |
| | H3k9Ac-555 | 555 | Ch3 | 1/250 dilution (4 μg/mL) | N/A |
| | H3k4me2-647 | 647 | Ch4 | 1/800 | N/A |
| Cycle 6 | Hoechst | N/A | Ch1 | 1:2000 | N/A |
| | H3k27me3-488 | 488 | Ch2 | 1/1600 | N/A |
| | N/A | N/A | Ch3 | N/A | N/A |
| | H4k12Ac-647 | 647 | Ch4 | 1/1600 | N/A |
| Cycle 7 | Hoechst | N/A | Ch1 | 1:2000 | N/A |
| | Concanavalin A | 488 | Ch2 | 1:20 | N/A |
| | Phalloidins | 555 | Ch3 | 1:40 | N/A |
| | WGA | 647 | Ch4 | 1:400 | N/A |

**b**

| Patient | Age | Sex | Organ/Anatomic Site | Pathology diagnosis | Grade | Stage | Type | Gleason Score | Gleason Grade |
|---|---|---|---|---|---|---|---|---|---|
| 1xA | 74 | M | Prostate | Hyperplasia of prostate tissue | - | - | Hype rplasi | - | - |
| 4xC | 82 | M | Prostate | Adenocarcinoma | 3 | III | Malig | 5+5 | 5 |
| 6xA | 34 | M | Prostate | Prostate tissue | - | - | Norm | - | - |
| 2xA | 74 | M | Prostate | Hyperplasia of prostate tissue | - | - | Hype rplasi | - | - |
| 6xH | 60 | M | Prostate | Adenocarcinoma | 1 | I | Malig | 2+3 | 2 |
| 9xH | 65 | M | Prostate | Adenocarcinoma | 2 | I | Malig | 3+3 | 3 |
| 4xH | 75 | M | Prostate | Adenocarcinoma | 1 | IIB | Malig | 2+2 | 2 |
| 5xF | 64 | M | Prostate | Adenocarcinoma | 2 | III | Malig | 3+4 | 3 |
| 6xC | 72 | M | Prostate | Adenocarcinoma | 3 | III | Malig | 5+5 | 5 |
| 7xH | 71 | M | Prostate | Adenocarcinoma | 2 | I | Malig | 3+2 | 3 |
| 8xE | 78 | M | Prostate | Adenocarcinoma | 3 | IV | Malig | 5+5 | 5 |
| 8xH | 71 | M | Prostate | Adenocarcinoma | 2 | IIA | Malig | 3+4 | 3-4 |
| 10xH | 66 | M | Prostate | Adenocarcinoma | 2 | I | Malig | 3+3 | 3 |

**Supplementary Table 1. Multiplex immunofluorescence markers and patient cohort information.**

(a) Antibody panel used for multiplexed immunofluorescence staining across cycles.

(b) Clinical and pathological characteristics of the patient cohort. Table includes patient ID, age, sex, organ site, pathology diagnosis, grade, stage, tissue type (hyperplasia, normal, malignant), Gleason score, and corresponding Gleason grade.

a

### Spatial Transcriptomics

#### Xenium Dataset

| Benign | GG1 |
|---|---|
| 52 | 142 |

#### Visium Dataset

| Benign | GG4 |
|---|---|
| 35 | 9 |

b

### Spatial Proteomics

##### Dataset 3

| Gleason Group | Number of Tissues | Total Number of Glands |
|---|---|---|
| Benign | 6 | 89 |
| GG1 | 1 | 12 |
| GG2 | 2 | 28 |
| GG3 | 4 | 31 |

##### Dataset 4

| Gleason Grade | Number of tissues | Total glands |
|---|---|---|
| G3 | 2 | 69 |
| G4 | 1 | 25 |
| TAN | 2 | 105 |

##### Dataset 5

| Patient Type | Number of tissues | Total glands |
|---|---|---|
| P | 3 | 50 |
| PF | 3 | 46 |

**Supplementary Table 2. Summary of spatial transcriptomics and spatial proteomics datasets used for gland-level morphomechanical analysis.**

(a)Spatial transcriptomics datasets included Xenium and Visium prostate tissue datasets. The Xenium dataset contained 52 benign glands and 142 GG1 glands, while the Visium dataset contained 35 benign glands and 9 GG4 glands. (b) Spatial proteomics datasets included multiplexed protein imaging cohorts across benign, Gleason grade group, tumor-adjacent normal, and FOXA1 control/knockout tissue groups. Dataset 3 included 6 benign tissues with 89 glands, 1 GG1 tissue with 12 glands, 2 GG2 tissues with 28 glands, and 4 GG3 tissues with 31 glands. Dataset 4 included 2 G3 tissues with 69 glands, 1 G4 tissue with 25 glands, and 2 TAN tissues with 105 glands. Dataset 5 included 3 control P tissues with 50 glands and 3 FOXA1-knockout PF tissues with 46 glands.
